## Supplementary data for "High-speed 3D DNA PAINT and unsupervised clustering for unlocking 3D DNA origami cryptography"

[a]Department of Physics, Arizona State University, Tempe, Arizona, USA.; [b]Center for Molecular Design and Biomimetics at the Biodesign Institute, Arizona State University, Tempe, Arizona, USA.; [c]School of Molecular Sciences, Arizona State University, Tempe, Arizona, USA.; [d]School of Computing and Augmented Intelligence, Arizona State University, Tempe, Arizona, USA.; [e]School of Electrical, Computer and Energy Engineering, Arizona State University, Tempe, Arizona, USA.

[1]To whom correspondence should be addressed. E-mail: {gwisna,rhariadi}@asu.edu

### S1. Methods

**Unsupervised classification as described by Huijben et al.[66]** We follow the protocol for unsupervised classification to classify a pre-labeled mixture of “NSF” and “ASU” datasets into several classes. The superparticles of each class are then fed into our template alignment method to read out the bit. We count the number of each label in each class to calculate the accuracy (Fig. S15).

**ResNet CNN supervised classification.** Picasso’s Simulate and Render modules were utilized to generate 75x75 images of each of the 26 letters from the alphabet. Images were then filtered using the root mean squared distance between each pixel and the center of mass of the image, totaling 41,096 images. The dataset was split into a training dataset of 22,749 images and a testing dataset of 18,347 images. Twenty percent of the training set was further split into a validation dataset used to select the final model.

**ResNet implementation.** Our ResNet implementation uses transfer learning on ResNet-50.[72] The final layer was replaced with a linear layer that gives 12 outputs with a sigmoid activation, one for each binding site in the encryption template of the “NSF” pattern. A threshold of 0.5 was used to distinguish between bits that are on and off, which are then decoded into the corresponding letter.

**Model training and evaluation.** The network was trained with the Pytorch implementation of the Adam optimizer at a learning rate of $10^{-3}$ for the linear layer and $10^{-4}$ for the other layers for 20 epochs. A cosine annealing learning rate scheduler and binary cross entropy loss were used. After training, the epoch with the lowest loss over the validation set was selected as the final model. This model then ran predictions over the testing dataset (Supplementary Fig. S16).

### S2. Supplementary Tables

| Name | Sequence | Note |
| --- | --- | --- |
| 0[47]1[31] | AGAAAGGAACAACTAAAGGAATTCAAAAAAA | Core staple |
| 1[96]3[95] | AAACAGCTTTTTGCGGGATCGTCAACACTAAA | Core staple |
| 2[111]0[112] | AAGGCCGCTGATACCGATAGTTGCGACGTTAG | Core staple |
| 3[160]4[144] | TTGACAGGCCACCACCAGAGCCGCGATTTGTA | Core staple |
| 5[160]6[144] | GCAAGGCCTCACCAGTAGCACCATGGGCTTGA | Core staple |
| 6[239]4[240] | GAAATTATTGCCTTTAGCGTCAGACCGGAACC | Core staple |
| 7[224]9[223] | AACGCAAAGATAGCCGAACAAACCCTGAAC | Core staple |
| 8[239]6[240] | AAGTAAGCAGACACCACGGAATAATATTGACG | Core staple |
| 9[224]11[223] | AAAGTCACAAAATAAACAGCCAGCGTTTTA | Core staple |
| 10[239]8[240] | GCCAGTTAGAGGGTAATTGAGCGCTTTAAGAA | Core staple |
| 11[224]13[223] | GCGAACCTCCAAGAACGGGTATGACAATAA | Core staple |
| 13[96]15[95] | TAGGTAAACTATTTTTGAGAGATCAAACGTTA | Core staple |
| 0[79]1[63] | ACAACTTTCAACAGTTTCAGCGGATGTATCGG | Core staple |
| 1[128]4[128] | TGACAACTCGCTGAGGCTTGCATTATACCAAGCGCGATGATAAA | Core staple |
| 2[143]1[159] | ATATTCGGAACCATCGCCCACGCAGAGAAGGA | Core staple |
| 3[224]5[223] | TTAAAGCCAGAGCCGCCACCCTCGACAGAA | Core staple |
| 5[224]7[223] | TCAAGTTTCATTAAAGGTGAATATAAAAGA | Core staple |
| 6[271]4[272] | ACCGATTGTCGGCATTTTCGGTCATAATCA | Core staple |
| 7[248]9[255] | GTTTATTTTGTCACAATCTTACCGAAGCCCTTTAATATCA | Core staple |
| 8[271]6[272] | AATAGCTATCAATAGAAAATTCAACATTCA | Core staple |
| 9[256]11[255] | GAGAGATAGAGCGTCTTTCCAGAGGTTTTGAA | Core staple |
| 10[271]8[272] | ACGCTAACACCCACAAGAATTGAAAATAGC | Core staple |
| 11[256]13[255] | GCCTTAAACCAATCAATAATCGGCACGCGCCT | Core staple |
| 13[128]15[127] | GAGACAGCTAGCTGATAAATTAATTTTTGT | Core staple |
| 0[111]1[95] | TAAATGAATTTTCTGTATGGGATTAATTTCTT | Core staple |
| 1[160]2[144] | TTAGGATTGGCTGAGACTCCTCAATAACCGAT | Core staple |
| 2[175]0[176] | TATTAAGAAGCGGGGTTTTGCTCGTAGCAT | Core staple |
| 4[79]2[80] | GCGCAGACAAGAGGCAAAAGAATCCCTCAG | Core staple |
| 6[47]4[48] | TACGTTAAAGTAATCTTGACAAGAACCGAACT | Core staple |
| 7[32]9[31] | TTTAGGACAAATGCTTTAAACAATCAGGTC | Core staple |
| 8[47]6[48] | ATCCCCCTATACCACATTCAACTAGAAAAATC | Core staple |
| 9[32]11[31] | TTTACCCCAACATGTTTTAAATTTCCATAT | Core staple |
| 10[47]8[48] | CTGTAGCTTGACTATTATAGTCAGTTCATTGA | Core staple |
| 11[32]13[31] | AACAGTTTTGTACCAAAAACATTTTATTTC | Core staple |
| 12[79]10[80] | AAATTAAGTTGACCATTAGATACTTTTGCG | Core staple |
| 13[160]14[144] | GTAATAAGTTAGGCAGAGGCATTTATGATATT | Core staple |
| 0[175]0[144] | TCCACAGACAGCCCTCATAGTTAGCGTAACGA | Core staple |
| 1[192]4[192] | GCGGATAACCTATTATTCTGAAACAGACGATTGGCCTTGAAGAGCCAC | Core staple |
| 2[207]0[208] | TTTCGGAAGTGCCGTCGAGAGGGTGAGTTTCG | Core staple |
| 4[143]3[159] | TCATCGCCAACAAAGTACAACGGACGCCAGCA | Core staple |
| 6[79]4[80] | TTATACCACCAAATCAACGTAACGAACGAG | Core staple |
| 7[56]9[63] | ATGCAGATACATAACGGGAATCGTCATAAATAAAGCAAAG | Core staple |
| 8[79]6[80] | AATACTGCCCAAAAGGAATTACGTGGCTCA | Core staple |
| 9[64]11[63] | CGGATTGCAGAGCTTAATTGCTGAAACGAGTA | Core staple |
| 10[79]8[80] | GATGGCTTATCAAAAAGATTAAGAGCGTCC | Core staple |
| 11[64]13[63] | GATTTAGTCAATAAAGCCTCAGAGAACCCTCA | Core staple |
| 12[143]11[159] | TTCTACTACGCGAGCTGAAAAGGTTACCGCGC | Core staple |
| 13[192]15[191] | GTAAAGTAATCGCCATATTTAACAAAACTTTT | Core staple |
| 0[239]1[223] | AGGAACCCATGTACCGTAACACTTGATATAA | Core staple |
| 1[224]3[223] | GTATAGCAAACAGTTAATGCCCAATCCTCA | Core staple |
| 2[239]0[240] | GCCCGTATCCGGAATAGGTGTATCAGCCCAAT | Core staple |
| 4[207]2[208] | CCACCCTCTATTCACAAACAAATACCTGCCTA | Core staple |
| 6[111]4[112] | ATTACCTTTGAATAAGGCTTGCCCAAATCCGC | Core staple |
| 7[96]9[95] | TAAGAGCAAATGTTTAGACTGGATAGGAAGCC | Core staple |
| 8[111]6[112] | AATAGTAAACACTATCATAACCCTCATTGTGA | Core staple |
| 9[96]11[95] | CGAAAGACTTTGATAAGAGGTCATATTTCGCA | Core staple |
| 10[111]8[112] | TTGCTCCTTTCAAATATCGCGTTTGAGGGGGT | Core staple |
| 11[96]13[95] | AATGGTCAACAGGCAAGGCAAAGAGTAATGTG | Core staple |
| 12[207]10[208] | GTACCGCAATTCTAAGAACGCGAGTATTATTT | Core staple |
| 13[224]15[223] | ACAACATGCCAACGCTCAACAGTCTTCTGA | Core staple |
| 0[271]1[255] | CCACCCTCATTTTCAGGGATAGCAACCGTACT | Core staple |
| 1[256]4[256] | CAGGAGGTGGGGTCAGTGCCTTGAGTCTCTGAATTTACCGGGAACCAG | Core staple |
| 2[271]0[272] | GTTTTAACTTAGTACCGCCACCCAGAGCCA | Core staple |
| 4[271]2[272] | AAATCACCTTCCAGTAAGCGTCAGTAATAA | Core staple |
| 6[143]5[159] | GATGGTTTGAACGAGTAGTAAATTTACCATTA | Core staple |
| 7[120]9[127] | CGTTTACCAGACGACAAAGAAGTTTTGCCATAATTCGA | Core staple |
| 8[143]7[159] | CTTTTGCAGATAAAAACCAAAATAAAGACTCC | Core staple |
| 9[128]11[127] | GCTTCAATCAGGATTAGAGAGTTATTTTCA | Core staple |
| 10[143]9[159] | CCAACAGGAGCGAACCAGACCGGAGCCTTTAC | Core staple |
| 11[128]13[127] | TTTGGGGATAGTAGTAGCATTAAAAGGCCG | Core staple |

| | | |
|---|---|---|
| 12[271]10[272] | TGTAGAAATCAAGATTAGTTGCTCTTACCA | Core staple |
| 13[256]15[255] | GTTTATCAATATGCGTTATACAAACCGACCGT | Core staple |
| 1[32]3[31] | AGGCTCCAGAGGCTTTGAGGACACGGGTAA | Core staple |
| 2[47]0[48] | ACGGCTACAAAAGGAGCCTTTAATGTGAGAAT | Core staple |
| 3[32]5[31] | AATACGTTTGAAAGAGGACAGACTGACCTT | Core staple |
| 5[32]7[31] | CATCAAGTAAAACGAACTAACGAGTTGAGA | Core staple |
| 6[175]4[176] | CAGCAAAAGGAAACGTCACCAATGAGCCGC | Core staple |
| 7[160]8[144] | TTATTACGAAGAACTGGCATGATTGCGAGAGG | Core staple |
| 8[175]6[176] | ATACCCAACAGTATGTTAGCAAATTAGAGC | Core staple |
| 9[160]10[144] | AGAGAGAAAAAAATGAAAATAGCAAGCAAACT | Core staple |
| 10[175]8[176] | TTAACGTCTAACATAAAAACAGGTAACGGA | Core staple |
| 11[160]12[144] | CCAATAGCTCATCGTAGGAATCATGGCATCAA | Core staple |
| 13[32]15[31] | AACGCAAAATCGATGAACGGTACCGGTTGA | Core staple |
| 14[47]12[48] | AACAAGAGGGATAAAAATTTTTAGCATAAAGC | Core staple |
| 1[64]4[64] | TTTATCAGGACAGCATCGGAACGACACCAACCTAAAACGAGGTCAATC | Core staple |
| 2[79]0[80] | CAGCGAAACTTGCTTTCGAGGTGTTGCTAA | Core staple |
| 3[96]5[95] | ACACTCATCCATGTTACTTAGCCGAAAGCTGC | Core staple |
| 5[96]7[95] | TCATTCAGATGCGATTTTAAGAACAGGCATAG | Core staple |
| 6[207]4[208] | TCACCGACGCACCGTAATCAGTAGCAGAACCG | Core staple |
| 7[184]9[191] | CGTAGAAAATACATACCGAGGAAACGCAATAAGAAGCGCA | Core staple |
| 8[207]6[208] | AAGGAAACATAAAGGTGGCAACATTATCACCG | Core staple |
| 9[192]11[191] | TTAGACGGCCAAATAAGAAACGATAGAAGGCT | Core staple |
| 10[207]8[208] | ATCCCAATGAGAATTAACTGAACAGTTACCAG | Core staple |
| 11[192]13[191] | TATCCGGTCTCATCGAGAACAAGCGACAAAAG | Core staple |
| 13[64]15[63] | TATATTTTGTCATTGCCTGAGAGTGGAAGATT | Core staple |
| 14[79]12[80] | GCTATCAGAAATGCAATGCCTGAATTAGCA | Core staple |
| 14[111]12[112] | GAGGGTAGGATTCAAAAGGGTGAGACATCCAA | Core staple |
| 15[96]17[95] | ATATTTTGGCTTTCATCAACATTATCCAGCCA | Core staple |
| 16[111]14[112] | TGTAGCCATTAAAATTCGCATTAAATGCCGGA | Core staple |
| 17[224]19[223] | CATAAATCTTTGAATACCAAGTGTTAGAAC | Core staple |
| 19[32]21[31] | GTCGACTTCGGCCAACGCGCGGGGTTTTTC | Core staple |
| 21[56]23[63] | AGCTGATTGCCCTTCAGAGTCCACTATTAAAGGGTGCCGT | Core staple |
| 22[79]20[80] | TGGAACAACCGCCTGGCCCTGAGGCCCGCT | Core staple |
| 23[96]22[112] | CCCGATTTAGAGCTTGACGGGGAAAAAGAATA | Core staple |
| 16[271]14[272] | CTTAGATTTAAGGCGTTAAATAAAGCCTGT | Core staple |
| 14[143]13[159] | CAACCGTTTCAAATCACCATCAATTCGAGCCA | Core staple |
| 15[128]18[128] | TAAATCAAAATAATTCGCGTCTCGGAAACCAGGCAAAGGGAAGG | Core staple |
| 16[143]15[159] | GCCATCAAGCTCATTTTTTAACCACAAATCCA | Core staple |
| 18[47]16[48] | CCAGGGTTGCCAGTTTGAGGGGACCCGTGGGA | Core staple |
| 19[96]21[95] | CTGTGTGATTGCGTTGCGCTCACTAGAGTTGC | Core staple |
| 21[96]23[95] | AGCAAGCGTAGGGTTGAGTGTTGTAGGGAGCC | Core staple |
| 22[111]20[112] | GCCCGAGAGTCCACGCTGGTTTGCAGCTAACT | Core staple |
| 23[128]23[159] | AACGTGGCGAGAAAGGAAGGGAAACCAGTAA | Core staple |
| 18[271]16[272] | CTTTTACAAAATCGTCGCTATTAGCGATAG | Core staple |
| 14[175]12[176] | CATGTAATAGAATATAAAGTACCAAGCCGT | Core staple |
| 15[160]16[144] | ATCGCAAGTATGTAAATGCTGATGATAGGAAC | Core staple |
| 16[175]14[176] | TATAACTAACAAAGAACGCGAGAACGCCAA | Core staple |
| 18[79]16[80] | GATGTGCTTCAGGAAGATCGCACAATGTGA | Core staple |
| 19[160]20[144] | GCAATTCACATATTCCTGATTATCAAAGTGTA | Core staple |
| 21[120]23[127] | CCCAGCAGGCGAAAAATCCCTTATAAATCAAGCCGGCG | Core staple |
| 22[143]21[159] | TCGGCAAATCCTGTTTGATGGTGGACCCTCAA | Core staple |
| 23[160]22[176] | TAAAAGGGACATTCTGGCCAACAAAGCATC | Core staple |
| 20[271]18[272] | CTCGTATTAGAAATTGCGTAGATACAGTAC | Core staple |
| 14[207]12[208] | AATTGAGAATTCTGTCCAGACGACTAAACCAA | Core staple |
| 15[192]18[192] | TCAAATATAACCTCCGGCTTAGGTAACAATTTCATTTGAAGGCGAATT | Core staple |
| 16[207]14[208] | ACCTTTTTATTTTAGTTAATTTCATAGGGCTT | Core staple |
| 18[111]16[112] | TCTTCGCTGCACCGCTTCTGGTGCGGCCTTCC | Core staple |
| 19[224]21[223] | CTACCATAGTTTGAGTAACATTTAAAATAT | Core staple |
| 21[160]22[144] | TCAATATCGAACCTCAAATATCAATTCCGAAA | Core staple |
| 22[175]20[176] | ACCTTGCTTGGTCAGTTGGCAAAGAGCGGA | Core staple |
| 23[192]22[208] | ACCCTTCTGACCTGAAAGCGTAAGACGCTGAG | Core staple |
| 22[271]20[272] | CAGAAGATTAGATAATACATTTGTCGACAA | Core staple |
| 14[239]12[240] | AGTATAAAGTTCAGCTAATGCAGATGTCTTTC | Core staple |
| 15[224]17[223] | CCTAAATCAAAATCATAGGTCTAAACAGTA | Core staple |
| 16[239]14[240] | GAATTTATTTAATGGTTTGAAATATTCTTACC | Core staple |
| 18[143]17[159] | CAACTGTTGCGCCATTCGCCATTCAAACATCA | Core staple |
| 20[79]18[80] | TTCCAGTCGTAATCATGGTCATAAAAGGGG | Core staple |
| 21[184]23[191] | TCAACAGTTGAAAGGAGCAAATGAAAAATCTAGAGATAGA | Core staple |
| 22[207]20[208] | AGCCAGCAATTGAGGAAGGTTATCATCATTTT | Core staple |
| 23[224]22[240] | GCACAGACAATATTTTTGAATGGGGTCAGTA | Core staple |
| 14[271]12[272] | TTAGTATCACAATAGATAAGTCCACGAGCA | Core staple |

| | | |
|---|---|---|
| 15[256]18[256] | GTGATAAAAAGACGCTGAGAAGAGATAACCTTGCTTCTGTTCGGGAGA | Core staple |
| 17[32]19[31] | TGCATCTTTCCCAGTCACGACGGCCTGCAG | Core staple |
| 18[175]16[176] | CTGAGCAAAAATTAATTACATTTTGGGTTA | Core staple |
| 20[143]19[159] | AAGCCTGGTACGAGCCGGAAGCATAGATGATG | Core staple |
| 21[224]23[223] | CTTTAGGGCCTGCAACAGTGCCAATACGTG | Core staple |
| 22[239]20[240] | TTAACACCAGCACTAACAACTAATCGTTATTA | Core staple |
| 23[256]22[272] | CTTTAATGCGCGAACTGATAGCCCCACCAG | Core staple |
| 15[32]17[31] | TAATCAGCGGATTGACCGTAATCGTAACCG | Core staple |
| 16[47]14[48] | ACAAACGGAAAAGCCCCAAAAACACTGGAGCA | Core staple |
| 17[96]19[95] | GCTTTCCGATTACGCCAGCTGGCGGCTGTTTC | Core staple |
| 18[207]16[208] | CGCGCAGATTACCTTTTTTAATGGGAGAGACT | Core staple |
| 20[207]18[208] | GCGGAACATCTGAATAATGGAAGGTACAAAAT | Core staple |
| 21[248]23[255] | AGATTAGAGCCGTCAAAAAACAGAGGTGAGGCCTATTAGT | Core staple |
| 23[32]22[48] | CAAATCAAGTTTTTTGGGGTCGAAACGTGGA | Core staple |
| 0[143]1[127] | TCTAAAGTTTTGTCGTCTTTCCAGCCGACAA | Core staple |
| 15[64]18[64] | GTATAAGCCAACCCGTCGGATTCTGACGACAGTATCGGCCGCAAGGCG | Core staple |
| 16[79]14[80] | GCGAGTAAAAATATTTAAATTGTTACAAAG | Core staple |
| 17[160]18[144] | AGAAAACAAAGAAGATGATGAAACAGGCTGCG | Core staple |
| 18[239]16[240] | CCTGATTGCAATATATGTGAGTGATCAATAGT | Core staple |
| 21[32]23[31] | TTTTCACTCAAAGGGCGAAAAACCATCACC | Core staple |
| 22[47]20[48] | CTCCAACGCAGTGAGACGGGCAACCAGCTGCA | Core staple |
| 23[64]22[80] | AAAGCACTAAATCGGAACCCTAATCCAGTT | Core staple |
| 0[207]1[191] | TCACCAGTACAAACTACAACGCCTAGTACCAG | Core staple |
| 4[47]2[48] | GACCAACTAATGCCACTACGAAGGGGGTAGCA | Core staple |
| 20[47]18[48] | TTAATGAACTAGAGGATCCCCGGGGGGTAACG | Core staple |
| 4[111]2[112] | GACCTGCTCTTTGACCCCCAGCGAGGGAG | Core staple |
| 20[111]18[112] | CACATTAAAATTGTTATCCGCTCATGCGGGCC | Core staple |
| 4[175]2[176] | CACCAGAAAGGTTGAGGCAGGTCATGAAAG | Core staple |
| 20[175]18[176] | ATTATCATTCAATATAATCCTGACAATTAC | Core staple |
| 4[239]2[240] | GCCTCCCTCAGAATGGAAAGCGCAGTAACAGT | Core staple |
| 20[239]18[240] | ATTTTAAAATCAAAATTATTTGCACGGATTCG | Core staple |
| 12[47]10[48] | TAAATCGGGATTCCCAATTCTGCGATATAATG | Core staple |
| 12[111]10[112] | TAAATCATATAACCTGTTTAGCTAACCTTTAA | Core staple |
| 12[175]10[176] | TTTTATTTAAGCAAATCAGATATTTTTTGT | Core staple |
| 12[239]10[240] | CTTATCATTCCCGACTTGCGGGAGCCTAATT | Core staple |
| 4[63]6[56] | TTTTATAAGGGAACCGGATATTCATTACGTCAGGACGTTGGGAA | Core staple |
| 4[127]6[120] | TTTTTTGTGTCGTGACGAGAAACACCAAATTTCAACTTTAAT | Core staple |
| 4[191]6[184] | TTTTCACCCTCAGAAACCATCGATAGCATTGAGCCATTTGGGAA | Core staple |
| 4[255]6[248] | TTTTAGCCACCACTGTAGCGCGTTTTCAAGGGAGGGAAGGTAAA | Core staple |
| 18[63]20[56] | TTTTATTAAGTTTACCGAGCTCGAATTCGGGAAACCTGTCGTGC | Core staple |
| 18[127]20[120] | TTTTGCGATCGGCAATTCCACACAACAGGTGCCTAATGAGTG | Core staple |
| 18[191]20[184] | TTTTATTCATTTTTGTTTGGATTATACTAAGAAACCACCAGAAG | Core staple |
| 18[255]20[248] | TTTTAACAATAACGTAAAACAGAAATAAAAATCCTTTGCCCGAA | Core staple |
| biotin-4[63]6[56] | /5Biosg/TTTTATAAGGGAACCGGATATTCATTACGTCAGGACGTTGGGAA | Biotinylated staple |
| biotin-4[127]6[120] | /5Biosg/TTTTTTGTGTCGTGACGAGAAACACCAAATTTCAACTTTAAT | Biotinylated staple |
| biotin-4[191]6[184] | /5Biosg/TTTTCACCCTCAGAAACCATCGATAGCATTGAGCCATTTGGGAA | Biotinylated staple |
| biotin-4[255]6[248] | /5Biosg/TTTTAGCCACCACTGTAGCGCGTTTTCAAGGGAGGGAAGGTAAA | Biotinylated staple |
| biotin-18[63]20[56] | /5Biosg/TTTTATTAAGTTTACCGAGCTCGAATTCGGGAAACCTGTCGTGC | Biotinylated staple |
| biotin-18[127]20[120] | /5Biosg/TTTTGCGATCGGCAATTCCACACAACAGGTGCCTAATGAGTG | Biotinylated staple |
| biotin-18[191]20[184] | /5Biosg/TTTTATTCATTTTTGTTTGGATTATACTAAGAAACCACCAGAAG | Biotinylated staple |
| biotin-18[255]20[248] | /5Biosg/TTTTAACAATAACGTAAAACAGAAATAAAAATCCTTTGCCCGAA | Biotinylated staple |
| 4[47]2[48]-2T-R1 | GACCAACTAATGCCACTACGAAGGGGGTAGCATTTCCTCCTCCTCCTCCTCCT | 20 nm docking strand with 32 nt binder |
| 20[47]18[48]-2T-R1 | TTAATGAACTAGAGGATCCCCGGGGGGTAACGTTTCCTCCTCCTCCTCCTCCT | 20 nm docking strand with 32 nt binder |
| 4[111]2[112]-2T-R1 | GACCTGCTCTTTGACCCCCAGCGAGGGAGTTATTTCCTCCTCCTCCTCCTCCT | 20 nm docking strand with 32 nt binder |
| 20[111]18[112]-2T-R1 | CACATTAAAATTGTTATCCGCTCATGCGGGCCTTTCCTCCTCCTCCTCCTCCT | 20 nm docking strand with 32 nt binder |
| 4[175]2[176]-2T-R1 | CACCAGAAAGGTTGAGGCAGGTCATGAAAGTTTCCTCCTCCTCCTCCTCCT | 20 nm docking strand with 32 nt binder |
| 20[175]18[176]-2T-R1 | ATTATCATTCAATATAATCCTGACAATTACTTTCCTCCTCCTCCTCCTCCT | 20 nm docking strand with 32 nt binder |
| 4[239]2[240]-2T-R1 | GCCTCCCTCAGAATGGAAAGCGCAGTAACAGTTTTCCTCCTCCTCCTCCTCCT | 20 nm docking strand with 32 nt binder |
| 20[239]18[240]-2T-R1 | ATTTTAAAATCAAAATTATTTGCACGGATTCGTTTCCTCCTCCTCCTCCTCCT | 20 nm docking strand with 32 nt binder |
| 12[47]10[48]-2T-R1 | TAAATCGGGATTCCCAATTCTGCGATATAATGTTTCCTCCTCCTCCTCCTCCT | 20 nm docking strand with 32 nt binder |
| 12[111]10[112]-2T-R1 | TAAATCATATAACCTGTTTAGCTAACCTTTAATTTCCTCCTCCTCCTCCTCCT | 20 nm docking strand with 32 nt binder |
| 12[175]10[176]-2T-R1 | TTTTATTTAAGCAAATCAGATATTTTTTGTTTTCCTCCTCCTCCTCCTCCT | 20 nm docking strand with 32 nt binder |
| 12[239]10[240]-2T-R1 | CTTATCATTCCCGACTTGCGGGAGCCTAATTTTTTCCTCCTCCTCCTCCTCCT | 20 nm docking strand with 32 nt binder |
| 6[47]2[48]-2T-R1 | TACGTTAAAGTAATCTTGACAAGAACCGAACTGACCAACTAATGCCACTACGA AGGGGGTAGCATTTCCTCCTCCTCCTCCTCCT | 10 nm, 14 nm, and 20 nm docking strand with 64 nt binder |
| 14[47]10[48]-2T-R1 | AACAAGAGGGATAAAAATTTTTAGCATAAAGCTAAATCGGGATTCCCAATTCTG CGATATAATGTTTCCTCCTCCTCCTCCTCCT | 10 nm, 14 nm, and 20 nm docking strand with 64 nt binder |
| 22[47]18[48]-2T-R1 | CTCCAACGCAGTGAGACGGGCAACCAGCTGCATTAATGAACTAGAGGATCCCCGG GGGGTAACGTTTCCTCCTCCTCCTCCTCCT | 10 nm, 14 nm, and 20 nm docking strand with 64 nt binder |
| 6[111]2[112]-2T-R1 | ATTACCTTTGAATAAGGCTTGCCCAAATCCGCGACCTGCTCTTTGACCCCCAGCG AGGGAGTTATTTCCTCCTCCTCCTCCTCCT | 10 nm, 14 nm, and 20 nm docking strand with 64 nt binder |

| | | |
|---|---|---|
| 14[111]10[112]-2T-R1 | GAGGGTAGGATTCAAAAGGGTGAGACATCCAATAAATCATATAACCTGTTTAGCTA<br>ACCTTTAATTTCCTCCTCCTCCTCCTCCT | 10 nm, 14 nm, and 20 nm docking strand with 64 nt binder |
| 22[111]18[112]-2T-R1 | GCCCGAGAGTCCACGCTGGTTTGCAGCTAACTCACATTAAAATTGTTATCCGCTCAT<br>GCGGGCCTTTCCTCCTCCTCCTCCTCCT | 10 nm, 14 nm, and 20 nm docking strand with 64 nt binder |
| 6[175]2[176]-2T-R1 | CAGCAAAAGGAAACGTCACCAATGAGCCGCCACCAGAAAGGTTGAGGCAGGTCATGA<br>AAGTTTCCTCCTCCTCCTCCTCCT | 10 nm, 14 nm, and 20 nm docking strand with 64 nt binder |
| 14[175]10[176]-2T-R1 | CATGTAATAGAATATAAAGTACCAAGCCGTTTTTATTTAAGCAAATCAGATATTTTTT<br>GTTTTCCTCCTCCTCCTCCTCCT | 10 nm, 14 nm, and 20 nm docking strand with 64 nt binder |
| 22[175]18[176]-2T-R1 | ACCTTGCTTGGTCAGTTGGCAAAGAGCGGAATTATCATTCAATATAATCCTGACAATTA<br>CTTTCCTCCTCCTCCTCCTCCT | 10 nm, 14 nm, and 20 nm docking strand with 64 nt binder |
| 6[239]2[240]-2T-R1 | GAAATTATTGCCTTTAGCGTCAGACCGGAACCGCCTCCCTCAGAATGGAAAGCGCAGTA<br>ACAGTTTTCCTCCTCCTCCTCCTCCT | 10 nm, 14 nm, and 20 nm docking strand with 64 nt binder |
| 14[239]10[240]-2T-R1 | AGTATAAAGTTCAGCTAATGCAGATGTCTTTCCTTATCATTCCCGACTTGCGGGAGCCTA<br>ATTTTTTCCTCCTCCTCCTCCTCCT | 10 nm, 14 nm, and 20 nm docking strand with 64 nt binder |
| 22[239]18[240]-2T-R1 | TTAACACCAGCACTAACAACTAATCGTTATTAATTTTAAAATCAAAATTATTTGCACGGA<br>TTCGTTTCCTCCTCCTCCTCCTCCT | 10 nm, 14 nm, and 20 nm docking strand with 64 nt binder |
| 10[47]6[48]-2T-R1 | CTGTAGCTTGACTATTATAGTCAGTTCATTGAATCCCCCTATACCACATTCAACTAGAAA<br>AATCTTTCCTCCTCCTCCTCCTCCT | 10 nm, and 14 nm docking strand with 64 nt binder |
| 18[79]14[80]-2T-R1 | GATGTGCTTCAGGAAGATCGCACAATGTGAGCGAGTAAAAATATTTAAATTGTTACAAAG<br>TTTCCTCCTCCTCCTCCTCCT | 10 nm, and 14 nm docking strand with 64 nt binder |
| 21[56]22[80]-2T-R1 | GCTGATTGCCCTTCAGAGTCCACTATTAAAGGGTGCCGTAAAGCACTAAATCGGAACCCTA<br>ATCCAGTTTTTCCTCCTCCTCCTCCTCCT | 10 nm, and 14 nm docking strand with 64 nt binder |
| 6[143]6[144]-2T-R1 | GATGGTTTGAACGAGTAGTAAATTTACCATTAGCAAGGCCTCACCAGTAGCACCATGGGCTT<br>GATTTCCTCCTCCTCCTCCTCCT | 10 nm, and 14 nm docking strand with 64 nt binder |
| 14[143]14[144]-2T-R1 | CAACCGTTTCAAATCACCATCAATTCGAGCCAGTAATAAGTTAGGCAGAGGCATTTATGATA<br>TTTTTCCTCCTCCTCCTCCTCCT | 10 nm, and 14 nm docking strand with 64 nt binder |
| 22[143]22[144]-2T-R1 | TCGGCAAATCCTGTTTGATGGTGGACCCTCAATCAATATCGAACCTCAAATATCAATTCCGA<br>AATTTCCTCCTCCTCCTCCTCCT | 10 nm, and 14 nm docking strand with 64 nt binder |
| 10[207]6[208]-2T-R1 | ATCCCAATGAGAATTAACTGAACAGTTACCAGAAGGAAACATAAAGGTGGCAACATTATCACC<br>GTTTCCTCCTCCTCCTCCTCCT | 10 nm, and 14 nm docking strand with 64 nt binder |
| 18[207]14[208]-2T-R1 | CGCGCAGATTACCTTTTTTAATGGGAGAGACTACCTTTTTATTTTAGTTAATTTCATAGG<br>GCTTTTTCCTCCTCCTCCTCCTCCT | 10 nm, and 14 nm docking strand with 64 nt binder |
| 21[184]22[208]-2T-R1 | ACAGTTGAAAGGAGCAAATGAAAAATCTAGAGATAGAACCCTTCTGACCTGAAAGCGTAA<br>GACGCTGAGTTTCCTCCTCCTCCTCCTCCT | 10 nm, and 14 nm docking strand with 64 nt binder |
| 10[271]6[272]-2T-R1 | ACGCTAACACCCACAAGAATTGAAAATAGCAATAGCTATCAATAGAAAATTCAACAT<br>TCATTTCCTCCTCCTCCTCCTCCT | 10 nm, and 14 nm docking strand with 64 nt binder |
| 18[271]14[272]-2T-R1 | CTTTTACAAAATCGTCGCTATTAGCGATAGCTTAGATTTAAGGCGTTAAATAAAGCCT<br>GTTTTCCTCCTCCTCCTCCTCCT | 10 nm, and 14 nm docking strand with 64 nt binder |
| 21[248]22[272]-2T-R1 | GATTAGAGCCGTCAAAAAACAGAGGTGAGGCCTATTAGTCTTTAATGCGCGAACTGA<br>TAGCCCCACCAGTTTCCTCCTCCTCCTCCTCCT | 10 nm, and 14 nm docking strand with 64 nt binder |
| 10[111]6[112]-2T-R1 | TTGCTCCTTTCAAATATCGCGTTTGAGGGGGTAATAGTAAACACTATCATAACCCTC<br>ATTGTGATTTCCTCCTCCTCCTCCTCCT | 10 nm docking strand with 64 nt binder |
| 10[143]10[144]-2T-R1 | CCAACAGGAGCGAACCAGACCGGAGCCTTTACAGAGAGAAAAAAATGAAAATAGCAA<br>GCAAACTTTTCCTCCTCCTCCTCCTCCT | 10 nm docking strand with 64 nt binder |
| 10[175]6[176]-2T-R1 | TTAACGTCTAACATAAAAACAGGTAACGGAATACCCAACAGTATGTTAGCAAATTAG<br>AGCTTTCCTCCTCCTCCTCCTCCT | 10 nm docking strand with 64 nt binder |
| 10[239]6[240]-2T-R1 | GCCAGTTAGAGGGTAATTGAGCGCTTTAAGAAAAGTAAGCAGACACCACGGAATAAT<br>ATTGACGTTTCCTCCTCCTCCTCCTCCT | 10 nm docking strand with 64 nt binder |
| 14[207]10[208]-2T-R1 | AATTGAGAATTCTGTCCAGACGACTAAACCAAGTACCGCAATTCTAAGAACGCGAGT<br>ATTATTTTTTCCTCCTCCTCCTCCTCCT | 10 nm docking strand with 64 nt binder |
| 14[271]10[272]-2T-R1 | TTAGTATCACAATAGATAAGTCCACGAGCATGTAGAAATCAAGATTAGTTGCTCTT<br>ACCATTTCCTCCTCCTCCTCCTCCT | 10 nm docking strand with 64 nt binder |
| 14[79]10[80]-2T-R1 | GCTATCAGAAATGCAATGCCTGAATTAGCAAAATTAAGTTGACCATTAGATACTTT<br>TGCGTTTCCTCCTCCTCCTCCTCCT | 10 nm docking strand with 64 nt binder |
| 18[111]14[112]-2T-R1 | TCTTCGCTGCACCGCTTCTGGTGCGGCCTTCCTGTAGCCATTAAAATTCGCATTAAA<br>TGCCGGATTTCCTCCTCCTCCTCCTCCT | 10 nm docking strand with 64 nt binder |
| 18[143]18[144]-2T-R1 | CAACTGTTGCGCCATTCGCCATTCAAACATCAAGAAAACAAAGAAGATGATGAAACA<br>GGCTGCGTTTCCTCCTCCTCCTCCTCCT | 10 nm docking strand with 64 nt binder |
| 18[175]14[176]-2T-R1 | CTGAGCAAAAATTAATTACATTTTGGGTTATATAACTAACAAAGAACGCGAGAACG<br>CCAATTTCCTCCTCCTCCTCCTCCT | 10 nm docking strand with 64 nt binder |
| 18[239]14[240]-2T-R1 | CCTGATTGCAATATATGTGAGTGATCAATAGTGAATTTATTTAATGGTTTGAAATAT<br>TCTTACCTTTCCTCCTCCTCCTCCTCCT | 10 nm docking strand with 64 nt binder |
| 18[47]14[48]-2T-R1 | CCAGGGTTGCCAGTTTGAGGGGACCCGTGGGAACAAACGGAAAAGCCCCAAAAACA<br>CTGGAGCATTTCCTCCTCCTCCTCCTCCT | 10 nm docking strand with 64 nt binder |
| 2[143]2[144]-2T-R1 | ATATTCGGAACCATCGCCCACGCAGAGAAGGATTAGGATTGGCTGAGACTCCTCAA<br>TAACCGATTTTCCTCCTCCTCCTCCTCCT | 10 nm docking strand with 64 nt binder |
| 21[184]22[208]-2T-R1 | ACAGTTGAAAGGAGCAAATGAAAAATCTAGAGATAGAACCCTTCTGACCTGAAAGCG<br>TAAGACGCTGAGTTTCCTCCTCCTCCTCCTCCT | 10 nm docking strand with 64 nt binder |
| 21[224]22[240]-2T-R1 | CTTTAGGGCCTGCAACAGTGCCAATACGTGGCACAGACAATATTTTTGAATGGGGTC<br>AGTATTTCCTCCTCCTCCTCCTCCT | 10 nm docking strand with 64 nt binder |
| 21[32]22[48]-2T-R1 | TTTTCACTCAAAGGGCGAAAAACCATCACCCAAATCAAGTTTTTTGGGGTCGAAACGT<br>GGATTTCCTCCTCCTCCTCCTCCT | 10 nm docking strand with 64 nt binder |

| | | |
|---|---|---|
| 21[96]22[112]-2T-R1 | AGCAAGCGTAGGGTTGAGTGTTGTAGGGAGCCCCCGATTTAGAGCTTGACGGGGAAAA AGAATATTTCCTCCTCCTCCTCCTCCT | 10 nm docking strand with 64 nt binder |
| 22[207]18[208]-2T-R1 | AGCCAGCAATTGAGGAAGGTTATCATCATTTTGCGGAACATCTGAATAATGGAAGGT ACAAAATTTTCCTCCTCCTCCTCCTCCT | 10 nm docking strand with 64 nt binder |
| 22[271]18[272]-2T-R1 | CAGAAGATTAGATAATACATTTGTCGACAACTCGTATTAGAAATTGCGTAGATACAG TACTTTCCTCCTCCTCCTCCTCCT | 10 nm docking strand with 64 nt binder |
| 22[79]18[80]-2T-R1 | TGGAACAACCGCCTGGCCCTGAGGCCCGCTTTCCAGTCGTAATCATGGTCATAAAA GGGGTTTCCTCCTCCTCCTCCTCCT | 10 nm docking strand with 64 nt binder |
| 23[128]22[176]-2T-R1 | AACGTGGCGAGAAAGGAAGGGAAACCAGTAATAAAAGGGACATTCTGGCCAACAAA GCATCTTTCCTCCTCCTCCTCCTCCT | 10 nm docking strand with 64 nt binder |
| 6[207]2[208]-2T-R1 | TCACCGACGCACCGTAATCAGTAGCAGAACCGCCACCCTCTATTCACAAACAAATA CCTGCCTATTTCCTCCTCCTCCTCCTCCT | 10 nm docking strand with 64 nt binder |
| 6[271]2[272]-2T-R1 | ACCGATTGTCGGCATTTTCGGTCATAATCAAAATCACCTTCCAGTAAGCGTCAGT AATAATTTCCTCCTCCTCCTCCTCCT | 10 nm docking strand with 64 nt binder |
| 6[79]2[80]-2T-R1 | TTATACCACCAAATCAACGTAACGAACGAGGCGCAGACAAGAGGCAAAAGAATCCCT CAGTTTCCTCCTCCTCCTCCTCCT | 10 nm docking strand with 64 nt binder |

**Table S1** **2D RRO strands.** Scaffold is MP13mp18. Core staples are all the strands that form the structure. Biotinylated staples are modified with biotin to immobilize origami on coverslip surfaces through a BSA-Biotin-Streptavidin-Biotin-DNA origami arrangement. To fold a 20 nm pattern on 2D RRO with 64 binders, we mix scaffold strands; the 10 nm, 14 nm, and 20 nm strands with 64 nt binders along with core staples (positions corresponding to 20 nm docking positions and biotin positions should be excluded beforehand) and biotinylated staple strands. To fold a 14 nm pattern on 2D RRO with 64 binder, we mix scaffold strands, the 10 nm, 14 nm, and 20 nm strands with 64 nt binder, the 10 nm, and 14 nm docking strands with 64 nt binder along with core staples (positions corresponding to 20 nm docking positions and biotin positions should be excluded beforehand). To fold a 10 nm pattern on 2D RRO with 64 binder, we mix scaffold strands, the 10 nm, 14 nm, and 20 nm strands with 64 nt binder, the 10 nm, and 14 nm docking strands with 64 nt binder, the 10 nm docking strands with 64 nt binder along with core staples (positions corresponding to 20 nm docking positions and biotin positions should be excluded beforehand).

| Name | Sequence | Note |
|---|---|---|
| 06_cuboctahedron_147_1-1913-V | ATCACCGTACTTTTTTCAGGAGGTTTTAAAGATTCAATTTTTAAGGGTGAGA | Core staple |
| 06_cuboctahedron_147_1-4390-E | CAGTAACAGTAGTATAGCCCGGAATAGGTGTAGATGAATATA | Core staple |
| 06_cuboctahedron_147_1-4369-E | CGGGAGAAACGCCGTCGAGAGGGTTGATATAACCTTTTACAT | Core staple |
| 06_cuboctahedron_147_1-4348-E | CGCCTGATTGCTCAGTACCAGGCGGATAAGTAATAACGGATT | Core staple |
| 06_cuboctahedron_147_1-4327-E | AAGTTACAAAATTAGGATTAGCGGGGTTTTGCTTTGAATACC | Core staple |
| 06_cuboctahedron_147_1-4306-E | GCGAATTATTCTGAGACTCCTCAAGAGAAGGATCGCGCAGAG | Core staple |
| 06_cuboctahedron_147_1-4285-E | CCTGAGCAAAAAACATGAAAGTATTAAGAGGCATTTCAATTA | Core staple |
| 06_cuboctahedron_147_2-2060-V | ATTTCGGAACCTTTTTTATTATTCTGAGAAGATGATGTTTTTAAACAAACAT | Core staple |
| 06_cuboctahedron_147_2-1609-V | ACTAAAGGAATTTTTTTGCGAATAATGTTTAATTTCATTTTTACTTTAATCA | Core staple |
| 06_cuboctahedron_147_2-1669-E | GTATGGGACAGACGTTAGTAAATGTAACGGGG | Core staple |
| 06_cuboctahedron_147_2-2110-E | TCAGTGCCTACTGGTAATAAGTTTAATTTTCT | Core staple |
| 06_cuboctahedron_147_2-2165-E | GTCATACATGAACAGTTAATGCCCCCTGCCTTTCCAGTAAGC | Core staple |
| 06_cuboctahedron_147_2-2144-E | TACAGGAGTGTTGAGTAACAGTGCCCGTATAGCTTTTGATGA | Core staple |
| 06_cuboctahedron_147_2-1650-E | AACTTTCAACTCTAAAGTTTTGTCGTCTTTCTTTTGCTAAAC | Core staple |
| 06_cuboctahedron_147_2-1629-E | AGTGAGAATACCCTCATAGTTAGCGTAACGAAGTTTCAGCGG | Core staple |
| 06_cuboctahedron_147_2-1619-E | GAAAGGAACACCACAGACAG | Core staple |
| 06_cuboctahedron_147_3-5755-V | AGGGCGAAAAATTTTTCCGTCTATCATAGATTTTCAGTTTTTGTTTAACGTC | Core staple |
| 06_cuboctahedron_147_3-7016-E | CAGTCAAATCGAACGTGGACTCCAACGTCAAAAGGCCGGAGA | Core staple |
| 06_cuboctahedron_147_3-6995-E | GATATTCAACGAACAAGAGTCCACTATTAAAACCATCAATAT | Core staple |
| 06_cuboctahedron_147_3-6974-E | ATAAATTAATGTTGAGTGTTGTTCCAGTTTGCGTTCTAGCTG | Core staple |
| 06_cuboctahedron_147_3-6953-E | TAGCTATTTTAAAAGAATAGCCCGAGATAGGGCCGGAGAGGG | Core staple |
| 06_cuboctahedron_147_3-6932-E | CAAAGGCTATCGGCAAAATCCCTTATAAATCTGAGAGATCTA | Core staple |
| 06_cuboctahedron_147_3-6911-E | CTGAGAGTCTGTTTGATGGTGGTTCCGAAATCAGGTCATTGC | Core staple |
| 06_cuboctahedron_147_4-5902-V | GCCCCAGCAGGTTTTTCGAAAATCCTGGAGCAAACAATTTTTGAGAATCGAT | Core staple |
| 06_cuboctahedron_147_4-5451-V | TTCCTCGTTAGTTTTTAATCAGAGCGACATTTGAGGATTTTTTTTAGAAGTA | Core staple |
| 06_cuboctahedron_147_4-5511-E | CTTAATGCCGCGTAACCACCACACTGATTGCC | Core staple |
| 06_cuboctahedron_147_4-5952-E | CTTCACCGTGAGACGGGCAACAGCCCGCCGCG | Core staple |
| 06_cuboctahedron_147_4-6007-E | TGGGCGCCAGGCAAGCGGTCCACGCTGGTTTGGTTTGCGTAT | Core staple |
| 06_cuboctahedron_147_4-5986-E | TTTTCACCAGCCTGGCCCTGAGAGAGTTGCAGGTGGTTTTTC | Core staple |
| 06_cuboctahedron_147_4-5492-E | GCGCGTACTAGCAAGTGTAGCGGTCACGCTGGCCGCTACAGG | Core staple |
| 06_cuboctahedron_147_4-5471-E | ACGAGCACGTGGAGCGGGCGCTAGGGCGCTGTGGTTGCTTTG | Core staple |
| 06_cuboctahedron_147_4-5461-E | ATAACGTGCTGAAAGCGAAA | Core staple |
| 06_cuboctahedron_147_5-3530-V | TAGAAACCAATTTTTTCAATAATCGGGCGCAGTCTCTTTTTTGAATTTACCG | Core staple |
| 06_cuboctahedron_147_5-4128-V | TTCCCTTAGAATTTTTTCCTTGAAAAAATCGCAAGACTTTTTAAAGAACGCG | Core staple |
| 06_cuboctahedron_147_5-4243-E | AATTAATTACCCATCCTAATTTACGAGCATGCAAGAAAACAA | Core staple |
| 06_cuboctahedron_147_5-4222-E | TCATTTGAATCTGAACAAGAAAAATAATATCATTTAACAATT | Core staple |
| 06_cuboctahedron_147_5-4201-E | ATGGAAACAGTTTATCAACAATAGATAAGTCTACCTTTTTTA | Core staple |
| 06_cuboctahedron_147_5-4180-E | ATATATGTGAAGCTAATGCAGAACGCGCCTGTACATAAATCA | Core staple |
| 06_cuboctahedron_147_5-4159-E | TGCTTCTGTAACGACAATAAACAACATGTTCGTGAATAACCT | Core staple |
| 06_cuboctahedron_147_5-4138-E | TTAATTAATTTAAAGTAATTCTGTCCAGACGAATCGTCGCTA | Core staple |
| 06_cuboctahedron_147_6-1462-V | GACAACAACCATTTTTTCGCCCACGCTTCATGAGGAATTTTTGTTTCCATTA | Core staple |
| 06_cuboctahedron_147_6-1577-E | GTTGAAAATCTAGTAAATTGGGCTTGAGATGAATTTTTTCAC | Core staple |
| 06_cuboctahedron_147_6-1556-E | GGCTCCAAAAGACGAGAAACACCAGAACGAGTCCAAAAAAAA | Core staple |
| 06_cuboctahedron_147_6-1535-E | TTGTATCGGTTCAGTGAATAAGGCTTGCCCTGGAGCCTTTAA | Core staple |
| 06_cuboctahedron_147_6-1514-E | CTTTCGAGGTCAACGTAACAAAGCTGCTCATTTATCAGCTTG | Core staple |
| 06_cuboctahedron_147_6-1493-E | ACAGCTTGATCCGGATATTCATTACCCAAATGAATTTCTTAA | Core staple |
| 06_cuboctahedron_147_6-1472-E | CGCCGACAATCAAGAGTAATCTTGACAAGAAACCGATAGTTG | Core staple |
| 06_cuboctahedron_147_7-6764-V | TGTTAAAATTCTTTTTGCATTAAATTGGCCAACGCGCTTTTTGGGGAGAGGC | Core staple |
| 06_cuboctahedron_147_7-286-V | AGAGTACCTTTTTTTTAATTGCTCCTTTGAGATTTAGTTTTTGAATACCACA | Core staple |
| 06_cuboctahedron_147_7-346-E | TTTTAATTGCCCGAAAGACTTCAAAGCCCCAA | Core staple |
| 06_cuboctahedron_147_7-6814-E | AAACAGGAGGTTGATAATCAGAAAATATCGCG | Core staple |
| 06_cuboctahedron_147_7-6869-E | GTAAAACTAGAAATTGTAAACGTTAATATTTGAACGGTAATC | Core staple |
| 06_cuboctahedron_147_7-6848-E | TATGTACCCCAGATTGTATAAGCAAATATTTCATGTCAATCA | Core staple |
| 06_cuboctahedron_147_7-327-E | GCGAACCAGACATCAAAAAGATTAAGAGGAACGAGCTTCAAA | Core staple |
| 06_cuboctahedron_147_7-306-E | CTCCAACAGGAGTCAGAAGCAAAGCGGATTGCCGGAAGCAAA | Core staple |
| 06_cuboctahedron_147_7-296-E | TCAGGATTAGTGACTATTAT | Core staple |
| 06_cuboctahedron_147_8-6460-V | AACCAGGCAAATTTTTGCGCCATTCGTTCCCAGTCACTTTTTGACGTTGTAA | Core staple |
| 06_cuboctahedron_147_8-6520-E | GTATCGGCTGCCAGTTTGAGGGGAATTCATTG | Core staple |
| 06_cuboctahedron_147_8-493-E | AATCCCCCCGGAATCGTCATAAATCGACGACA | Core staple |
| 06_cuboctahedron_147_8-548-E | AAATGTTTAGGAAAACGAGAATGACCATAAAGGGTAATAGTA | Core staple |
| 06_cuboctahedron_147_8-527-E | TCCAATACTGTCAAATGCTTTAAACAGTTCAACTGGATAGCG | Core staple |
| 06_cuboctahedron_147_8-6501-E | CGCACTCCAGGGCGCATCGTAACCGTGCATCCTCAGGAAGAT | Core staple |
| 06_cuboctahedron_147_8-6480-E | GCACCGCTTCTAGGTCACGTTGGTGTAGATGCCAGCTTTCCG | Core staple |
| 06_cuboctahedron_147_8-6470-E | TGGTGCCGGACGTAATGGGA | Core staple |
| 06_cuboctahedron_147_9-5304-V | TGTAGCAATACTTTTTTTCTTTGATTTGAAATGGATTTTTTTATTTACATTG | Core staple |
| 06_cuboctahedron_147_9-5419-E | GGAGGCCGATTTAGAGCCGTCAATAGATAATGGAGCTAAACA | Core staple |
| 06_cuboctahedron_147_9-5398-E | TAGACAGGAAGGAGCACTAACAACTAATAGATAAAGGGATTT | Core staple |
| 06_cuboctahedron_147_9-5377-E | AATCCTGAGAAGGTTATCTAAAATATCTTTACGGTACGCCAG | Core staple |
| 06_cuboctahedron_147_9-5356-E | AATCAGTGAGACAGTTGAAAGGAATTGAGGAAGTGTTTTTAT | Core staple |
| 06_cuboctahedron_147_9-5335-E | AAAGAGTCTGATCTGGTCAGTTGGCAAATCAGCCACCGAGTA | Core staple |

| | | |
|---|---|---|
| 06_cuboctahedron_147_9-5314-E | AATTAACCGTAATATCAAACCCTCAATCAATTCCATCACGCA | Core staple |
| 06_cuboctahedron_147_10-3099-V | AGCGCATTAGATTTTTCGGGAGAATTTAAGAAAAGTATTTTTAGCAGATAGC | Core staple |
| 06_cuboctahedron_147_10-3214-E | GTTACAAAATCAGAATCAAGTTTGCCTTTAGCTAATTTGCCA | Core staple |
| 06_cuboctahedron_147_10-3193-E | TTATTTATCCCAGCACCGTAATCAGTAGCGAAAACAGCCATA | Core staple |
| 06_cuboctahedron_147_10-3172-E | AGAAACGATTCACCAATGAAACCATCGATAGCAATCCAAATA | Core staple |
| 06_cuboctahedron_147_10-3151-E | GTCAAAAATGCATTAGCAAGGCCGGAAACGTTTTTGTTTAAC | Core staple |
| 06_cuboctahedron_147_10-3130-E | CTTTACAGAGAATCACCAGTAGCACCATTACAAAATAGCAGC | Core staple |
| 06_cuboctahedron_147_10-3109-E | AAAACAGGGATTTGGGAATTAGAGCCAGCAAAGAATAACATA | Core staple |
| 06_cuboctahedron_147_11-2785-V | AAAAGAAACGCTTTTTAAAGACACCACACCGTCACCGTTTTTACTTGAGCCA | Core staple |
| 06_cuboctahedron_147_11-2845-E | TCCTTATTCAAAAGAACTGGCATGGCCAGCTG | Core staple |
| 06_cuboctahedron_147_11-6373-E | GCGAAAGGGCCTCTTCGCTATTACATTAAGAC | Core staple |
| 06_cuboctahedron_147_11-6428-E | GCGCAACTGTAAGTTGGGTAACGCCAGGGTTCCATTCAGGCT | Core staple |
| 06_cuboctahedron_147_11-6407-E | ATCGGTGCGGGGGATGTGCTGCAAGGCGATTTGGGAAGGGCG | Core staple |
| 06_cuboctahedron_147_11-2826-E | TAGCAAACGTACGCAATAATAACGGAATACCACGCAGTATGT | Core staple |
| 06_cuboctahedron_147_11-2805-E | ACATAAAGGTTACCAGAAGGAAACCGAGGAAAGAAAATACAT | Core staple |
| 06_cuboctahedron_147_11-2795-E | GGCAACATATCGAACAAAGT | Core staple |
| 06_cuboctahedron_147_12-1881-E | CCTCAGAACCTAGTAGTAGCATTTGTGTAGGAGTACCGCCAC | Core staple |
| 06_cuboctahedron_147_12-1860-E | AACCGCCACCAGGTGGCATCAATTCTACTAAGCCACCCTCAG | Core staple |
| 06_cuboctahedron_147_12-1839-E | CACCCTCATTCATTTGGGGCGCGAGCTGAAACTCAGAGCCAC | Core staple |
| 06_cuboctahedron_147_12-1818-E | CAAGCCCAATTAACCTGTTTAGCTATATTTTTTCAGGGATAG | Core staple |
| 06_cuboctahedron_147_12-1797-E | TACCGTAACAATACATTTCGCAAATGGTCAAAGGAACCCATG | Core staple |
| 06_cuboctahedron_147_12-1776-E | CACCAGTACATAGATTTAGTTTGACCATTAGCTGAGTTTCGT | Core staple |
| 06_cuboctahedron_147_13-139-V | ATTCCCAATTCTTTTTTGCGAACGAGAACTACAACGCTTTTTCTGTAGCATT | Core staple |
| 06_cuboctahedron_147_13-832-E | CTTATGCGATTTTCATTCCATATAACAGTTGTTGTGAATTAC | Core staple |
| 06_cuboctahedron_147_13-811-E | GCTCATTATACTAAAGTACGGTGTCTGGAAGTTTAAGAACTG | Core staple |
| 06_cuboctahedron_147_13-790-E | GTTGGGAAGACAACATGTTTTAAATATGCAACCAGTCAGGAC | Core staple |
| 06_cuboctahedron_147_13-769-E | TAATAAAACGGCTGAATATAATGCTGTAGCTAAAATCTACGT | Core staple |
| 06_cuboctahedron_147_13-748-E | CAACATTATTCGGATGGCTTAGAGCTTAATTAACTAACGGAA | Core staple |
| 06_cuboctahedron_147_13-727-E | GATTCATCAGTTTGATAAGAGGTCATTTTTGACAGGTAGAAA | Core staple |
| 06_cuboctahedron_147_14-5723-E | CACTACGTGAAACAGAAATAAAGAAATTGCGGGGCGATGGCC | Core staple |
| 06_cuboctahedron_147_14-5702-E | AATCAAGTTTATCAAAATTATTTGCACGTAAACCATCACCCA | Core staple |
| 06_cuboctahedron_147_14-5681-E | GGTGCCGTAAGGAAGGGTTAGAACCTACCATTTTGGGGTCGA | Core staple |
| 06_cuboctahedron_147_14-5660-E | GGAACCCTAATGGATTATACTTCTGAATAATAGCACTAAATC | Core staple |
| 06_cuboctahedron_147_14-5639-E | GATTTAGAGCATCAATATAATCCTGATTGTTAGGGAGCCCCC | Core staple |
| 06_cuboctahedron_147_14-5618-E | AGCCGGCGAAATTATCAGATGATGGCAATTCTTGACGGGGAA | Core staple |
| 06_cuboctahedron_147_15-4569-V | GGAATTATCATTTTTTCATATTCCTGCGTGGCGAGAATTTTTAGGAAGGGAA | Core staple |
| 06_cuboctahedron_147_15-4031-E | CTTTTTAAAAAATCATAGGTCTGATTTTAAAA | Core staple |
| 06_cuboctahedron_147_15-4619-E | GTTTGAGTGCCCGAACGTTATTAAGAGACTAC | Core staple |
| 06_cuboctahedron_147_15-4674-E | AAACAATTCGAAAGAAACCACCAGAAGGAGCTTAGACTTTAC | Core staple |
| 06_cuboctahedron_147_15-4653-E | TAAATCCTTTAACATTATCATTTTGCGGAACACAACTCGTAT | Core staple |
| 06_cuboctahedron_147_15-4012-E | GGTTGGGTTAAGAGTCAATAGTGAATTTATCCCTCCGGCTTA | Core staple |
| 06_cuboctahedron_147_15-3991-E | GTAAATGCTGGCTTAGATTAAGACGCTGAGATATAACTATAT | Core staple |
| 06_cuboctahedron_147_15-3981-E | ATGCAAATCCCATAGCGATA | Core staple |
| 06_cuboctahedron_147_16-3498-E | TATCATTCCATCATTAAAGCCAGAATGGAAACTGTCTTTCCT | Core staple |
| 06_cuboctahedron_147_16-3477-E | TAAACCAAGTTATTCACAAACAAATAAATCCAGAACGGGTAT | Core staple |
| 06_cuboctahedron_147_16-3456-E | CGAGAACAAGAGGTCAGACGATTGGCCTTGAACCGCACTCAT | Core staple |
| 06_cuboctahedron_147_16-3435-E | TATTTTCATCGCATTGACAGGAGGTTGAGGCCAAGCCGTTTT | Core staple |
| 06_cuboctahedron_147_16-3414-E | TACCGCGCCCCCACCACCAGAGCCGCCGCCAGTAGGAATCAT | Core staple |
| 06_cuboctahedron_147_16-3393-E | AATCAGATATCCCTCAGAGCCGCCACCAGAAAATAGCAAGCA | Core staple |
| 06_cuboctahedron_147_17-2344-V | CCGCCACCCTCTTTTTAGAGCCACCAAGAAGGCTTATTTTTTCCGGTATTCT | Core staple |
| 06_cuboctahedron_147_17-1365-E | AAAGACAGGCGGGATCGTCACCCTATCACCGG | Core staple |
| 06_cuboctahedron_147_17-2394-E | AACCAGAGTCTTTTCATAATCAAACAGCAGCG | Core staple |
| 06_cuboctahedron_147_17-2449-E | TCGGTCATAGTCAGAGCCGCCACCCTCAGAACATCGGCATTT | Core staple |
| 06_cuboctahedron_147_17-2428-E | GCGTTTGCCACCACCACCGGAACCGCCTCCCCCCCCTTATTA | Core staple |
| 06_cuboctahedron_147_17-1346-E | GGGTAGCAACAGGGAGTTAAAGGCCGCTTTTCATCGGAACGA | Core staple |
| 06_cuboctahedron_147_17-1325-E | CTTTGAGGACTATTCGGTCGCTGAGGCTTGCGGCTACAGAGG | Core staple |
| 06_cuboctahedron_147_17-1315-E | TAAAGACTTTATAACCGATA | Core staple |
| 06_cuboctahedron_147_18-6732-E | AGCTCATTTTTGCCAGCTGCATTAATGAATCTTTGTTAAATC | Core staple |
| 06_cuboctahedron_147_18-6711-E | GAACGCCATCTTCCAGTCGGGAAACCTGTCGTTAACCAATAG | Core staple |
| 06_cuboctahedron_147_18-6690-E | GCGTCTGGCCGCGTTGCGCTCACTGCCCGCTAAAAATAATTC | Core staple |
| 06_cuboctahedron_147_18-6669-E | AGCTTTCATCGTGAGCTAACTCACATTAATTTTCCTGTAGCC | Core staple |
| 06_cuboctahedron_147_18-6648-E | TGAGCGAGTAAAAGCCTGGGGTGCCTAATGAAACATTAAATG | Core staple |
| 06_cuboctahedron_147_18-6627-E | GATTCTCCGTCGAGCCGGAAGCATAAAGTGTACAACCCGTCG | Core staple |
| 06_cuboctahedron_147_19-6186-V | CTCACAATTCCTTTTTACACAACATAGGGAACAAACGTTTTTGCGGATTGAC | Core staple |
| 06_cuboctahedron_147_19-5207-E | CCAGCCATGTAATATCCAGAACAAACCGAGCT | Core staple |
| 06_cuboctahedron_147_19-6236-E | CGAATTCGTAGAGGATCCCCGGGTTATTACCG | Core staple |
| 06_cuboctahedron_147_19-6291-E | GTGCCAAGCTCCTGTGTGAAATTGTTATCCGAACGACGGCCA | Core staple |
| 06_cuboctahedron_147_19-6270-E | AGGTCGACTCTAATCATGGTCATAGCTGTTTTGCATGCCTGC | Core staple |
| 06_cuboctahedron_147_19-5188-E | AAACGCTCATCTCAAACTATCGGCCTTGCTGTGCAACAGGAA | Core staple |
| 06_cuboctahedron_147_19-5167-E | CATTTTGACGTCACTTGCCTGAGTAGAAGAAGGAAATACCTA | Core staple |
| 06_cuboctahedron_147_19-5157-E | CTCAATCGTCAGTAATAACA | Core staple |

| | | |
|---|---|---|
| 06_cuboctahedron_147_20-3246-V | TAACGAGCGTCTTTTTTTTCCAGAGCCGTCAGACTGTTTTTTAGCGCGTTTT | Core staple |
| 06_cuboctahedron_147_20-3737-E | ATATTTAAAGTAGGGCTTAATTGATCAAGATT | Core staple |
| 06_cuboctahedron_147_20-3296-E | AGTTGCTAGTTTTGAAGCCTTAAAGAATCGCC | Core staple |
| 06_cuboctahedron_147_20-3351-E | GCGTTTTAGCTATCCTGAATCTTACCAACGCAAGAACGCGAG | Core staple |
| 06_cuboctahedron_147_20-3330-E | CTTGCGGGAGTTTTGCACCCAGCTACAATTTGAACCTCCCGA | Core staple |
| 06_cuboctahedron_147_20-3718-E | TGTAATTTAGAGTATAAAGCCAACGCTCAACCAACGCCAACA | Core staple |
| 06_cuboctahedron_147_20-3697-E | TTCGAGCCAGTGCGTTATACAAATTCTTACCGCAGAGGCATT | Core staple |
| 06_cuboctahedron_147_20-3687-E | TAATAAGAGATAGTATCATA | Core staple |
| 06_cuboctahedron_147_21-3834-V | AATTACTAGAATTTTTAAAGCCTGTTATATAAAGTACTTTTTCGACAAAAGG | Core staple |
| 06_cuboctahedron_147_21-4913-E | GTATTAACGATAAAACAGAGGTGATTGAAATA | Core staple |
| 06_cuboctahedron_147_21-3884-E | CCGACCGTACCTAAATTTAATGGTGGCGGTCA | Core staple |
| 06_cuboctahedron_147_21-3939-E | TCAAATATATAAGAATAAACACCGGAATCATAGAAAACTTTT | Core staple |
| 06_cuboctahedron_147_21-3918-E | TCATCTTCTGGTGATAAATAAGGCGTTAAATTTTAGTTAATT | Core staple |
| 06_cuboctahedron_147_21-4894-E | CAGTGCCACGCCGAACGAACCACCAGCAGAAACCGCCTGCAA | Core staple |
| 06_cuboctahedron_147_21-4873-E | CAGCAAATGAAAAACATCGCCATTAAAAATACTGAGAGCCAG | Core staple |
| 06_cuboctahedron_147_21-4863-E | AAAATCTAAATGATAGCCCT | Core staple |
| 06_cuboctahedron_147_22-5010-V | ATTAGTCTTTATTTTTATGCGCGAACGCATCACCTTGTTTTTCTGAACCTCA | Core staple |
| 06_cuboctahedron_147_22-3002-E | GCCCAATAGATAACCCACAAGAATAACCCTTC | Core staple |
| 06_cuboctahedron_147_22-5060-E | TGACCTGATGGCCAACAGAGATAGTGAGTTAA | Core staple |
| 06_cuboctahedron_147_22-5115-E | AGTCACACGAAGACAATATTTTTGAATGGCTGCAGATTCACC | Core staple |
| 06_cuboctahedron_147_22-5094-E | AGGGACATTCAAGCGTAAGAATACGTGGCACCCAGTAATAAA | Core staple |
| 06_cuboctahedron_147_22-2983-E | AAACAATGAAATTGAGCGCTAATATCAGAGAATAAGAGCAAG | Core staple |
| 06_cuboctahedron_147_22-2962-E | TATCTTACCGCCTGAACAAAGTCAGAGGGTAATAGCAATAGC | Core staple |
| 06_cuboctahedron_147_22-2952-E | AAGCCCTTTTAACTGAACAC | Core staple |
| 06_cuboctahedron_147_23-580-V | CAAAAGAAGTTTTTTTTTGCCAGAGGTCAAAAATCAGTTTTTGTCTTTACCC | Core staple |
| 06_cuboctahedron_147_23-1071-E | TAAGGGAAGAACGAGGCGCAGACGCTATCATA | Core staple |
| 06_cuboctahedron_147_23-630-E | ACCCTCGTCATAGTAAGAGCAACAGTCAATCA | Core staple |
| 06_cuboctahedron_147_23-685-E | CAGATACATACAAAATAGCGAGAGGCTTTTGTTCAACTAATG | Core staple |
| 06_cuboctahedron_147_23-664-E | AATTACGAGGTTACCAGACGACGATAAAAACACGCCAAAAGG | Core staple |
| 06_cuboctahedron_147_23-1052-E | AACTTTGAAACTGCTCCATGTTACTTAGCCGCCGAACTGACC | Core staple |
| 06_cuboctahedron_147_23-1031-E | AACGGTGTACAATTGTGTCGAAATCCGCGACGAGGACAGATG | Core staple |
| 06_cuboctahedron_147_23-1021-E | AGACCAGGCGTCGCCTGATA | Core staple |
| 06_cuboctahedron_147_24-1168-V | GTACAACGGAGTTTTTATTTGTATCACATAGGCTGGCTTTTTTGACCTTCAT | Core staple |
| 06_cuboctahedron_147_24-2688-E | CAACCGATCGCCAAAGACAAAAGGAGAATACA | Core staple |
| 06_cuboctahedron_147_24-1218-E | CTAAAACAAAACGAAAGAGGCAAAGCGACATT | Core staple |
| 06_cuboctahedron_147_24-1273-E | TACGTAATGCTTATACCAAGCGCGAAACAAAAACGGGTAAAA | Core staple |
| 06_cuboctahedron_147_24-1252-E | CACCAACCTACTCATCTTTGACCCCCAGCGACACTACGAAGG | Core staple |
| 06_cuboctahedron_147_24-2669-E | AGGTAAATATAAAATTCATATGGTTTACCAGTGAGGGAGGGA | Core staple |
| 06_cuboctahedron_147_24-2648-E | ATTCATTAAATTATTTTGTCACAATCAATAGTGACGGAAATT | Core staple |
| 06_cuboctahedron_147_24-2638-E | GGTGAATTATCGGAATAAGT | Core staple |
| 06_cuboctahedron_147_5-4128-Vertex-3T-R1 | TTCCCTTAGAATTTTTTCCTTGAAAAAATCGCAAGACTTTTTAAAGAACGCGTTTCCTCCTCCTCCTCCTCCT | Vertex 1a docking strand |
| 06_cuboctahedron_147_21-3834-Vertex-3T-R1 | AATTACTAGAATTTTTAAAGCCTGTTATATAAAGTACTTTTTCGACAAAAGGTTTCCTCCTCCTCCTCCTCCT | Vertex 1b docking strand |
| 06_cuboctahedron_147_17-2344-Vertex-3T-R1 | CCGCCACCCTCTTTTTAGAGCCACCAAGAAGGCTTATTTTTTCCGGTATTCTTTTCCTCCTCCTCCTCCTCCT | Vertex 2a docking strand |
| 06_cuboctahedron_147_20-3246-Vertex-3T-R1 | TAACGAGCGTCTTTTTTTTCCAGAGCCGTCAGACTGTTTTTTAGCGCGTTTTTTTCCTCCTCCTCCTCCTCCT | Vertex 2b docking strand |
| 06_cuboctahedron_147_10-3099-Vertex-3T-R1 | AGCGCATTAGATTTTTCGGGAGAATTTAAGAAAAGTATTTTTAGCAGATAGCTTTCCTCCTCCTCCTCCTCCT | Vertex 3a docking strand |
| 06_cuboctahedron_147_11-2785-Vertex-3T-R1 | AAAAGAAACGCTTTTTAAAGACACCACACCGTCACCGTTTTTACTTGAGCCATTTCCTCCTCCTCCTCCTCCT | Vertex 3b docking strand |
| 06_cuboctahedron_147_9-5304-Vertex-3T-R1 | TGTAGCAATACTTTTTTTTCTTTGATTTGAAATGGATTTTTTTATTTACATTGTTTCCTCCTCCTCCTCCTCCT | Vertex 4a docking strand |
| 06_cuboctahedron_147_22-5010-Vertex-3T-R1 | ATTAGTCTTTATTTTTATGCGCGAACGCATCACCTTGTTTTTCTGAACCTCATTTCCTCCTCCTCCTCCTCCT | Vertex 4b docking strand |
| 06_cuboctahedron_147_1-1913-Vertex-3T-R1 | ATCACCGTACTTTTTTCAGGAGGTTTTAAAGATTCAATTTTTAAGGGTGAGATTTCCTCCTCCTCCTCCTCCT | Vertex 5a docking strand |
| 06_cuboctahedron_147_3-5755-Vertex-3T-R1 | AGGGCGAAAAATTTTTCCGTCTATCATAGATTTTCAGTTTTTGTTTAACGTCTTTCCTCCTCCTCCTCCTCCT | Vertex 5b docking strand |
| 06_cuboctahedron_147_2-1609-Vertex-3T-R1 | ACTAAAGGAATTTTTTTGCGAATAATGTTTAATTTCATTTTTACTTTAATCATTTCCTCCTCCTCCTCCTCCT | Vertex 6a docking strand |
| 06_cuboctahedron_147_13-139-Vertex-3T-R1 | ATTCCCAATTCTTTTTTGCGAACGAGAACTACAACGCTTTTTCTGTAGCATTTTTCCTCCTCCTCCTCCTCCT | Vertex 6b docking strand |
| 06_cuboctahedron_147_7-286-Vertex-3T-R1 | AGAGTACCTTTTTTTTAATTGCTCCTTTGAGATTTAGTTTTTGAATACCACATTTCCTCCTCCTCCTCCTCCT | Vertex 7a docking strand |
| 06_cuboctahedron_147_23-580-Vertex-3TR1 | CAAAAGAAGTTTTTTTTTGCCAGAGGTCAAAAATCAGTTTTTGTCTTTACCCTTTCCTCCTCCTCCTCCTCCT | Vertex 7b docking strand |
| 06_cuboctahedron_147_4-5902-Vertex-3T-R1 | GCCCCAGCAGGTTTTTCGAAAATCCTGGAGCAAACAATTTTTGAGAATCGATTTTCCTCCTCCTCCTCCTCCT | Vertex 8a docking strand |
| 06_cuboctahedron_147_7-6764-Vertex-3T-R1 | TGTTAAAATTCTTTTTGCATTAAATTGGCCAACGCGCTTTTTGGGGAGAGGCTTTCCTCCTCCTCCTCCTCCT | Vertex 8b docking strand |
| 06_cuboctahedron_147_4-5451-Vertex-3T-R1 | TTCCTCGTTAGTTTTTAATCAGAGCGACATTTGAGGATTTTTTTTAGAAGTATTTCCTCCTCCTCCTCCTCCT | Vertex 9a docking strand |
| 06_cuboctahedron_147_15-4569-Vertex-3T-R1 | GGAATTATCATTTTTTCATATTCCTGCGTGGCGAGAATTTTTAGGAAGGGAATTTCCTCCTCCTCCTCCTCCT | Vertex 9b docking strand |
| 06_cuboctahedron_147_2-2060-Vertex-3T-R1 | ATTTCGGAACCTTTTTTATTATTCTGAGAAGATGATGTTTTTAAACAAACATTTTCCTCCTCCTCCTCCTCCT | Vertex 10a docking strand |
| 06_cuboctahedron_147_5-3530-Vertex-3T-R1 | TAGAAACCAATTTTTTCAATAATCGGGCGCAGTCTCTTTTTTGAATTTACCGTTTCCTCCTCCTCCTCCTCCT | Vertex 10b docking strand |
| 06_cuboctahedron_147_6-1462-Vertex-3T-R1 | GACAACAACCATTTTTTCGCCCACGCTTCATGAGGAATTTTTGTTTCCATTATTTCCTCCTCCTCCTCCTCCT | Vertex 11a docking strand |
| 06_cuboctahedron_147_24-1168-Vertex-3T-R1 | GTACAACGGAGTTTTTATTTGTATCACATAGGCTGGCTTTTTTGACCTTCATTTTCCTCCTCCTCCTCCTCCT | Vertex 11b docking strand |
| 06_cuboctahedron_147_8-6460-Vertex-3T-R1 | AACCAGGCAAATTTTTGCGCCATTCGTTCCCAGTCACTTTTTGACGTTGTAATTTCCTCCTCCTCCTCCTCCT | Vertex 12a docking strand |
| 06_cuboctahedron_147_19-6186-Vertex-3T-R1 | CTCACAATTCCTTTTTACACAACATAGGGAACAAACGTTTTTGCGGATTGACTTTCCTCCTCCTCCTCCTCCT | Vertex 12b docking strand |
| 06_cuboctahedron_147_21-3884-Edge-3T-R1 | CCGACCGTACCTAAATTTAATGGTGGCGGTCATTTCCTCCTCCTCCTCCTCCT | Edge docking strand |
| 06_cuboctahedron_147_20-3296-Edge-3T-R1 | AGTTGCTAGTTTTGAAGCCTTAAAGAATCGCCTTTCCTCCTCCTCCTCCTCCT | Edge docking strand |
| 06_cuboctahedron_147_10-3172-Edge-3T-R1 | AGAAACGATTCACCAATGAAACCATCGATAGCAATCCAAATATTTCCTCCTCCTCCTCCTCCT | Edge docking strand |
| 06_cuboctahedron_147_22-5060-Edge-3T-R1 | TGACCTGATGGCCAACAGAGATAGTGAGTTAATTTCCTCCTCCTCCTCCTCCT | Edge docking strand |
| 06_cuboctahedron_147_3-6974-Edge-3T-R1 | ATAAATTAATGTTGAGTGTTGTTCCAGTTTGCGTTCTAGCTGTTTCCTCCTCCTCCTCCTCCT | Edge docking strand |
| 06_cuboctahedron_147_12-1839-Edge-3T-R1 | CACCCTCATTCATTTGGGGCGCGAGCTGAAACTCAGAGCCACTTTCCTCCTCCTCCTCCTCCT | Edge docking strand |
| 06_cuboctahedron_147_13-790-Edge-3T-R1 | GTTGGGAAGACAACATGTTTTAAATATGCAACCAGTCAGGACTTTCCTCCTCCTCCTCCTCCT | Edge docking strand |

| | | |
|---|---|---|
| 06_cuboctahedron_147_7-6814-Edge-3T-R1 | AAACAGGAGGTTGATAATCAGAAAATATCGCGTTTCCTCCTCCTCCTCCT | Edge docking strand |
| biotin-06_cuboctahedron_147_21-3918-Edge | /5Biosg/TCATCTTCTGGTGATAAATAAGGCGTTAAATTTTAGTTAATT | Biotinylated staple |
| biotin-06_cuboctahedron_147_21-4894-Edge | /5Biosg/CAGTGCCACGCCGAACGAACCACCAGCAGAAACCGCCTGCAA | Biotinylated staple |
| biotin-06_cuboctahedron_147_22-5094-Edge | /5Biosg/AGGGACATTCAAGCGTAAGAATACGTGGCACCCAGTAATAAA | Biotinylated staple |
| biotin-06_cuboctahedron_147_22-2983-Edge | /5Biosg/AAACAATGAAATTGAGCGCTAATATCAGAGAATAAGAGCAAG | Biotinylated staple |
| biotin-06_cuboctahedron_147_10-3130-Edge | /5Biosg/CTTTACAGAGAATCACCAGTAGCACCATTACAAAATAGCAGC | Biotinylated staple |
| biotin-06_cuboctahedron_147_10-3193-Edge | /5Biosg/TTATTTATCCCAGCACCGTAATCAGTAGCGAAAACAGCCATA | Biotinylated staple |
| biotin-06_cuboctahedron_147_20-3330-Edge | /5Biosg/CTTGCGGGAGTTTTGCACCCAGCTACAATTTGAACCTCCCGA | Biotinylated staple |
| biotin-06_cuboctahedron_147_20-3697-Edge | /5Biosg/TTCGAGCCAGTGCGTTATACAAATTCTTACCGCAGAGGCATT | Biotinylated staple |

**Table S2** **3D wireframe cuboctahedron DNA origami strands.** Scaffold is MP13mp18. To make a pattern on a 3D wireframe cuboctahedron DNA origami, we mix scaffold strands, biotinylated strands, and core staple strands (positions corresponding to the pattern docking positions and biotin positions should be excluded beforehand) along with the corresponding docking strands that make up the pattern.

| Letters | Binary | Letters | Binary |
|---|---|---|---|
| A | 1000001 | T | 1010100 |
| B | 1000010 | U | 1010101 |
| C | 1000011 | V | 1010110 |
| D | 1000100 | W | 1010111 |
| E | 1000101 | X | 1011000 |
| F | 1000110 | Y | 1011001 |
| G | 1000111 | Z | 1011010 |
| H | 1001000 | Space | 100000 |
| I | 1001001 | 1 | 110001 |
| J | 1001010 | 2 | 110010 |
| K | 1001011 | 3 | 110011 |
| L | 1001100 | 4 | 110100 |
| M | 1001101 | 5 | 110101 |
| N | 1001110 | 6 | 110110 |
| O | 1001111 | 7 | 110111 |
| P | 1010000 | 8 | 111000 |
| Q | 1010001 | 9 | 111001 |
| R | 1010010 | 0 | 110000 |
| S | 1010011 | | |

**Table S3 Letters to binary and number to binary.** In our demonstration, the last six digits of the binary encoding are assigned to the alphabet, while the last four digits are allocated to the numbers.

| Strands | Concentration |
|---|---|
| M13mp18 (for 2D RRO and Cuboctahedron) or P8064 (for Tunnel) | 20 nM |
| Core staple (positions corresponding to the pattern docking positions and biotin positions should be excluded beforehand) | 200 nM/strand |
| Biotinylated staple | 1000 nM/strand |
| Corresponding docking strands | 1250 nM/strand |
| TAE $MgCl_2$ buffer | 1× |

**Table S4 Mixing concentrations for 2D RRO, cuboctahedron, and Tunnel origami experiments.**

| **Imaging Parameters** | **NSF 2D dataset** | **20 nm RRO 32 nt binder** | **20 nm RRO 64 nt binder** | **14 nm RRO 64 nt binder** | **10 nm RRO 64 nt binder** | **ASU one-redundancy 2D dataset** | **ASU two-redundancy 2D dataset** | **0407 3D dataset** |
|---|---|---|---|---|---|---|---|---|
| DNA origami concentration | 1 nm, no fiduciary drift correction markers | 1 nm, no fiduciary drift correction markers | 1 nM, no fiduciary drift correction markers | 1 nM, no fiduciary drift correction markers | 1 nm, no fiduciary drift correction markers | 1 nm, no fiduciary drift correction markers | 1 nm, no fiduciary drift correction markers | 1.5 nM with 0.5 nM of 20 nm RRO for fiduciary drift correction markers |
| Imager concentration | 5 nM | 5 nM | 5 nM | 5 nM | 2 nM | 1 nM | 1 nM | 1 nM |
| PCA, PCD, Trolox concentration | 1.25X PCA, 1× PCD and 1× Trolox | 1.25X PCA, 1× PCD and 1× Trolox | 1.25X PCA, 1× PCD and 1× Trolox | 1.25X PCA, 1× PCD and 1× Trolox | 1.25X PCA, 1× PCD and 1× Trolox | 1.25X PCA, 1× PCD and 1× Trolox | 1.25X PCA, 1× PCD and 1× Trolox | 1.25X PCA, 1× PCD and 1× Trolox |
| Camera exposure time | 50 ms | 50 ms | 50 ms | 50 ms | 50 ms | 50 ms | 50 ms | 50 ms |
| Laser power density | 800 $W/cm^2$ | 800 $W/cm^2$ | 800 $W/cm^2$ | 800 $W/cm^2$ | 1250 $W/cm^2$ | 1250 $W/cm^2$ | 1250 $W/cm^2$ | 1250 $W/cm^2$ |
| No. of frames | 15,000 | 15,000 | 15,000 | 30,000 | 90,000 | 30,000 | 30,000 | 43,510 |
| TIRF | Yes | Yes | Yes | Yes | Yes | Yes | Yes | Yes |
| 3D lens | No | No | No | No | No | No | No | Yes |

**Table S5 DNA-PAINT super-resolution imaging parameters for each experiment**

| Name | Sequence | Note |
|---|---|---|
| 5[32]25[31] | ATATCTATTATCTGGTCAGTTGGCTTATCTAATCTTTCCTTACCGCAC | Core staple |
| 12[55]28[32] | AGAAATAATAGATTTTATATTATTTATCCAGCGCATTAGA | Core staple |
| 12[183]29[191] | ATTCGCCAGCAACTGTCGCCACCCCACCCTCAGAGCCCAT | Core staple |
| 0[231]21[231] | TCTGGCCTAGCTTTCACAGGTCAGTACCTTTA | Core staple |
| 6[183]2[184] | AGTTTTAAGACGATAATCTGGTCACAACCAGCTTACGGCTATGCCGGG | Core staple |
| 22[95]3[79] | AAAAACAGCTTGATACCGATACTTAGCGGGTT | Core staple |
| 3[80]2[80] | GAGTGTTGTTCTCCGAGTGGTCAGTTTGGAAC | Core staple |
| 19[192]30[184] | ACAACATTGTTTCATTTGACAGGATTATTCTGAAAGCCAC | Core staple |
| 15[8]28[0] | ATATTCCCCAGAAGAGCTATCGCAAGAAACAATGAA | Core staple |
| 3[24]31[31] | GCCATTGCTGGATTATGAACGCGAAGGGCTTAGAACAAAG | Core staple |
| 6[231]24[208] | ATAGGTCACGTTGGTGGGAGCAAAGAGCGGAATCGTCAT | Core staple |
| 7[48]5[51] | AAATTAAACGCCACCCTCAATCAATAGTCTTTAATG | Core staple |
| 1[216]0[200] | TCTTAGCCTCCTGTTGCTCGTCATAAACATC | Core staple |
| 7[248]24[232] | GATTGTAATCAGAAAAGCTCAGGTCTTATTATAGTCACAGTT | Core staple |
| 10[235]9[252] | TTATAATCATATGTACCCCGGTTGA | Core staple |
| 6[247]3[250] | GCAAATATGCAAAGCGTTTTGTTATAAATTTTTGT | Core staple |
| 3[56]19[55] | ACCTTCTACCCTACTGCGGGATCTTACCAGTATAAAGAAAAAGC | Core staple |
| 23[28]22[5] | TAATATCCGGTATTCTCCCATCCTAATTTA | Core staple |
| 25[8]7[15] | GGGTATTACTAATAAGGAATT | Core staple |
| 2[79]23[71] | AAGAGACAGAGATAGAGACCTGAAAAATCAAGCTATTTTG | Core staple |
| 22[207]21[223] | GCGGATTACCAGCCGGGTCACTGTTGAGTAAGAGCGCCCTAAGAGAG | Core staple |
| 12[79]11[63] | ATTCATTTCAACATATCAAAGACACCACGGTCTTTCCAGTAACAAA | Core staple |
| 12[215]27[215] | AAAAAAGGGTGAGAATAGGATTAGCGGGTG | Core staple |
| 20[159]21[135] | CCAGTCAGGAGCTTGCCCTGACGAGAAGGCAGAAAGAAC | Core staple |
| 1[80]19[95] | TATTAAAGAACCGGTCGCAAGGTGTATTCGGT | Core staple |
| 22[250]21[250] | TGCATCAAAAAAAGCCCGAAAG | Core staple |
| 21[40]4[32] | AGCGAACCCAGATATAAAACGCTCTTTTGAATGGCCAGAA | Core staple |
| 9[136]24[144] | GGTTGGCCGTTCCGGCATTCCACATTTCGCCAAGTACGCT | Core staple |
| 30[79]0[80] | ACCAAAAGTACCCGACTTGAGCCACAACCATCAACCGATAGACTCCAA | Core staple |
| 17[96]15[95] | CGAGGGTATTCATCTTCTGACCTAACGCGAGA | Core staple |
| 26[159]12[144] | CACCCTCCACAGGCTTACCAGTCCCGGAA | Core staple |
| 10[31]27[23] | GAGGATTTAGAAGTATTTAAATCCAATTGAGCTGAGTTAA | Core staple |
| 28[215]12[184] | AGTACTCCTCAAGAGAAGCCACCAGCCGGATAGGCCGGAGACAGGCC | Core staple |
| 23[208]22[208] | TCATTGAATCCCCCTTAAGAGGTCATTTTT | Core staple |
| 28[103]13[103] | ACAAAGGGCGACATTCATGCTGATGCAAAAAC | Core staple |
| 16[191]14[160] | CAATGTGCTGCAAGGCGATTTCAGAGGTGGAGTGCCATCTCTCACCGG | Core staple |
| 13[136]12[120] | AACTGACCTTTGTGAGAGATAGACTTTCTCCG | Core staple |
| 22[183]18[184] | CAACACTAAATGCAGATACATAACGATTCATCGCCAGCATCCAAGGGT | Core staple |
| 25[192]9[207] | TGTTTAGATACCAGGCCAGAATTAATGCCGGA | Core staple |
| 15[80]18[80] | TTGCTAGAAATTTAATGGTTTGAACAGCAGCGAAAGACAGGGGAGTTA | Core staple |
| 27[96]10[96] | GACAAGCCTCTGTTATGTTGGCACGGAAAAATCGGTCTGAGAGACT | Core staple |
| 6[151]22[144] | TTTCCCGTTCAACTTTAATCATTTTATGCGATTGTAAA | Core staple |
| 24[55]7[47] | TCTTACGTTTTTATTTTCATCCTGAATAACCTCAAATATC | Core staple |
| 8[95]12[80] | TGACCTTTAATTAATTCATATGGTCGGCTTAGATAACTATATGGAATT | Core staple |
| 27[216]12[216] | CCTTGAGTAACAGGCTTAATCAACGCAAGGAT | Core staple |
| 8[63]10[56] | AGAGGTGTATTAACACCTACATTTAATGCCTGCAACA | Core staple |
| 7[16]4[5] | GAGGAAGGAAATCAACGAAACCAACCGTTGCCTGAGTAGAAG | Core staple |
| 25[64]24[80] | AGCGAATAAGTTTATTTTGTCACATTGCTTTC | Core staple |
| 10[183]27[191] | CGCTTCTGCCAGGCAAGCCGTCGAGAACCGCCTCCCTCAG | Core staple |
| 15[160]28[168] | CGGGAACGGATCAGCTTACGCAACTTTGCCACTCAGACAT | Core staple |
| 13[144]17[159] | AACTTTGATGAGTTTCCACCGTAACAGAATACCGGATATTCACGG | Core staple |
| 23[5]6[8] | CGAGCATGTAAGTTGAA | Core staple |
| 14[23]16[3] | AGAAACCATGATTATCGTACCGACAAAAGGTAAAGT | Core staple |
| 24[231]26[216] | CAGAAAACCAAGAGAAGTAATCGTAAATTTTGGCTATACTTAACGGGG | Core staple |
| 30[183]16[192] | CAGAACCAGTTGGGTAACGCCTATAACAGTTGCAAATGGT | Core staple |
| 14[159]15[159] | AAACAGGACAGATGAGACCAGGCGCATCCA | Core staple |
| 3[120]24[128] | GCGCCAGGGTGGTCGTGAGGCGAAGAATTATGTTCAACAG | Core staple |
| 19[144]19[127] | ATAAGCGTGTTTTCACGGTCATACCGGGATTGCCCTTCACAACACTCA | Core staple |
| 1[24]18[24] | TGGCAGATGAGTAAAAAATCGCCATATTTAACTGTAATTTAGGACAAC | Core staple |
| 12[31]29[23] | GGTTAGCCCGAACGTTATTTTGCGTAATAAGATTAGAGAG | Core staple |
| 24[127]9[135] | TTTCAGCGGTAAATGAATTTTCTGGAGCCACCAGTTGGGC | Core staple |
| 18[55]2[40] | AATCATAATTACAACAAACGCCTAGCCAACGCCACACGACGCTCAATC | Core staple |
| 27[32]10[32] | AACTGAACAATGGAAGTACCATATCAAAATTACTGAGAGCCAGCATTT | Core staple |
| 28[167]13[167] | AATCAAAATCACCACAAGAATCGGCGAAAC | Core staple |
| 9[80]8[64] | TTCATTTGTTTAATGGAAACAGAAGATAAAAC | Core staple |
| 27[112]14[120] | TTAGCCAGGGATAGCAACAACGCCAATCATAACGACCTGC | Core staple |
| 23[104]5[103] | GCCTTAGAAAGGAACGGGGAGAGGCGGTCCCTTATAAATTAGAATC | Core staple |
| 11[64]9[79] | AGAAGATGATGAAACAAAACAAAATTAACAAT | Core staple |
| 14[119]15[135] | TCCATGTTTATTTGTATCATCGCCTGATAAAT | Core staple |
| 9[208]25[223] | GAGGGTAAGAGATCCGTCCAATACTGAAT | Core staple |

| | | |
|---|---|---|
| 16[119]31[111] | GAGATTTTAGTTAATGCAACGGAATTATTAGCAAAA | Core staple |
| 19[168]3[167] | ACGTTAATTTTAGGAATGTCACTGAGCCAGCGGTGCCGGTGTGGTGCC | Core staple |
| 27[160]9[175] | ACCAGAGCCGCCATGTGGCCTTTAGTGGGAAAGTGCCATGTTTCGTCT | Core staple |
| 11[192]26[208] | TCACCATCAATATGATATTCGGGTCAGGTT | Core staple |
| 17[8]30[3] | TGTCCAGAAGCCCTTTTTA | Core staple |
| 0[39]21[39] | AGGCCACCTCACCAGTTCAACAGTGGCGTTTT | Core staple |
| 31[32]15[31] | TTACCAGAATGAAAATAGCAGCCTAATAACATAATATAAAAGATGATG | Core staple |
| 31[112]1[119] | TCACCAGTCACAGGAAGTTTCCATTTGCCCCAGCAGAG | Core staple |
| 9[48]25[63] | CTAGTCAGAGGCGAAGAGCCCAACGCTAACG | Core staple |
| 10[135]26[112] | ATCGACATGGATCAAACTTAAATTGAGACGCATTTGTAAC | Core staple |
| 28[191]15[183] | CTCAGAACTGGGAAGGCGGGCCTCTTCGCTATGGCGAAAG | Core staple |
| 3[168]5[175] | ATCCCACGGCAGCAACCGCAAGAAATGACTTGTAGAACGT | Core staple |
| 14[87]28[72] | CAATTGAATACCAAGTCTTATTACAGCAAACG | Core staple |
| 11[0]10[2] | ATTCGACAACTCGTATAGACTTTACAAA | Core staple |
| 30[95]14[88] | GTGAATTATTGAGGGAGGGAAGGTCGGTCCAATCGCAAGA | Core staple |
| 6[207]4[208] | GCGCATCGGCACTCAATCCGCCGGGCAACGGGAACAGCGGTTGCGG | Core staple |
| 13[104]9[103] | GAGGCGCGGATAGCCTCATAGGATCTAAAGTTTATTTATCAAAA | Core staple |
| 13[216]14[224] | TTTTAGAACCCTCGCAAAATTAAGCAATAGCAAGGCAAAGAATTAATATA | Core staple |
| 1[96]17[95] | CACGCTGGTTAAACGGGTAAACAATTTGGGAAGGCTTGCACATCGGAA | Core staple |
| 27[0]12[0] | ATAACCCATTAAAAGT | Core staple |
| 10[215]10[184] | TGCAACCGTTCTAGCTGATACTTTCCGGCAC | Core staple |
| 12[244]27[236] | AAGCCTCAGAGCATTTGA | Core staple |
| 17[64]14[56] | TCACCCTATACCGACAAGACTCTACCAGATGAATATA | Core staple |
| 19[56]30[64] | CTGTTTAGTATCATATTTTGCGTAACGGAAAACTGGC | Core staple |
| 6[95]4[72] | GCTATATGTGAGTGAAAATTTCTTATAGCCCGAGATAGTA | Core staple |
| 29[0]13[7] | ATAGCAATGAGCGGAATTGAGTAA | Core staple |
| 3[5]3[23] | TAGTAATAACATCACCCA | Core staple |
| 27[237]29[244] | TGATACAGGTAAGTTCCA | Core staple |
| 29[192]13[215] | GAAAGTATTGTCGGTGGCGATGTAGGTAAAGATTCAAAT | Core staple |
| 17[24]14[24] | AATAACAGAGGCATTTAATAAGAGAAAAACAGTAATCCTGATTGTCAA | Core staple |
| 6[127]8[120] | TTTCCAGTAATGAGTGAGCTAACTGAGCCGGAAGCATAAA | Core staple |
| 18[111]0[96] | AGGCTTTGACTTTTTCATGTAACGCCTGGCCCTGAGAGAGTTGCA | Core staple |
| 5[152]4[136] | CGTAATCTGCCAACGGCCACAGTTGAGGAT | Core staple |
| 15[184]16[208] | GGGGATAACCTGTTTAGCTATATTTTCATTTATTAGAT | Core staple |
| 23[72]24[56] | CACCCGTGTAACCTTGCTTCTCCTAAAACATAATACCGTCCTGAA | Core staple |
| 31[240]0[232] | ACAAATAATCAAATATTTCGAGCTTCAAAAATAATTCGCG | Core staple |
| 6[159]26[160] | ATAGCTGGAAATTGGAGGTTTCCCTCAGAACAGTATATATACGC | Core staple |
| 16[239]29[231] | AACATCCAAGGTGTTAGTCTCTGA | Core staple |
| 5[52]3[55] | CGCGAACTGATTGGCACAGACAATATATGGAAAT | Core staple |
| 30[207]17[223] | GGAACCTAGGTTGAGGCAGGTCAGACGATTGCAACTAAAAACGAGTA | Core staple |
| 10[55]8[40] | GTGCCACGTTTGCACGAGCCTAATTTGCCCTGAACAAGCACATCACCT | Core staple |
| 22[143]21[159] | TTGGGCTTCCGTGAGCCTCCTAGCACCGTCGGCCCCCTAGAACTG | Core staple |
| 5[104]6[128] | CTTGAGTCGTGCCAGCTGCATTAATGAACGGCTGCCCGC | Core staple |
| 1[5]2[5] | AATTAACCGTCTTCTTTGAT | Core staple |
| 19[208]30[224] | GCTCAACATGTTTTAAATGCAAACGGAAATGGCCTTGATGAATGGAA | Core staple |
| 21[5]1[23] | ACAAGAAAAATAATATAATTACAATATGTAGCAT | Core staple |
| 4[135]22[120] | CCCTCGGCCAACGCGCAACTAAAGTAATAATT | Core staple |
| 0[167]22[168] | AGAATGCGCAGCGCAGTACTTATAGCTCACACATTCAACTTCATAACC | Core staple |
| 27[56]12[56] | ACAACATAAAGGTGGCAATTACCTGAGAAC | Core staple |
| 23[56]1[55] | AATGTTGATTAAGCAAGCAAATTCCCGACTATTTTGACCAGTAATA | Core staple |
| 3[233]5[237] | GGCGCATAAATTATTCTCCGTGGCGGATTGACCGTAA | Core staple |
| 10[159]27[159] | AGTTAAACGATGCTGAAAAGCCGAGAACCGCATGTACCGTAACGGA | Core staple |
| 24[183]6[160] | GAGGGGGTGTATCACCTACCAGACCGGAACGTGCCGGGTC | Core staple |
| 30[119]29[135] | CCGGAGGACTAATACCAAGCGCGAAACAATCTAGAGTAAAAAACCATC | Core staple |
| 2[103]3[119] | GTTTGATGAATCGGCAAAATTTGCGTATTGG | Core staple |
| 18[79]21[79] | AAGGCCGCTGCCGCATGCCCAGTTATACAAATGGTTTTGAAGCGTTGC | Core staple |
| 15[32]29[55] | GCAATTCATCATTTTAGTACCATAGGACAATCCAAATAAG | Core staple |
| 4[207]6[184] | TATGAGCATCAGCGGGGCGCTTTCTAACCGTGCATCTGCC | Core staple |
| 25[96]8[96] | AAATGTCGTCTTTCCCCGAAGAGTCAATAG | Core staple |
| 25[224]10[216] | GACCATAAATAAGTTTAGCATGTCGACCCTGTAATACTTT | Core staple |
| 0[199]2[200] | CCTTACACAGCAAATCGTTTGGGTGGTAAAAC | Core staple |
| 8[39]11[31] | TGCTGATCTTTAGGAGTAGATAATCAGAGGGTTTTGAACC | Core staple |
| 30[23]17[7] | TTACACCGACGACGACATGTTCAGCTAATGCAGAACAATTC | Core staple |
| 17[224]31[239] | GATGCATCAATTCTACTAAAGCCAATTCACAA | Core staple |
| 25[32]9[47] | TCATCGAGAACAAAGTACAGCAAATGAAAAAT | Core staple |
| 11[216]10[236] | AGAAGCCTTTATTAGCTAAATCGGAACA | Core staple |
| 13[40]27[55] | CTGTTGCGAGAAAAATACCAGTTACAAAATAA | Core staple |
| 21[200]1[215] | GAATGGCTTAGAGCTTGCGGCTAAAGGTT | Core staple |
| 26[207]6[208] | TTGCTCAGCTGGATAGTACAAAGGATTGCCTGAGAGTCTTAGATGG | Core staple |
| 12[119]16[120] | TGGTGAAGAGACGGTCTGTAGCATGACAACGTCACCAATGGTACAACG | Core staple |
| 16[207]19[207] | ACATTTCGATTCCCAATTCTGCGGTACGGTGTCTGGAAATTCTGTA | Core staple |

| | | |
|---|---|---|
| 21[136]2[128] | ACCAGAACGAGTATTAGCAGCGTGCCTGTTCTTCGTTTTC | Core staple |
| 28[71]13[71] | TAGAAAATACATGCCCAGGTTTAACGTAAA | Core staple |
| 24[143]5[151] | AAACAACTGTTTAATTTGCGCTCAGTACCGAGCTCGAATT | Core staple |
| 13[72]30[80] | ATCGCGCAGAGGCTAAAACATGTTGCAGTCGATCACCGTC | Core staple |
| 24[79]7[79] | GAGAGCTACAATTTTAAACGAACCACCAGCAG | Core staple |
| 31[176]0[168] | GAGCCGCCAGTTGAGAAAAACGAACTGTGGTGCTGCGGCC | Core staple |
| 27[136]7[151] | AGGAACCCCACCCTCATATGGGATCAACATACCACATTAATTGTGTGT | Core staple |
| 14[55]16[40] | CAGTAACACATCGGGATAAATAAGGCGCCAGT | Core staple |
| 22[119]2[104] | TTTTCACGGGCACCAAAGTGGCGAAAATCCT | Core staple |
| 13[8]26[0] | CATTATCATTAATTCAAGAATGCTAATATCAGAGAG | Core staple |
| 29[56]15[79] | AAACGATTCGTGTGAGAAACAATAACGGATTCGCCTGA | Core staple |
| 0[79]1[79] | CGTCAAAGGGCGAAAACATTCTGGCCATCCAC | Core staple |
| 13[168]25[183] | GTAGCTGCTTCAGCAGCACCACCGGAGGGTTGAGCCCGGAATAGGTAA | Core staple |
| 30[63]17[63] | ATGATTTTTTGTTTAAAATAAGAATAAACTCG | Core staple |
| 8[159]10[160] | GCTCACAAAACGCGGTCCGTTTAAGGGTAA | Core staple |
| 2[183]19[191] | TTACCTGCCGCGCCTGTGCTGTTCTGGTGACTCTAACGGA | Core staple |
| 29[136]27[135] | GATAGCAGGTCACCAGTACAAACTAGCCCAAT | Core staple |
| 4[71]5[87] | AGAATACGAGCGTAAATCGTCGCTATTAATTA | Core staple |
| 19[128]18[112] | TCTGCTCAAAGCTTTGACCCCCAGCGATTCAG | Core staple |
| 30[223]15[232] | AGCCCTGCTGCCCGCAGTTTGACCGGGGCGCGAGCTGAAAA | Core staple |
| 21[80]22[96] | GCCGACAATGAATACGTAATGCCACTACGAATTGAAAATCTCCAAA | Core staple |
| 22[167]7[175] | CTCTTACCGTGAAGTTGTACTCAGTTACCAGAGCACATCC | Core staple |
| 15[96]28[104] | AAACTTTTTCAAATAACTTAGCAAATATTTCCACAG | Core staple |
| 30[159]16[144] | AGCTAGCGATCAGGTTCCGAGGCTGGCTGAC | Core staple |
| 14[223]30[208] | TTTTAAATGCAATGCCTGAGTAATAAGAGGCTGAGTAACTATTTC | Core staple |
| 20[223]21[199] | GATTATTGCTGAATATAATGACAGGTAGAAAGCCAAAAG | Core staple |
| 5[176]22[184] | CAGCGTGGTGCTGCAGGTCATTGGAAACCAAAAGTAAGAG | Core staple |
| 8[191]8[160] | CGGCCTCAGGAAGCGCTGGCAGCCTCCGGTCC | Core staple |
| 7[80]25[95] | TACATAAATCAATTAGTTATCAGCATCAATAG | Core staple |
| 1[56]0[40] | AAAGGGAACCGTCTATCATTATAATCAGTG | Core staple |
| 31[3]20[5] | AGAAAAGTAAGCAGATAGCCATTATAGATAAGTCCTGA | Core staple |
| 29[232]28[216] | ATTTACCGCGTCATACATGTGCCCGTATAAAC | Core staple |
| 9[120]10[136] | CCGCCAGCACCCTCATGAAACAGCAAAAAAATCCCGTAAAATTTGTAC | Core staple |
| 7[176]25[191] | TCAGAGGGGACGACGATTTTGCCATAGTAAAA | Core staple |
| 15[136]13[135] | TGTGTACAACGGTGTCGAAATCCGGGGAACCG | Core staple |
| 21[232]6[232] | ATTAGAGGGATTAGCTCCTTTTGACAAATGCTTTAAAGAATTAATGGG | Core staple |
| 27[24]13[39] | GCCGAATTCGGGACAAGAATTGGATTATACTT | Core staple |
| 30[247]18[240] | ATCCTCATTAATAGTATCAAAGCG | Core staple |
| 17[160]31[175] | CCAGTGCCAAGCTTAACCATAGCCGGTCACCA | Core staple |
| 19[3]0[8] | GCGCCTGTTTATCAACAGAGGAGTCTGTCCATCAC | Core staple |
| 5[88]23[103] | ATTTTCCCTCAAAAGAAAAGGCTCCAAAAGGA | Core staple |
| 2[199]23[207] | GGCATCAGGGAGGTGTCGAGGCATATAGCGAGAGGCTTAT | Core staple |
| 16[143]30[120] | CTTCATCATGACAAGACAAGTTTGCCTTTCATTAGCAAGG | Core staple |
| 7[112]6[96] | GGGTGCCTCGGGAAACCTAAACATAGCGATA | Core staple |
| 9[16]24[8] | GCCGTCAACACTAACAAAACCAAGTATCATTCCAAGAAC | Core staple |
| 2[39]23[55] | GTCTGAAAAACAGGAAGAAGGCTTCGGGTAGGAATCATTACCGCGCCC | Core staple |
| 17[40]18[56] | TCGAGTTACGTCAAAAAGGAAACCGAGGAAACGCAATAAGGAACCGG | Core staple |
| 24[111]7[111] | ATTAATTGTATCGGTATTAAGACGCTGATG | Core staple |
| 1[120]1[151] | ACGGGCAACAGCTGGGTTTCTGCCAGCACTCA | Core staple |
| 5[13]23[27] | ACTATCGGCCTTGCTGGTACAATATTATCAA | Core staple |
| 9[176]11[191] | CGTATCGCACTCCAGCGGATAAGTAGCTCAAA | Core staple |
| 19[96]30[96] | CGCCTAAAGAGGATGATTAGAGCCCATTAAAG | Core staple |
| 26[239]7[247] | TACTGGTAATCAAAAACCCGAACGTCGATAAAAACAGGAA | Core staple |
| 9[104]24[112] | TCATATGCTCATTTGGTGTAAAGAGACGTTAGAGTGAGA | Core staple |
| 2[127]31[143] | TTTTCACCCCTAAAACAAAGAATAAGCACCATTACAGCGTCAGACTGT | Core staple |
| 24[207]8[192] | AAATTTGCAAAAGAAGCAGAGGTCCTATCTAT | Core staple |
| 10[95]27[95] | ACCTTTTTAACCAAGAAACATCTCTTAAAACGAAAAGCCAGCGCCAAA | Core staple |
| 18[183]30[160] | TTTCCCAGTCACGACGTTGTAAAAGCATTTTCCCCTTATT | Core staple |
| 2[250]1[250] | TAAATCAGCTCCAATAGGAACG | Core staple |
| 18[239]3[232] | AACCAGACCTCTTTAACGCGTCAATCATTAACATTTTACATTAAATGTCAACCCGTC | Core staple |
| 1[152]19[167] | GACGATCGCGGGCCTGGGAAGAAAAATCT | Core staple |
| 31[144]19[143] | AGCGCGTTTTCATCGCGATTACCCAAATCAACGTAACATTCAGTGA | Core staple |
| 13[8]26[0] | CATTATCATTAATTCAAGAATGCTAATATCAGAGAGT/3Bio/ | Biotinylated staple |
| 17[8]30[3] | TGTCCAGAAGCCCTTTTTAT/3Bio/ | Biotinylated staple |
| 31[3]20[5] | AGAAAAGTAAGCAGATAGCCATTATAGATAAGTCCTGAT/3Bio/ | Biotinylated staple |
| 23[28]22[5] | TAATATCCGGTATTCTCCCATCCTAATTTAT/3Bio/ | Biotinylated staple |
| 9[16]24[8] | GCCGTCAACACTAACAAAACCAAGTATCATTCCAAGAACT/3Bio/ | Biotinylated staple |
| 15[8]28[0] | TCCTCCTCCTCCTCCTCCTTATATTCCCCAGAAGAGCTATCGCAAGAAACAATGAAT/3Bio/ | Biotinylated staple and docking |
| 30[223]15[232] | AGCCCTGCTGCCCGCAGTTTGACCGGGGCGCGAGCTGAAAATTCCTCCTCCTCCTCCT | docking |

**Table S6** **DNA origami tunnel strands.** Scaffold is p8064. Core staples are all the strands that form the structure. Biotinylated staples are modified with biotin to immobilize origami on coverslip surfaces through a BSA-Biotin-Streptavidin-Biotin-DNA origami arrangement. To form DNA origami tunnel for DNA-PAINT experiment, we mix scaffold strands, biotinylated strands, and core staple strands (strands at positions corresponding to the docking positions and biotin positions should be excluded beforehand) along with the corresponding docking strands.

| Name | Sequence | Note |
|---|---|---|
| Tetrapod_10[118] | GATAAAAATTAGCAATAGCTAT | Structure |
| Tetrapod_10[65] | GAGTTTATAAATTAGCAAGGGAGGCCGATTATCCTGAGAGAACTCAATA | Structure |
| Tetrapod_11[126] | CTTACCGAATAACCACCAGCAGAA | Structure |
| Tetrapod_11[137] | TTTAATAATACAAAAATGAAAATA | Structure |
| Tetrapod_13[119] | TTTTTGTTTAACGTAGAGCAAAAATGGAGCCCT | Structure |
| Tetrapod_13[147] | GCAGCCTATTGAGTGGCAGATCTTTAATGCGCGAACGCAAA | Structure |
| Tetrapod_13[54] | CTCGTAATTTGCCAGTTCAATAGCCATTGCCTTG | Structure |
| Tetrapod_13[75] | AATAAACAGCCATATTACAACTATCATGGTTTGAT | Structure |
| Tetrapod_15[143] | ATTTAAGCCCTTAAGAAAAGTAAGCA | Structure |
| Tetrapod_15[77] | AAAACGCATAGATTAGTTGGCAAATCAACCACCGCCTGAGTAAAATCA | Structure |
| Tetrapod_16[83] | ACGCTAATCATATTCCTGATTTGAATAAC | Structure |
| Tetrapod_17[144] | TCAATCAAGGAAACAATA | Structure |
| Tetrapod_18[72] | AAAATTATCACGAGCGTCTTTCCAGAGCCTACCAGAAGTATAATCCAGA | Structure |
| Tetrapod_18[83] | CTTGCTTAGACTACCTTTTTCC | Structure |
| Tetrapod_19[91] | GACAAAAGTATGTGAGTAATGGACCTGAATCTTACCA | Structure |
| Tetrapod_22[153] | AACAAACTTACCGCGCCCAATAGCAAGCAAAGAAGTGATTGC | Structure |
| Tetrapod_25[136] | CAAACAGCTTAGTTTCAGCGGAAACATTGAATGAGTTAGAGTCT | Structure |
| Tetrapod_27[126] | TTTATAAAATATCGCCCACGCATAACCGATAGCACATTGCAAGGAG | Structure |
| Tetrapod_27[136] | TATCCTGATACGAGTGAGATCGGTTTTGTAAATTACATAGTGAGAATGAA | Structure |
| Tetrapod_29[119] | TTCGCATCAAAGGACGTTTAGTTCAACGATACC | Structure |
| Tetrapod_29[56] | AGGTTGGAGGTCAAACTAAAATGTCGTCGTACAAA | Structure |
| Tetrapod_29[77] | AGGATGCAGGTGAGACGAAAGTAAATGAACACT | Structure |
| Tetrapod_3[46] | AAGGCTTTGATCAGAGCGGGAGCTAAACAGCCGCGTACTGAT | Structure |
| Tetrapod_3[94] | CAGCATTGACAGGATTTAAAGGGAGCCCCCGATTTAG | Structure |
| Tetrapod_30[149] | TTTCTTAAATCCCCATTCTGCAATGTGCGATTCGAGGTAGAAAG | Structure |
| Tetrapod_31[77] | ACGTTAGAGGCAAATCATGAGGAAGTTTCGAGTTAAAACCA | Structure |
| Tetrapod_32[69] | CGCTATTACGGGATCGGTCAAACAGACCAACAAAG | Structure |
| Tetrapod_33[144] | TGTCTAACGGGTTTAATT | Structure |
| Tetrapod_37[125] | CGCGTTTTAATTCGAAATATAATGAGATGAACAA | Structure |
| Tetrapod_37[91] | AAGATTAAGTACGGCTGACGAGATTTAGTTGAAAGAGGAAATAAGG | Structure |
| Tetrapod_38[153] | AAACGAAACCAAAAACATTATGACCCTGAATCTACCATTGTG | Structure |
| Tetrapod_39[84] | CTTGCCTGCCAATACTGCGGCGGATTGCATCAAA | Structure |
| Tetrapod_41[151] | TTCTCCGATATCATTGATTC | Structure |
| Tetrapod_41[161] | CAGCTTTCATCAACTCATTGTTAGAAGA | Structure |
| Tetrapod_44[80] | ATACATTCGCATTACGCCAGC | Structure |
| Tetrapod_46[185] | ATATCTGCCACATTA | Structure |
| Tetrapod_47[165] | AAATGTGAGTGTGGCGACTGTAGC | Structure |
| Tetrapod_47[45] | TTGTAAAACGAAACGAGG | Structure |
| Tetrapod_47[70] | CTTTATTATCAAACGCCGCGACCAGGAGCGGCCAGGGATGTG | Structure |
| Tetrapod_9[137] | GAAACTGATATTATTTAC | Structure |
| Tetrapod_9[147] | GACACCACGGAATATATTAGTTCACCAG | Structure |
| Tetrapod_0[108] | TTATGGAACCTTTTGGGCCACGCTGGTTTGCACCCTCGCAGGTC | Structure |
| Tetrapod_0[129] | GCTGTGAAAGTAAAAACCAGGCGAAAATCAACCGCCTATTCACA | Structure |
| Tetrapod_5[101] | TAATCCACCAGAGCGGTGTCGAGGTGCCG | Structure |
| Tetrapod_5[112] | GACAGAATCAAGTTGAGCCACCCCCAGCGTCTA | Structure |
| Tetrapod_5[132] | TAGCGTCAGACTGTACCCTCAGCTGTTT | Structure |
| Tetrapod_6[146] | CGTCAAAGGGCGATTAAGA | Structure |
| Tetrapod_8[104] | TTAGAGCCAGCAAAAGAGTCTTACTTCAAATAC | Structure |
| Tetrapod_13[108] | TAAGAAAATATTTTGACCCGTTGTAGCAAGTCCATCAGGCGGTC | Structure |
| Tetrapod_16[111] | AGCTACAATTTTATAACAGTATTTGAAAGATGA | Structure |
| Tetrapod_16[122] | ATTTGCCGTTTATTACAGTACCTTTTACATAGGCAGAACAACAT | Structure |
| Tetrapod_21[94] | CCGTTATAAAGAACGTCTTACCTTTTTATCAGATGAGATTTTC | Structure |
| Tetrapod_21[114] | TAAATTTCGAGCAGTAACATTTAACA | Structure |
| Tetrapod_21[125] | AAACACCGGAATCATTGTAATTTCGGGA | Structure |
| Tetrapod_22[139] | AAAACAAAATTATTATTTT | Structure |
| Tetrapod_23[105] | ATATACCAGTAATATTCTGTCCAGACGATACCGCACTCATCTA | Structure |
| Tetrapod_24[104] | TTTTATGGAGATGATCATTTTTACCGTAATTTT | Structure |
| Tetrapod_29[108] | TGACCTACGAAGGGATTTAGGAACCCATGCAGGGATCGCTGAGG | Structure |
| Tetrapod_32[90] | ACTATCGACATCATAACTGACCAACTGAAAAAGGAATTACGAGGACTGGA | Structure |
| Tetrapod_32[111] | ATCGAACAAGACCCATTCAACCAGTTGAGAAAC | Structure |
| Tetrapod_32[122] | TGCGCCTCAGATACAGGGTAAATTGGGCTTGCTGTAGCGAACGA | Structure |
| Tetrapod_37[105] | GCCCGAAAGACTTCATGTTTTACGAGTATAGAAAG | Structure |
| Tetrapod_39[105] | ACCAGAAAATATGTATAACAGTTGATTCGAATTAG | Structure |
| Tetrapod_40[96] | ATCGGCCTCAGGAGCGGAGGATCTGGAAAATTAACTCTCAGGCACAGATC T | Structure |
| Tetrapod_40[117] | GTTTGAGGGGACGAGTAAACGTCGGAAACCAA | Structure |
| Tetrapod_40[139] | CATCGTAACCGTGCCATCAAACGCCTGAATGAA | Structure |
| Tetrapod_45[101] | GGGAGAGGGTACGTATTCTTGTGCAAGTTGCCATATTCACC | Structure |
| Tetrapod_45[112] | TATTGCTAAACTGGAGCTGATGGGATTGTCGTT | Structure |
| Tetrapod_45[133] | CGAAGTCCGTGAAGAATATGACAGACAGATGAA | Structure |

| | | |
|---|---|---|
| Tetrapod_47[112] | CAGTTCAAAATTAAGAGAGTCTGGAGCAACTCATTTT | Structure |
| Tetrapod_1[164] | TTAGCCCCGGGTACCGAGAGTCACACCACCCTGTCACCATTTCCAG | Structure |
| Tetrapod_1[186] | CAGATATAAGTATAGCCGTATCACGAACCGCGCCTGTAACGA | Structure |
| Tetrapod_14[65] | CCTGAGTAGAAAGTGTTCCAGCAGGTCACGCGAA | Structure |
| Tetrapod_14[83] | TAGTAATGGCCACCGCAACAGACACCCGCCAGCCGCCGC | Structure |
| Tetrapod_15[53] | TTACCGCCAGATAATACACCCTCATATTAATTATCAAAAACT | Structure |
| Tetrapod_18[52] | TTTTGAAACCATTAAATCCTTTGCCACAATTCGATT | Structure |
| Tetrapod_2[59] | CGTGTCCAGTCTGTAAAGCCTGGGGGTTAATTTGTAAATCGT | Structure |
| Tetrapod_2[83] | AGCTTGAGCCAGCTATACGAGCCGGAAGAAATTTAAGGT | Structure |
| Tetrapod_20[58] | ATATCATCTTCTGACCTCATAAAGGGGAAACACGCGCGCCTTC | Structure |
| Tetrapod_20[72] | TGGTTGCGTATGGCAATTCATCAAGAGCGGATCGTCGCATCAATACTGA | Structure |
| Tetrapod_21[39] | ATATATTTTATGCCTAACTCACTGGCGTATTAGAC | Structure |
| Tetrapod_23[39] | ATTATTTGCAGCTGATGGTCAATAAGGAAGGTGGC | Structure |
| Tetrapod_23[52] | AAAACTGATTGTTTGGATGAACAAACCCTTAGACCTCAAAAAA | Structure |
| Tetrapod_23[60] | AATAAAGAAAGTTATATATCATAGAGCCGGCTGCGCGTTACA | Structure |
| Tetrapod_25[66] | ACCGGGAGAATGGTAGGTCGGAAATGATTAGCGTCCACTAAAATCC | Structure |
| Tetrapod_25[88] | CCCTAAATGCACTTCCGCTCGAATACTCCTCACTCCAAGATGGTG | Structure |
| Tetrapod_30[62] | CTACAACCACCCTCACCCTCAGTACTGGGGAAAGCACCG | Structure |
| Tetrapod_30[83] | GAGTTTCCAGAGCCGGCCGCTGGGGTCAAATCCTCGCC | Structure |
| Tetrapod_31[53] | TCTAAAGTTTCACTCATCTTTGAGACACTATGGGGTAATTC | Structure |
| Tetrapod_32[51] | ATCATCGCCTACAAAGTCCCC | Structure |
| Tetrapod_36[38] | TTAAAAAATCAGGTCTTTTTATTAGGAACCAGATCAAAAAGTTTGG | Structure |
| Tetrapod_36[59] | ATTGCTATTATAGTCAGAGCATTTTCCGCCTCAATCGGCATTAAAG | Structure |
| Tetrapod_38[52] | CGCATAGGCTGGAACGAGCTCGTTTAACGGCTAAGA | Structure |
| Tetrapod_38[72] | TGAACGGTGTTCATAAGAAGAGCAGACTAAAGATCGTCAGA | Structure |
| Tetrapod_39[42] | ACCCAAACTCAAATAAGTTTTCTGAATTGATGATATGCT | Structure |
| Tetrapod_39[63] | CTGCTCAATAAATATAGTAAACCAGAATTAATAAGGGA | Structure |
| Tetrapod_4[150] | GCCAGCGCGTTTTCATCGAGCAAAGAATCGTCTTCAGTGCAGA | Structure |
| Tetrapod_4[170] | GAACGGTCATAGCCCCCACCCTGAAATCCCCTCAACGTAGG | Structure |
| Tetrapod_4[65] | GGGCGGAAACGTCACCACATTACCTCAGTGAAACATCAAACAGGA | Structure |
| Tetrapod_4[79] | AATCCCTGAGTAATGAATCGGCCACTGTCGTCGGGGAAGTCTGAGCTGT | Structure |
| Tetrapod_6[160] | AACGTGGAAGAGAATTTTAACTTTGCGGGACTTTTAGAATACTGGGTTC | Structure |
| Tetrapod_6[181] | AACAAGAGGGGTTTCAGGAGTGCAGCGAACAGAGGCTTTGACACAACGG | Structure |
| Tetrapod_7[154] | GTTCCGACCTCAGAATTAAAGATGTTTAGCATAGTGGAACCGTACGCAT | Structure |
| Tetrapod_7[175] | CTTATAAAGCCACCGCAGTCTGCCAGAGCATAACCGCGCAGATTGT | Structure |
| Tetrapod_7[46]_nondocking_mid_arm | GGGCAACAGCTATGGTTTGTAGCGCAAATGAATATCAAATTTGAGGACAA | mid arm position structure |
| Tetrapod_7[60] | TGCGGGAGAGGCGGTTTCCCGCTTGCGAGAAGTGAATTTAA | Structure |
| Tetrapod_7[68] | ACCGCCTGGGCGCCGCAACCACCTGCCACGTCTGGTCAGAGCCGTACAA | Structure |
| Tetrapod_8[86] | CCAGTAGCACATGAAACGCTT | Structure |
| Tetrapod_12[177] | ATAAACAGGGAAGCGCATTTGGGAACAAGTAACACCACCAGCAGC | Structure |
| Tetrapod_14[188] | CGTGGCATAGAAAAACTCCTTTTTAACCTGTAAAGGTGTTTG | Structure |
| Tetrapod_15[182] | GGACATTATTGAGCAACCGAGTTTTTGTATACAGAAAAGTAA | Structure |
| Tetrapod_20[164] | ATCGTAGTATCATATGCGTTTGATTAAACGTGCAAGTATCGGATTGCGA | Structure |
| Tetrapod_23[168] | TTTGAATTTAATTGAGATAAGTTTGTATAGATCAATGTTCGG | Structure |
| Tetrapod_25[193] | CGACAAAACATATAGATTTTTTATACAGTAGGGCACCAAGTCAATTAC | Structure |
| Tetrapod_28[177] | GTTTGAATTTCGCGTCGTTTGAGCTTGGCTAACAAGTCACGAGTTGGG | Structure |
| Tetrapod_30[188] | TTTAATTAATTATTGATGAAGTTTTCCTGAAGCATGTAATATAGATTTTA | Structure |
| Tetrapod_31[182] | ATAATAATCAAACATGTGCAGTTTTTGGTAAATTTGACCATT | Structure |
| Tetrapod_36[163] | GGAGAAGCAAACTCCAATTTGGAGGGACAATCAACAGACAAATAAAAG | Structure |
| Tetrapod_39[168] | AATTACCGTCATTTGGTCAATTTTATTACGCAGAAGGAGCTA | Structure |
| Tetrapod_41[60] | AGGTCCGGCACCGCTTCTTTCGAGGCGAGGCTTATTCA | Structure |
| Tetrapod_44[170] | GCCTTCGCCATCTCATATTTATAAAAAGAGAAGCAGTC | Structure |
| Tetrapod_46[55] | CTGCAAGAACTGTTCATGACGTTTTTTACGACAAGAAACTCA | Structure |
| Tetrapod_47[175] | ATTACAGGGGTGAGCCCCGGTTTTGAGCTGATTAGCTACCTT | Structure |
| Tetrapod_9[193] | TATTTCAACCGATTGAGTTTCAGGTCAATAAGAGTTATGCGACGTTGG | Structure |
| Tetrapod_11[155] | GATAGCCACAAGATTACAGAGAGAATATTAGGTCACGTTGGTGTAGGGCC | Structure |
| Tetrapod_14[167] | GAATGGCAGTTTATTAGCAAACGTAGAAAATATTTACATTTCGCAAATTT<br>GC | Structure |
| Tetrapod_15[161] | TCACACGGATAACCCGAACAATTAATATTTTGTTAAAATTCGTA | Structure |
| Tetrapod_16[176] | GCGAACCTCCCGACTTGCGGTTCCAGCTTGCGATCGGAAAGGGTGCCAAG | Structure |
| Tetrapod_19[143] | CAGAGCCAACAAATTACTAGAAAAATTACTCGCGGGGATTTATTAGA | Structure |
| Tetrapod_22[174] | CTGAGCAAAATCAGGAAACCAATCAATAATCGTTCAGAGCAGGCAATGGG<br>GA | Structure |
| Tetrapod_23[150] | ACGGTAACAACACGCGCCTTATGAAGGTTTATAAGTCTATGAG | Structure |
| Tetrapod_24[168] | ATTACATAACAATCCTCGCTTGCCTGTTCCATATTATTCGCCATGATGA | Structure |
| Tetrapod_27[154] | GAGCAAATGTTTCTAGATAACGCTTGTGTTAGCAGCGTGAGTATTACGGC<br>A | Structure |
| Tetrapod_30[167] | GCTTGCTGAAATGATCGGCATACAAATATTCCTTTGTTTATCAACAATAG<br>A | Structure |
| Tetrapod_31[161] | GAACAACATCTTACAAGATTGTTTGGATGAACGGGAAAGAAACTG | Structure |
| Tetrapod_33[179] | CGGTTTTTAGAATGGGAAGACTCCTGTTTTCTTGTTGGGAGACATCAG | Structure |

| | | |
|---|---|---|
| Tetrapod_35[143] | TGACATTGCTGGCTTCAAAGCGAACTTATTAAAGGTGAATTATCAAAA | Structure |
| Tetrapod_38[174] | GAAGAAATAATACTGCATCAATTCTACTAATATTGTCAATCATATGTAAAAG | Structure |
| Tetrapod_39[150] | TCAAGAGCTTACATTAGATTCATACATAAAGGTGGCAACAGCCC | Structure |
| Tetrapod_40[181] | GATTGACCGTAATGGGATTTACATAAATCAGAGAACCAGTATATTTTT | Structure |
| Tetrapod_41[179] | AATATTTAAATTGTAAACGTTTTAGTTACCAGTATGTTTTGTCAAGGTAA | Structure |
| Tetrapod_42[79] | CGTCCTCTTCAGATCGCACTCCAGTTGAGGTTTTGAAGCCTTAGGAA | Structure |
| Tetrapod_43[151] | AAACACCATCACGGAAACCAGTTTTTATCAAGCACTGCACTGGCGGT | Structure |
| Tetrapod_43[49] | GATTGTGCTGGAACTGCTTGGCGTTATATAGAGCTGATACCCCAAT | Structure |
| Tetrapod_45[46] | AGCAAAATCAAAACTCAACGTTAAAATGCGAGTATTTAAAGGATTTATCA | Structure |
| Tetrapod_46[73] | TGGCGTGCGGGGAAGCGGTTGCTGTCTTTCCTTATCATAATG | Structure |
| Tetrapod_47[157] | AGCAGTCAAATCTAGCATTTGTAGTAGCATTAACATCCAGTT | Structure |
| Tetrapod_8[168] | ATATTGACGGAAATTATTCTTCAGACCGTGGCTTACTTTAATGTTAATA | Structure |
| Tetrapod_0[97] | CAATTCCACACAACGCGCACTAAATCGGAACCCTCCCGCTCA | Structure |
| Tetrapod_2[139] | AGTAACAGTGCCCGTAGGCCTTGAACCCTCATGCCTT | Structure |
| Tetrapod_3[119] | AGACGATTTAAACAGTTGAAACATTTCCTGTGTGAAATTG | Structure |
| Tetrapod_3[140] | AACAAATAGTGCCTTGGGCTGAGTCGTAATCATGGTCATA | Structure |
| Tetrapod_6[90] | TAAAATAGAGTTGCAGCAAACCACCACCAGGCCATCGATAGCAGCACCG | Structure |
| Tetrapod_6[118] | TCAGTTTATTATTCTAATGCCCCCTGCCTAGGTTGAGAGAGCCGCAGTAGC | Structure |
| Tetrapod_8[115] | CCATTTATTAAAAATACCGAACGATACCGTCACCGACTTGAG | Structure |
| Tetrapod_10[97] | AGTATTAAAGTTGAAAAGCACTAATTTATCCCAATCCAAA | Structure |
| Tetrapod_14[125]_nondocking_vertex | AAAACATCGCCAAGCTCAATCGTCTGGAAACAATGAAACTAAACGAT | vertex position structure |
| Tetrapod_15[98] | CTACATCTTTAGGGGAATTGAGGAAGGTTCAGAGGTGACGCAAATTGGGAA | Structure |
| Tetrapod_18[104] | AATCAATAGTAAAGTAAAGAGAAGTGATAAATAAGGCGT | Structure |
| Tetrapod_18[132] | CGGGTATTAAACCAAGCGACAATAAGGCATTAAGAAT | Structure |
| Tetrapod_19[133] | GTTCAGCTTCCAAGAACATCGTAAATCAAGATTAGTTGCT | Structure |
| Tetrapod_22[111] | ATTTCACAGAGAACAAGCAATGCACCC | Structure |
| Tetrapod_23[84] | AGGTTTTAAACCTCCGGCTTATGGTTTGAAATACCGA | Structure |
| Tetrapod_24[125] | AACTCGCTACGGCGCCCGACAATGACAACAACCGATTATCTG | Structure |
| Tetrapod_26[97] | CTTGCAGGCATTAAACACCTAAATATCTGCATATGATGTC | Structure |
| Tetrapod_30[125] | GATAGTTGCGCAATTGCTAAACAACTCTGAAACGACATCAGCTGGCA | Structure |
| Tetrapod_31[98] | CTGTATGGCACCAGGGTAAAATACGTAATTATTCGGTAGCAAGCGGTTTTG | Structure |
| Tetrapod_34[111] | CAAAATATGCAGATACTCATTCCACAACTAAAGAGGAA | Structure |
| Tetrapod_35[84] | TAGCGTTCTGGAAGTTATAACGCCATACCACGTTAGTA | Structure |
| Tetrapod_35[133] | GTAGATTTAATAAATCGCTAAATTGACCTGGAAGAGTTTC | Structure |
| Tetrapod_38[111] | ATTCATTATAAGCAATAAAGGCAGTTA | Structure |
| Tetrapod_38[132] | CATTATGCATAAAATACAGGCAAGGCAAACCAATTCTGCTCAACAAATAT | Structure |
| Tetrapod_42[118] | TTAACAGAGGTTCGACATTGCCTTGCCGGAGTAAGCG | Structure |
| Tetrapod_43[91] | ACAAAGGCTATCAGGTAGTGAGCGACTTATCACGACAGT | Structure |
| Tetrapod_46[118] | CAGGGTTAGCAATAGGAACGCATCTGCCA | Structure |
| Tetrapod_46[139] | CGAAGAAATAATTATTTTTGTTAAATCAGACAAGAGACCGTTCTAAAGCAA | Structure |
| Tetrapod_47[91] | CTCATGGCTATTTTTGAGCAAAGGTGTTATCTCGGAT | Structure |
| Tetrapod_47[133] | AGATGGTATTCAAATCGATGAACGGTAATCGCATTAACGCGTCTATGGGCG | Structure |
| Tetrapod_0[37] | TGAGCTAACTCACATCGGC TTT | Structure |
| Tetrapod_1[19] | TTT CGGCTACCATTAATTGGCGAAAGGACATCGGC TTT | Structure |
| Tetrapod_3[192] | CGTTATCACCGCGTTTGCCATCTTTTCCATCGGC TTT | Structure |
| Tetrapod_4[38] | GCACACCAGTGGGGCGCCAGGGTGGTTCATCGGC TTT | Structure |
| Tetrapod_4[211] | TTT CGGCTACATAATCAAACCAGTAAGCCATCGGC TTT | Structure |
| Tetrapod_6[192] | TTCCGAATAGCCCGAGATAGCATCGGC TTT | Structure |
| Tetrapod_9[19] | TTT TCGTAATCATCAAGGAACGGTACAAAGCATCAGAACTACTAATGCT TTT | Structure |
| Tetrapod_10[44] | TCTGCCAGAAAAGGGATTTTAGACAGGAACTACTAATGCT TTT | Structure |
| Tetrapod_12[44] | TAGCCAGAACAAACTATCGGCCTTGCGAACTACTAATGCT TTT | Structure |
| Tetrapod_12[211] | TTT ACCCTGAACACGGAATAC TTT | Structure |
| Tetrapod_17[19] | TTT TTGAGTAACTAGCGATAG TTT | Structure |
| Tetrapod_20[184] | ACAAATTCTTACCAGTATAGAACTACTAATGCT TTT | Structure |
| Tetrapod_20[211] | TTT TCGTAATCATCAAGAAGCCAACGAATAATATCGAACTACTAATGCT TTT | Structure |
| Tetrapod_22[184] | TTTTACAAAATCGCGCAGAGAACTACTAATGCT TTT | Structure |
| Tetrapod_25[19] | TTT TCGTAATCATCAAGAGTACCGCCCATCGGAACGAACTACTAATGCT TTT | Structure |
| Tetrapod_26[44] | CAGACCCTCACGTACTCAGGAGGTTTGAACTACTAATGCT TTT | Structure |
| Tetrapod_28[44] | AGCTAGCGTAGCATTCCACAGACAGCGAACTACTAATGCT TTT | Structure |
| Tetrapod_28[211] | TTT GTAGTGTCACTCCATGCA TTT | Structure |
| Tetrapod_33[19] | TTT ACCTGCTCCAAAACCAAA TTT | Structure |
| Tetrapod_36[184] | TTGGGATTAGAGAGTACCTGAACTACTAATGCT TTT | Structure |

| | | |
|---|---|---|
| Tetrapod_36[211] | TTT TCGTAATCATCAAGTTAATTGCTTATTTTCATGAACTACTAATGCT<br>TTT | Structure |
| Tetrapod_38[184] | AGGATTTTAAGAACTGGCTGAACTACTAATGCT TTT | Structure |
| Tetrapod_41[19] | TTT CGGCTACGCCATTCGCGCAGACCTTCATCGGC TTT | Structure |
| Tetrapod_41[39] | CAGGGAAACCAGGCAAAGCCATCGGC TTT | Structure |
| Tetrapod_44[211] | TTT CGGCTACAATGTGTAGGCCCCAAAACATCGGC TTT | Structure |
| Tetrapod_46[34] | TAACGCCAGCATCGGC TTT | Structure |
| Tetrapod_47[193] | AATAAAAATGCGCATCGGC TTT | Structure |
| Tetrapod_8[213] | TTT AGACAAAAGGGCGACAGGTTTACAAAGCGTACAGAGATAGAACCC<br>T TTT | Structure |
| Tetrapod_10[213] | TTT CCAAAAGAACTGGCATAATAATAAAAGTCAGGAGAATTAACTGAA<br>C TTT | Structure |
| Tetrapod_24[214] | TTT ACTTGCCCTCTCTGTAGTACGGTCTCCAAAACGTTGAAAATCTCC<br>A TTT | Structure |
| Tetrapod_37[16] | TTT GAATGACCATAAATCAACAGTTCACCGGATTTCATCAAGAGTAAT<br>C TTT | Structure |
| Tetrapod_0[214] | TTT CGGCTAC GCCGTCGAGAGGGTTGTACCAGGTTG | Structure |
| Tetrapod_2[214] | TTT CGGCTAC GTCATACATGGCTTTTTAC | Structure |
| Tetrapod_3[16] | TTT CGGCTAC GCGGGCGCTAGGGCGCGAAGAAACGTTGCGTGAG | Structure |
| Tetrapod_5[16] | TTT CGGCTAC GCTTTCCTCGTTAGAACGA | Structure |
| Tetrapod_6[214] | TTT CGGCTAC GGTTGAGTGCGGATAAGT CATCGGC TTT | Structure |
| Tetrapod_8[213] | TTT AGACAAAAGGGCGACAGGTTTACAAAGCGTACAGAGATAGAACCC<br>T | Structure |
| Tetrapod_10[213] | TTT CCAAAAGAACTGGCATAATAATAAAAGTCAGGAGAATTAACTGAA<br>C | Structure |
| Tetrapod_11[16] | TTT TCGTAATCATCAAG CCTTGCTGAAATCCTTGAGAAGACAAATC<br>CTCAA | Structure |
| Tetrapod_13[16] | TTT TCGTAATCATCAAG ACTTTACAACGAACGTTTTTGCGTATACT<br>TCAAA | Structure |
| Tetrapod_14[214] | TTT TCTGACCTGCAGCGCCAA TTT | Structure |
| Tetrapod_15[16] | TTT TCGTAATCATCAAG TGGTAATATAAGTATTAG GAACTACTA<br>ATGCT TTT | Structure |
| Tetrapod_19[16] | TTT CTTAGATTAAGACGCTGAAAACAATTATCATATTAATTTTAAAAG<br>T | Structure |
| Tetrapod_21[16] | TTT ACGCGAGAAAACTTTTAATCGCAACCATATCTGAATAATGGAAGG<br>G | Structure |
| Tetrapod_22[214] | TTT TCGTAATCATCAAG GGCGAATTATCCGGTATT GAACTACTA<br>ATGCT TTT | Structure |
| Tetrapod_23[16] | TTT TTAGAACCTAGACAAAGA TTT | Structure |
| Tetrapod_24[214] | TTT ACTTGCCCTCTCTGTAGTACGGTCTCCAAAACGTTGAAAATCTCC<br>A | Structure |
| Tetrapod_27[17] | TTT TCGTAATCATCAAGGAGGGTAGCACCAGACGCAAAAGGCT TTT | Structure |
| Tetrapod_30[214] | TTT AAAAAAAGGGGAAACGAT TTT | Structure |
| Tetrapod_39[16] | TTT TTGACAAGAAGAAAACGA TTT | Structure |
| Tetrapod_46[214] | TTT CGGCTACCCGCCTGAAACCCGTCGGCATCGGC TTT | Structure |
| Tetrapod_16[214] | TTT TCGTAATCATCAAGCTAAGAACGTTTTGGTGCCGCTGCGCGCGATT<br>AACG TTT | Structure |
| Tetrapod_18[214] | TTT TCGTAATCATCAAGCCATCCTAATTTACTGGGGCACCATCATATTG<br>CAAAC TTT | Structure |
| Tetrapod_34[214] | TTT TCGTAATCATCAAGTTGGGGCGCTTTTGATAATAGCAAATGTGAGC<br>GAACGGCG TTT | Structure |
| Tetrapod_40[214] | TTT CGGCTACATTCTCCGTTTTTTAGACGGAGGGTACTGGCCAAAGAAT<br>A TTT | Structure |
| Tetrapod_42[214] | TTT CGGCTACACAGGAAGATTTGAAACGCGATTAAGTTCA TTT | Structure |
| Tetrapod_43[16] | TTT CGGCTACTTCCATGAATTTATCCGGTTTATGTAATCA TTT | Structure |
| Tetrapod_7[16]_docking_a | TTTCGGCTACTTTCTTTTCGTATAACGTCATCGGCTTTCCTCCTCCTCCT<br>CCT | docking strand a |
| Tetrapod_31[16]_docking_b | TTTTCGTAATCATCAAGCCTCATAGTGATTATACCGAACTACTAATGCTT<br>TTCCTCCTCCTCCTCCT | docking strand b |
| Tetrapod_38[214]_docking_c | TTTTCGTAATCATCAAGCATTATACCCTTTATTTCGAACTACTAATGCTT<br>TTCCTCCTCCTCCTCCT | docking strand c |
| Tetrapod_28_47[16]_docking_d | TTTCGGCTACGGTTTTCCCGTAGGGAAACATCGGCTTTCCTCCTCCTCCT<br>CCT | docking strand d |
| Tetrapod_29[17]_biotin_b1 | /5Biosg/TTTTCGTAATCATCAAGAAGCGCGAAGATAAATTTAGCCGGC<br>TGACCATTCATT | biotinylated strand b1 |
| Tetrapod_35[16]_biotin_b2 | TTTATAGCGAGAGGCTTTTGACGATAATGTTACTGTGTCGAAATCCGCGT<br>TT/3Bio/ | biotinylated strand b2 |
| Tetrapod_32[215]_biotin_c1 | /5Biosg/TTTTCGTAATCATCAAGAACGCAAGGTTTTATTTTATTCAAA<br>ACGCAACAGCT | biotinylated strand c1 |
| Tetrapod_44[191]_biotin_c2 | AGAAATGCAATGCCTGAGTCATCGGCTTT/3Bio/ | biotinylated strand c2 |
| Tetrapod_8_45[16]_biotin_d1 | /5Biosg/TTTCGGCTACCTGCGTGTTTTTTCTTCACACCGTACTTTTTT<br>CAGGAGCC | biotinylated strand d1 |

| | | |
|---|---|---|
| Tetrapod_26[214]_biotin_d2 | TTTGACATCACGAAGGTGTGTCTTGTGATGATGAGCGATGCCAGAGTCTTTT/3Bio/ | biotinylated strand d2 |
| Tetrapod_7[16]_nondocking_a | TTTCGGCTACTTTCTTTTCGTATAACGTCATCGGCTTT | structure at position docking strand a (nondocking a) |
| Tetrapod_31[16]_nondocking_b | TTTTCGTAATCATCAAGCCTCATAGTGATTATACCGAACTACTAATGCTTTT | structure at position docking strand b (nondocking b) |
| Tetrapod_38[214]_nondocking_c | TTTTCGTAATCATCAAGCATTATACCCTTTATTTCGAACTACTAATGCTTTT | structure at position docking strand c (nondocking c) |
| Tetrapod_47[16]_nondocking_d | TTTCGGCTACGGTTTTCCCGTAGGGAAACATCGGCTTT | structure at position docking strand d (nondocking d) |
| Tetrapod_29[17]_nonbiotin_b1 | TTTTCGTAATCATCAAGAAGCGCGAAGATAAATTTAGCCGGCTGACCATTCATT | structure at position biotinylated strand b1 (nonbiotinylated b1) |
| Tetrapod_35[16]_nonbiotin_b2 | TTTATAGCGAGAGGCTTTTGACGATAATGTTACTGTGTCGAAATCCGCGTTT | structure at position biotinylated strand b2 (nonbiotinylated b2) |
| Tetrapod_32[215]_nonbiotin_c1 | TTTTCGTAATCATCAAGAACGCAAGGTTTTATTTTATTCAAAACGCAACAGCT | structure at position biotinylated strand c1 (nonbiotinylated c1) |
| Tetrapod_44[191]_nonbiotin_c2 | AGAAATGCAATGCCTGAGTCATCGGCTTT | structure at position biotinylated strand c2 (nonbiotinylated c2) |
| Tetrapod_45[16]_nonbiotin_d1 | TTTCGGCTACCTGCGTGTTTTTTCTTCACACCGTACTTTTTTCAGGAGCC | structure at position biotinylated strand d1 (nonbiotinylated d1) |
| Tetrapod_26[214]_nonbiotin_d2 | TTTGACATCACGAAGGTGTGTCTTGTGATGATGAGCGATGCCAGAGTCTTTT | structure at position biotinylated strand d2 (nonbiotinylated d2) |
| Tetrapod_14[125]_docking_vertex | AAAACATCGCCAAGCTCAATCGTCTGGAAACAATGAAACTAAACGATTTTCCTCCTCCTCCTCCT | vertex docking strand |
| Tetrapod_7[46]_docking_mid_arm | GGGCAACAGCTATGGTTTGTAGCGCAAATGAATATCAAATTTGAGGACAATTTCCTCCTCCTCCTCCT | mid arm docking strand |
| short | GCCGATGGTAGCCG | short staple scaffold |
| long | AGCATTAGTAGTTCCTTGATGATTACGA | long staple scaffold |
| Tetrapod_7[16]_dimer_connector_monomer_1 | GTACGGTCAACGGCTACTTTCTTTTCGTATAACGTCATCGGCGTACGGTCAA | dimerization connector strands for monomer 1 |
| Tetrapod_5[16]_dimer_connector_monomer_1 | GTACGGTCAACGGCTACGCTTTCCTCGTTAGAACGA | dimerization connector strands for monomer 1 |
| Tetrapod_3[16]_dimer_connector_monomer_1 | GTACGGTCAACGGCTACGCGGGCGCTAGGGCGCGAAGAAACGTTGCGTGAG | dimerization connector strands for monomer 1 |
| Tetrapod_15[16]_dimer_connector_monomer_1 | GTACGGTCAATCGTAATCATCAAGTGGTAATATAAGTATTAGGAACTACTAATGCT | dimerization connector strands for monomer 1 |
| Tetrapod_21[16]_dimer_connector_monomer_1 | ACGTTAGCATACGCGAGAAAACTTTTAATCGCAACCATATCTGAATAATGGAAGGG | dimerization connector strands for monomer 1 |
| Tetrapod_23[16]_dimer_connector_monomer_1 | ACGTTAGCATTTAGAACCTAGACAAAGAACGTTAGCAT | dimerization connector strands for monomer 1 |
| Tetrapod_19[16]_dimer_connector_monomer_1 | ACGTTAGCATCTTAGATTAAGACGCTGAAAACAATTATCATATTAATTTTAAAAGT | dimerization connector strands for monomer 1 |
| Tetrapod_13[16]_dimer_connector_monomer_1 | ACGTTAGCATTCGTAATCATCAAGACTTTACAACGAACGTTTTTGCGTATACTTCAAA | dimerization connector strands for monomer 1 |
| Tetrapod_7[16]_dimer_connector_monomer_2 | ATGCTAACGTCGGCTACTTTCTTTTCGTATAACGTCATCGGCATGCTAACGT | dimerization connector strands for monomer 2 |
| Tetrapod_5[16]_dimer_connector_monomer_2 | ATGCTAACGTCGGCTACGCTTTCCTCGTTAGAACGA | dimerization connector strands for monomer 2 |
| Tetrapod_3[16]_dimer_connector_monomer_2 | ATGCTAACGTCGGCTACGCGGGCGCTAGGGCGCGAAGAAACGTTGCGTGAG | dimerization connector strands for monomer 2 |
| Tetrapod_15[16]_dimer_connector_monomer_2 | ATGCTAACGTTCGTAATCATCAAGTGGTAATATAAGTATTAGGAACTACTAATGCT | dimerization connector strands for monomer 2 |
| Tetrapod_14_21[16]_dimer_connector_monomer_2 | TTGACCGTACACGCGAGAAAACTTTTAATCGCAACCATATCTGAATAATGGAAGGG | dimerization connector strands for monomer 2 |
| Tetrapod_23[16]_dimer_connector_monomer_2 | TTGACCGTACTTAGAACCTAGACAAAGATTGACCGTAC | dimerization connector strands for monomer 2 |
| Tetrapod_19[16]_dimer_connector_monomer_2 | TTGACCGTACCTTAGATTAAGACGCTGAAAACAATTATCATATTAATTTTAAAAGT | dimerization connector strands for monomer 2 |
| Tetrapod_13[16]_dimer_connector_monomer_2 | TTGACCGTACTCGTAATCATCAAGACTTTACAACGAACGTTTTTGCGTATACTTCAAA | dimerization connector strands for monomer 2 |
| Tetrapod_7[16]_no_dimerization | TTT CGGCTACTTTCTTTTCGTATAACGTCATCGGC TTT | no dimerization strands |
| Tetrapod_5[16]_no_dimerization | TTT CGGCTAC GCTTTCCTCGTTAGAACGA | no dimerization strands |
| Tetrapod_3[16]_no_dimerization | TTT CGGCTAC GCGGGCGCTAGGGCGCGAAGAAACGTTGCGTGAG | no dimerization strands |
| Tetrapod_15[16]_no_dimerization | TTT TCGTAATCATCAAG TGGTAATATAAGTATTAG GAACTACTAATGCT TTT | no dimerization strands |
| Tetrapod_14_21[16]_no_dimerization | TTT ACGCGAGAAAACTTTTAATCGCAACCATATCTGAATAATGGAAGGG | no dimerization strands |
| Tetrapod_23[16]_no_dimerization | TTT TTAGAACCTAGACAAAGA TTT | no dimerization strands |
| Tetrapod_19[16]_no_dimerization | TTT CTTAGATTAAGACGCTGAAAACAATTATCATATTAATTTTAAAAGT | no dimerization strands |
| Tetrapod_13[16]_no_dimerization | TTT TCGTAATCATCAAG ACTTTACAACGAACGTTTTTGCGTATACTTCAAA | no dimerization strands |

**Table S7** **DNA origami tetrapod strands.** Scaffold is p8634. Structure staples are all the strands that form the structure. Biotinylated staples are modified with biotin to immobilize origami on coverslip surfaces through a BSA-Biotin-Streptavidin-Biotin-DNA origami arrangement. Docking staples are for DNA-PAINT experiment. Dimerization connectors are for dimer assembly.

| Strands | monomer with 4 dockings | 1 at 0 | | 7 at 1 | | 7 at 2 | | 6 at 3 | |
|---|---|---|---|---|---|---|---|---|---|
| | | Monomer 1 | Monomer 2 | Monomer 1 | Monomer 2 | Monomer 1 | Monomer 2 | Monomer 1 | Monomer 2 |
| P8634 | Y | Y | Y | Y | Y | Y | Y | Y | Y |
| Structure | Y | Y | Y | Y | Y | Y | Y | Y | Y |
| vertex position structure | Y | Y | Y | N | Y | N | N | N | N |
| mid arm position structure | Y | N | N | N | N | N | N | N | Y |
| docking strand a | Y | N | N | N | N | N | N | N | N |
| docking strand b | Y | N | Y | N | Y | N | Y | N | Y |
| docking strand c | Y | N | N | N | Y | N | N | N | Y |
| docking strand d | Y | N | Y | Y | Y | Y | Y | Y | Y |
| vertex docking strand | N | N | N | Y | N | Y | Y | Y | N |
| mid arm docking strand | N | Y | Y | Y | Y | Y | Y | Y | Y |
| biotinylated strand b1 | Y | N | N | N | N | N | N | N | N |
| biotinylated strand b2 | Y | N | N | N | N | N | N | N | N |
| biotinylated strand c1 | Y | Y | Y | Y | Y | Y | Y | Y | Y |
| biotinylated strand c2 | Y | Y | Y | Y | Y | Y | Y | Y | Y |
| biotinylated strand d1 | Y | Y | Y | Y | Y | Y | Y | Y | Y |
| biotinylated strand d2 | Y | Y | Y | Y | Y | Y | Y | Y | Y |
| structure at position docking strand a (nondocking a) | N | Y | Y | Y | Y | Y | Y | Y | Y |
| structure at position docking strand b (nondocking b) | N | Y | N | Y | N | Y | N | Y | N |
| structure at position docking strand c (nondocking c) | N | Y | Y | Y | N | N | Y | Y | N |
| structure at position docking strand d (nondocking d) | N | Y | N | N | N | N | N | N | N |
| structure at position biotinylated strand b1 (nonbiotinylated b1) | N | Y | Y | Y | Y | Y | Y | Y | Y |
| structure at position biotinylated strand b2 (nonbiotinylated b2) | N | Y | Y | Y | Y | Y | Y | Y | Y |
| structure at position biotinylated strand c1 (nonbiotinylated c1) | N | N | N | N | N | N | N | N | N |
| structure at position biotinylated strand c2 (nonbiotinylated c2) | N | N | N | N | N | N | N | N | N |
| structure at position biotinylated strand d1 (nonbiotinylated d1) | N | N | N | N | N | N | N | N | N |
| structure at position biotinylated strand d2 (nonbiotinylated d2) | N | N | N | N | N | N | N | N | N |
| short staple scaffold | Y | Y | Y | Y | Y | Y | Y | Y | Y |
| long staple scaffold | Y | Y | Y | Y | Y | Y | Y | Y | Y |
| dimerization connector strands for monomer 1 | N | Y | N | Y | N | Y | N | Y | N |
| dimerization connector strands for monomer 2 | N | N | Y | N | Y | N | Y | N | Y |
| no dimerization strands | Y | N | N | N | N | N | N | N | N |

**Table S8** **Tetrapod Pattern Mixing.** The assembly contained 30 nM scaffold P8634; 300 nM of each structure strand; 6 $\mu$M of each biotinylated strand; 4.5 $\mu$M of each short and long staple scaffold; 300 nM of both non-dimerizing strands and dimerization connector strands specific to the monomer; 5 $\mu$M of each docking strand; and 300 nM of each non-docking strand.

| Parameters | Values |
|---|---|
| Box side length | 7 for 2D, and 9 for 3D |
| Min. Net Gradient | 15,000 for 2D, and 10,000 for 3D<br>(filtering can be done later using Picasso Filter module) |
| EM Gain | 1 |
| Baseline | 100 |
| Sensitivity | 0.46 |
| Quantum efficiency (at Cy3B emission) | 0.82 |
| Pixel size | 117 nm |
| Method | MLE, integrated Gaussian for 2D, and LQ, Gaussian for 3D |
| 3D via Astigmatism | Empty for 2D, and Use a calibration file for 3D |

**Table S9 Picasso localize module parameters.** The parameters are based on the Hamamatsu ORCA-Flash4.0 V3 digital sCMOS camera.

| Parameter | NSF 2D dataset | ASU one-redundancy 2D dataset | ASU two-redundancy 2D dataset | 0407 3D dataset |
|---|---|---|---|---|
| $N$ | 3 | 9 | 9 | 0 |
| $M$ | 13 | 49 | 49 | 13 |
| $T_{\mathrm{I}}$ | 0.15 | 0.2 | 0.2 | 95% |
| $T_{\mathrm{S}}$ | 0.5 | 0.7 | 0.8 | Not applicable |
| $W_{\mathrm{O}}$ | 1.5 | 1.5 | 1.5 | 1 |
| Alignment | Rough | Differential evolution | Differential evolution | Steps of translation followed by rotation concerning z-axis |

**Table S10 Parameter selection.** It is done by empirically finding good values for $N$ and $M$. If origami with higher numbers of binding sites are imaged, we may need a higher value of $M$ to account for false positives. $T_{\mathrm{I}}$ and $T_{\mathrm{S}}$ were selected through grid search. $W_{\mathrm{O}}$ was empirically selected. The best method for alignment for each dataset was empirically selected.

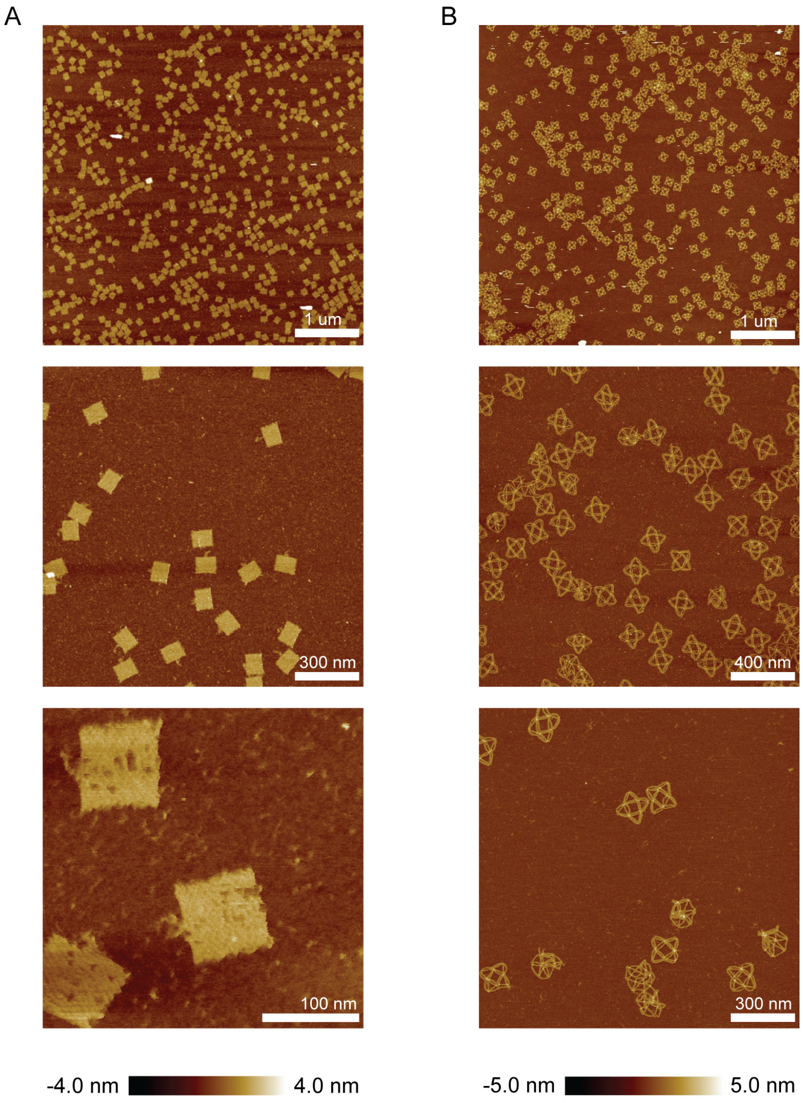

**Fig. S1 Additional AFM images of 2D RRO and 3D wireframe cuboctahedron DNA origami.**(**A**) AFM images of 2D RRO with varying fields of view. (**B**) AFM images of 3D cuboctahedron DNA origami with varying fields of view.

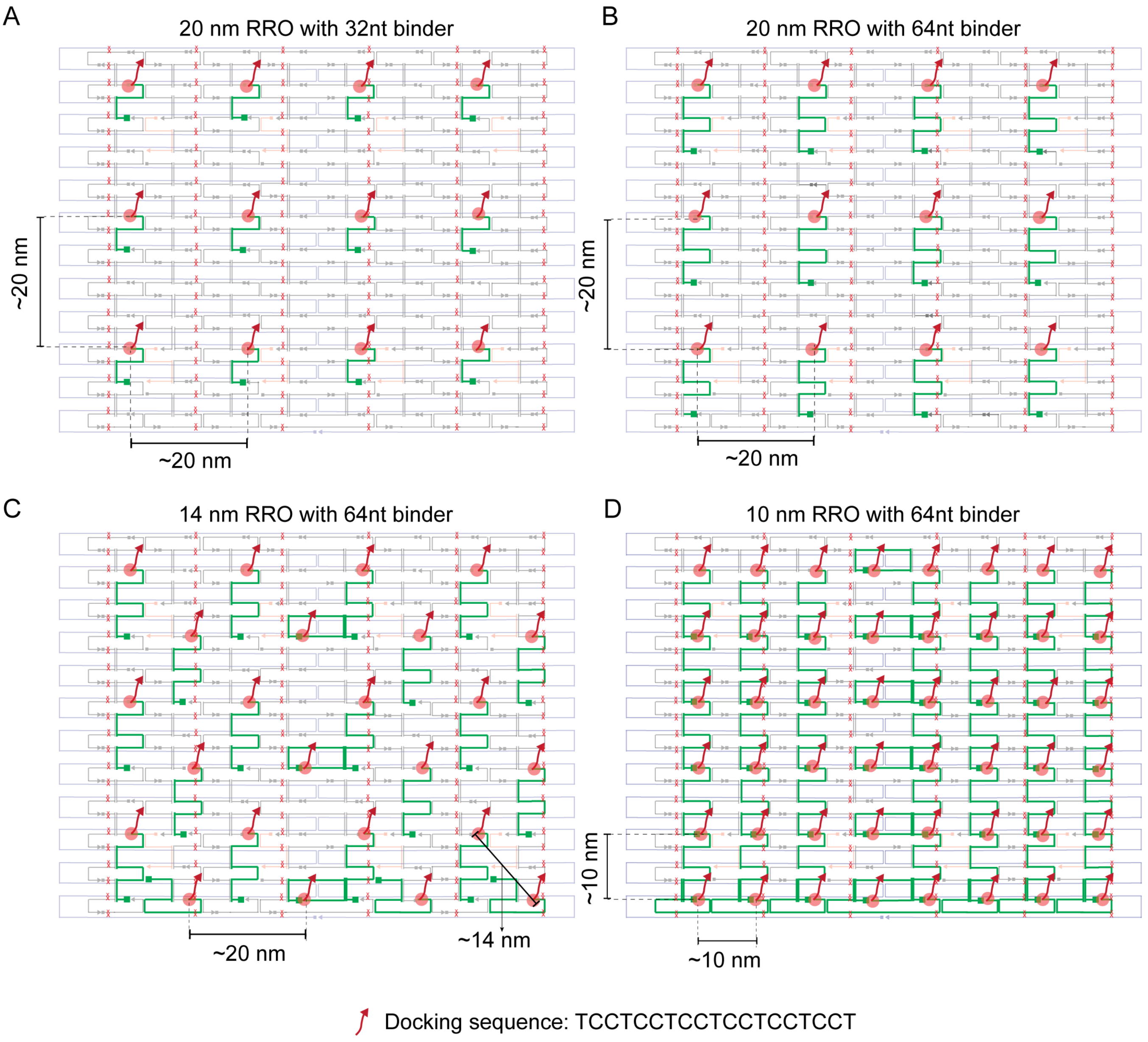

**Fig. S2 2D RRO cadnano design showing scaffold routing and staple strands interlacing** (**A**) Map of 20 nm 2D RRO with 32 nt binder and 20 nm separation between imager binding locations. (**B**) Map of 20 nm 2D RRO with 64nt binder and 20 nm separation between imager binding locations. (**C**) Map of 14 nm 2D RRO with long 64 nt binder and 14 nm and 20 nm separation between imager binding locations. (**D**) Map of 10 nm 2D RRO with 64 nt binder and 10 nm separation between imager binding locations.

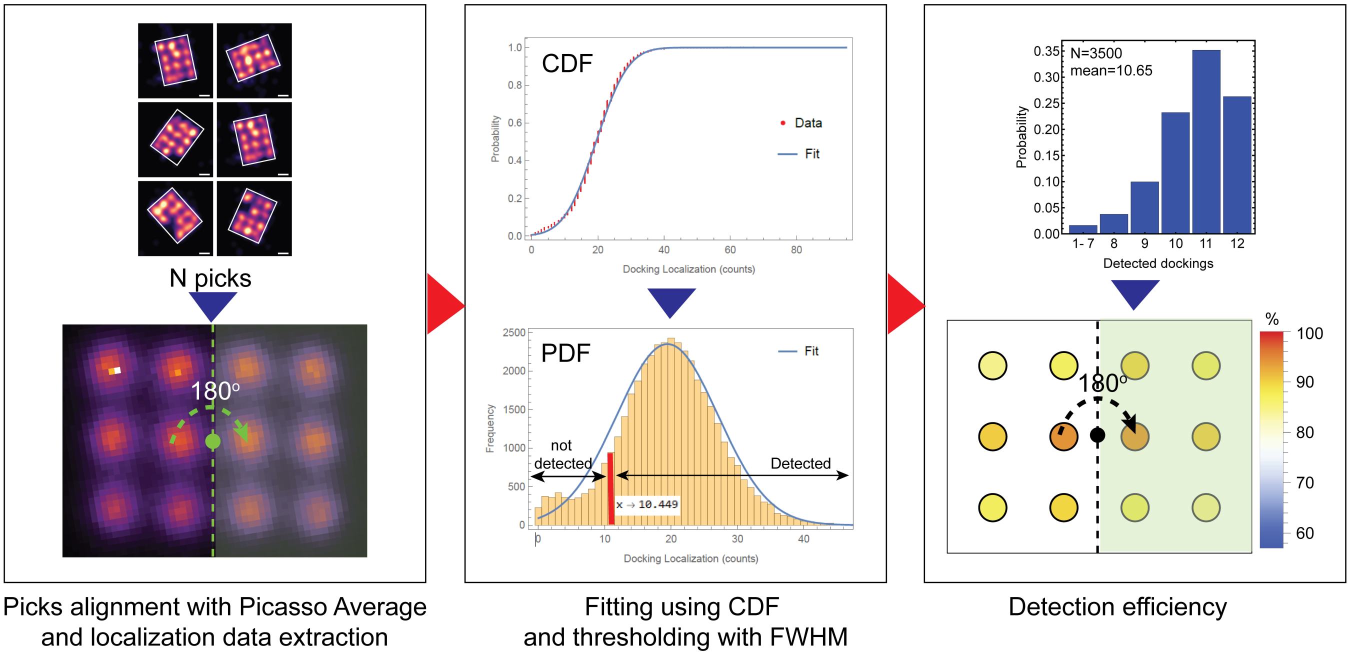

**Fig. S3** **Detection efficiency analysis procedure in 20 nm RRO with 32 nt binder.** Procedure for docking detection efficiency of 20 nm RRO with 32 nt binder. The leftmost panel shows the Picasso Render picking process (top) being input to the Picasso average module to align the picks (bottom). The middle panel is the incorporation distribution of localization for each docking in all picks (bottom) fitted by using cumulative distribution function (CDF) (top) and thresholded by using full-width half maximum (FWHM). The rightmost panel shows localization distribution after thresholding and each docking detection efficiency from all picks.

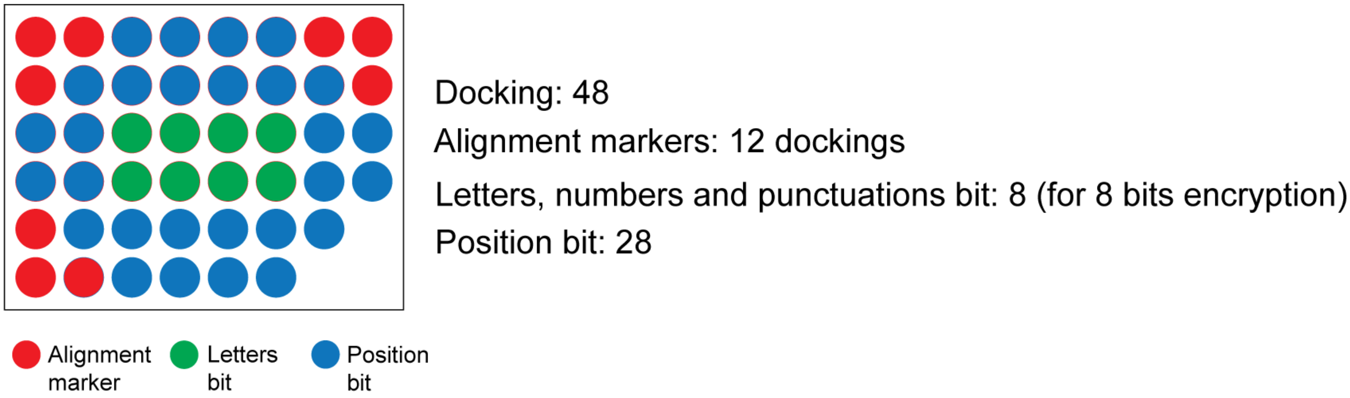

**Fig. S4 Theoretical design of 10 nm resolution encryption pattern resulting in $2^{28}$ combinations of numbers, letters, and punctuation marks forming texts assuming 100% incorporation efficiency.** Design of 10 nm encryption showing the alignment marker with 12 docking, which breaks the symmetry of the design by including only 9 dockings in design (red), letters bit with 8 dockings (green), and position bit with 28 dockings (blue).

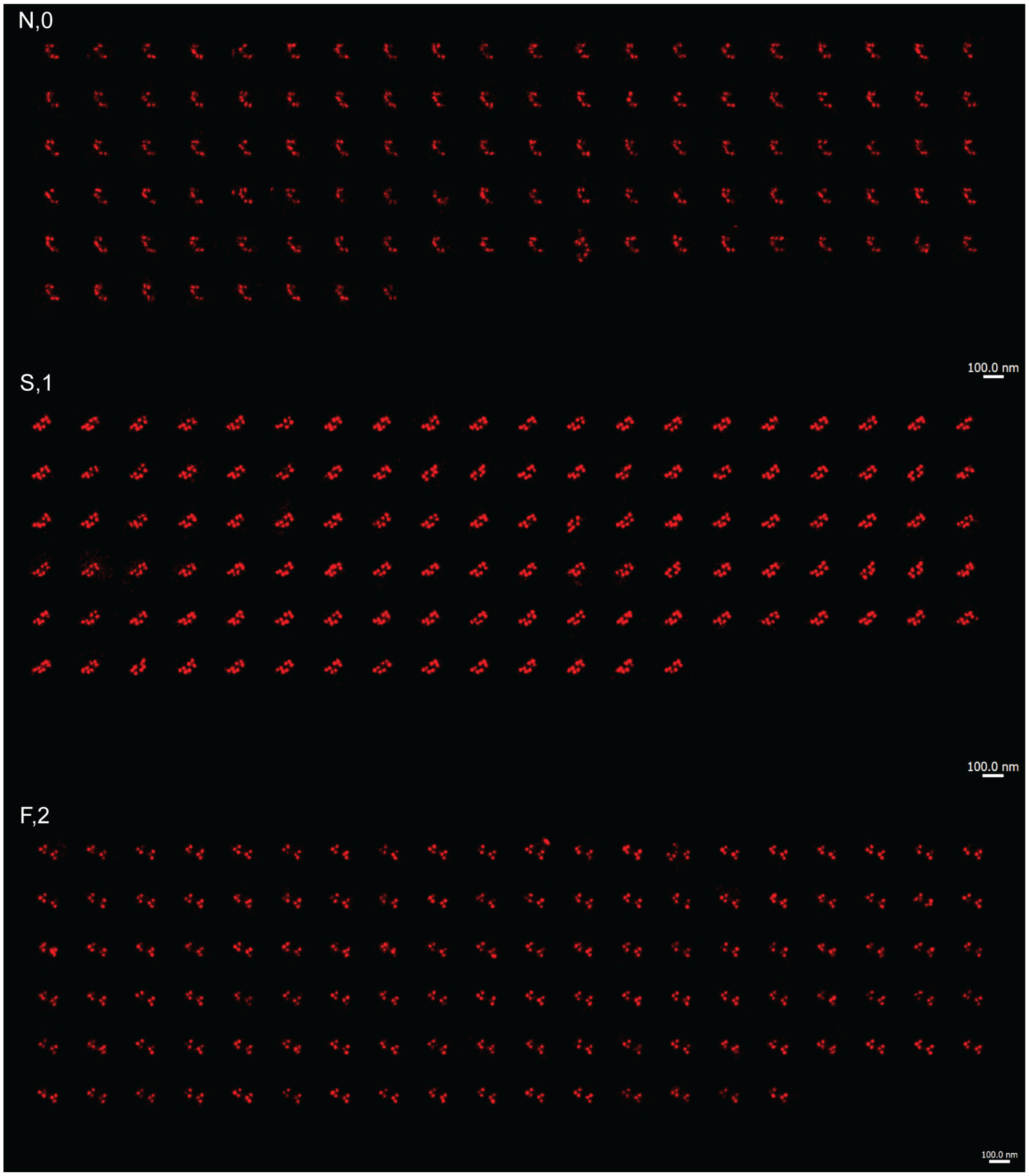

**Fig. S5 All picks in the analyzed "NSF" dataset.** Full data set of 20 nm encrypted "NSF" following Picasso Average alignment and Picasso render unfolding with a 100 nm scale bar.

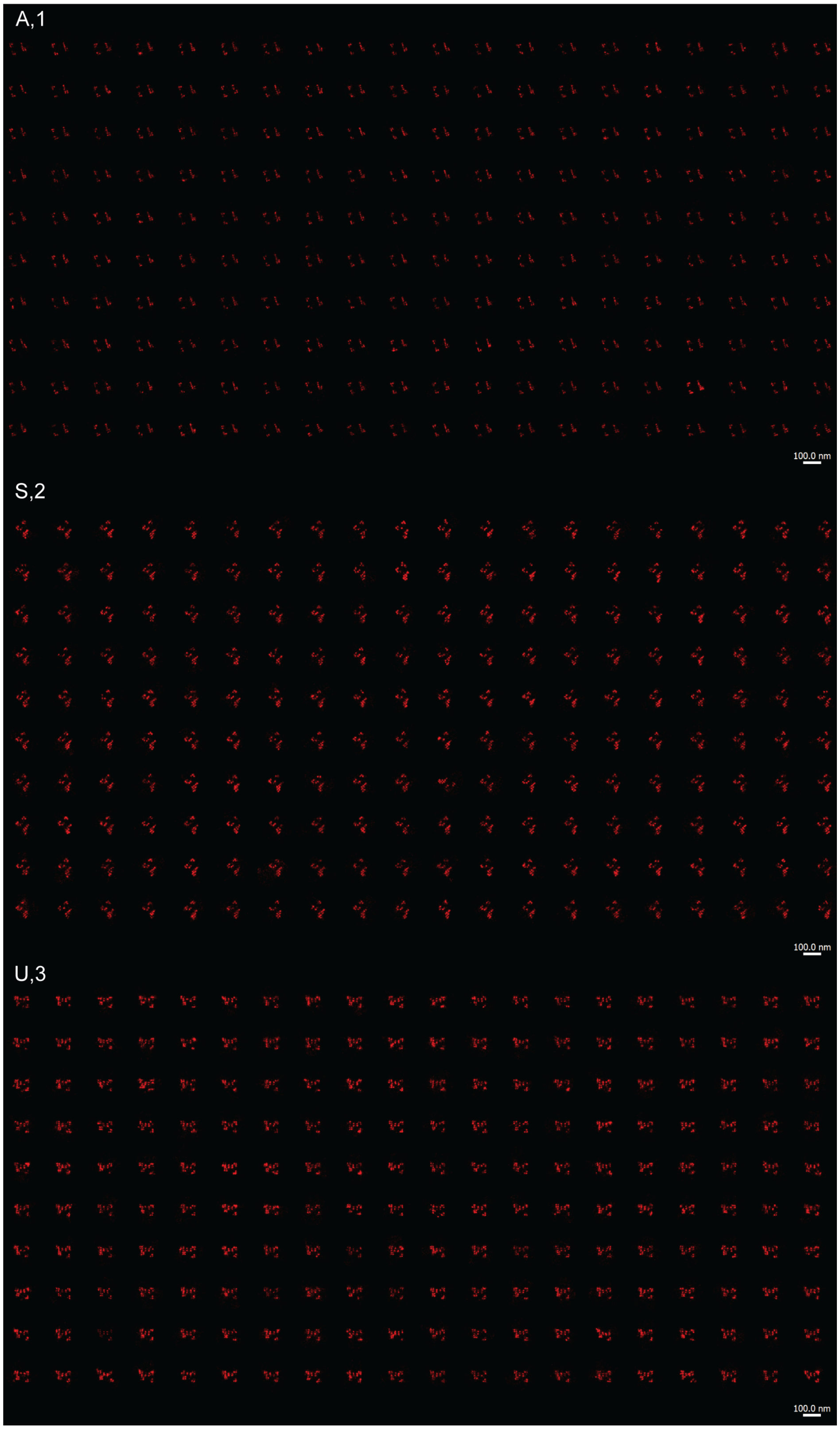

**Fig. S6 All picks in the analyzed "ASU" one redundancy dataset.** Full data set of 10 nm 1 redundancy encrypted "ASU" following Picasso Average alignment and Picasso render unfolding with a 100 nm scale bar.

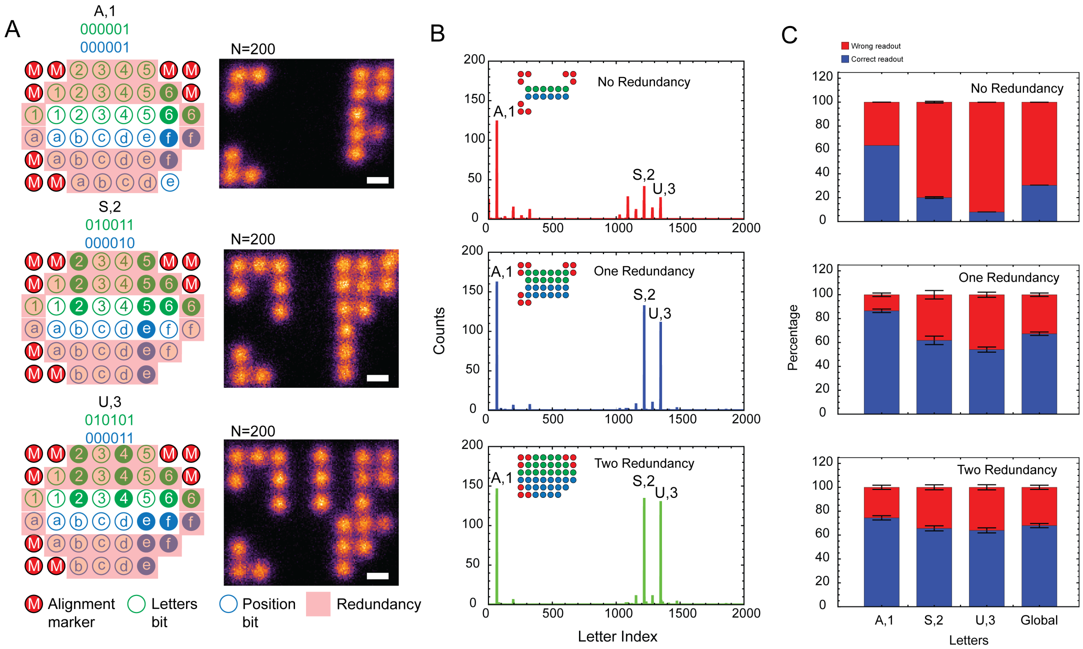

**Fig. S7 "ASU" two redundancy dataset on higher density 2D RRO along with All analyzed picks.** (**A**) The pattern encryption rules for "ASU" two redundancy dataset that shows the alignment marker, letters bit, position bit, and the redundancy (left) and the summed DNA-PAINT images of three letters of "ASU" with the scale bar of 10 nm (right). (**B**) The readout of the "ASU" dataset presented as letter index vs. counts analyzed by not including the redundancy (top) and the redundancy (middle and bottom). (**C**) The readout percentage of correct and wrong readout for each letter and global. The error bar is the standard deviation from 3 different processing runs with the same dataset.

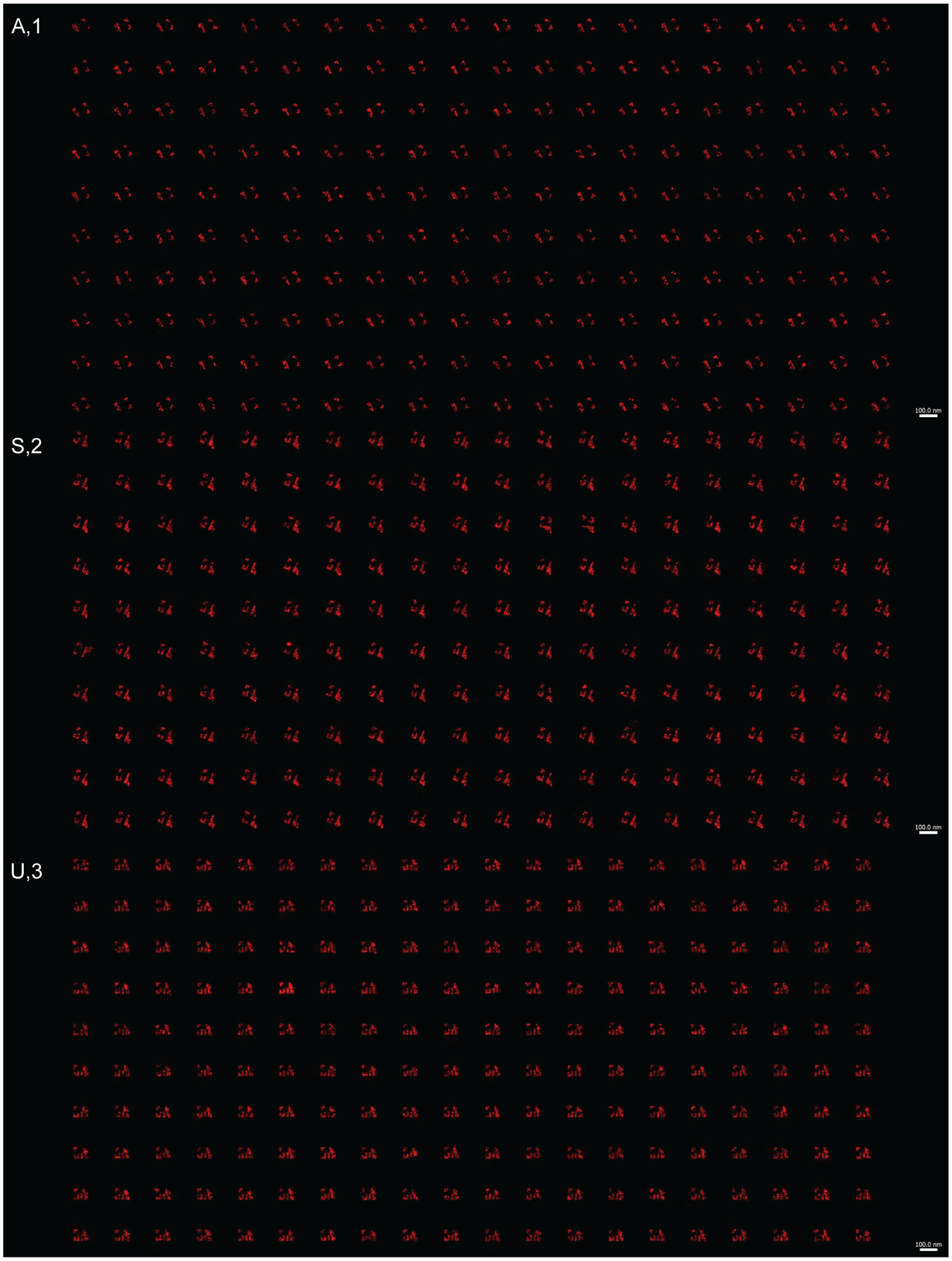

**Fig. S8 All analyzed picks in "ASU" two redundancy dataset on higher density 2D RRO.** Full data set of 10 nm 2 redundancy encrypted "ASU" following Picasso Average alignment and Picasso render unfolding with a 100 nm scale bar.

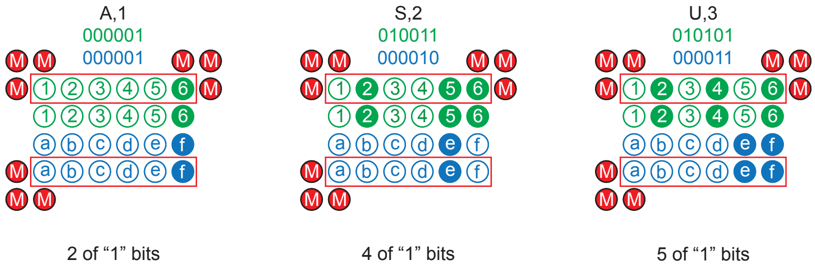

**Fig. S9 Argument for the readout accuracy with increasing bit usage by a pattern.** The encryption pattern schematic for “ASU” 1 redundancy with different bit “1” usage (top). According to Figure 2 of the main text, the estimated detection efficiency of docking is 85–90%. Assuming a detection efficiency of 85%, the probability of at least one pair of dockings (with one redundancy) being detected is $1-0.15^2$, resulting in a probability of 0.978. For A,1 pattern with 2 “1” bits, the probability of correctly reading the pattern is $0.978^2$=0.956. The same estimation applies for S,2 with 4 “1” bits and U,3 with 5 “1” bits, resulting in probabilities of $0.978^4$=0.915 and $0.978^5$=0.895, respectively. However, alignment accuracy and k-means assignment accuracy must also be considered. Assuming an alignment and k-means accuracy of around 90%, both processes contribute to an accuracy of $\sim$81%. Using this value, the overall probability is (0.956)·(0.81)=0.77 for A,1 pattern, (0.915)·(0.81)=0.74 for S,2 pattern, and (0.895)·(0.81)=0.72 for U,3 pattern. The correct readout percentage decreases as the number of “1” bits increases. Although the values do not exactly match the experimental results, this may be due to an additional factor in alignment accuracy that tends to decrease with more “1” bits.

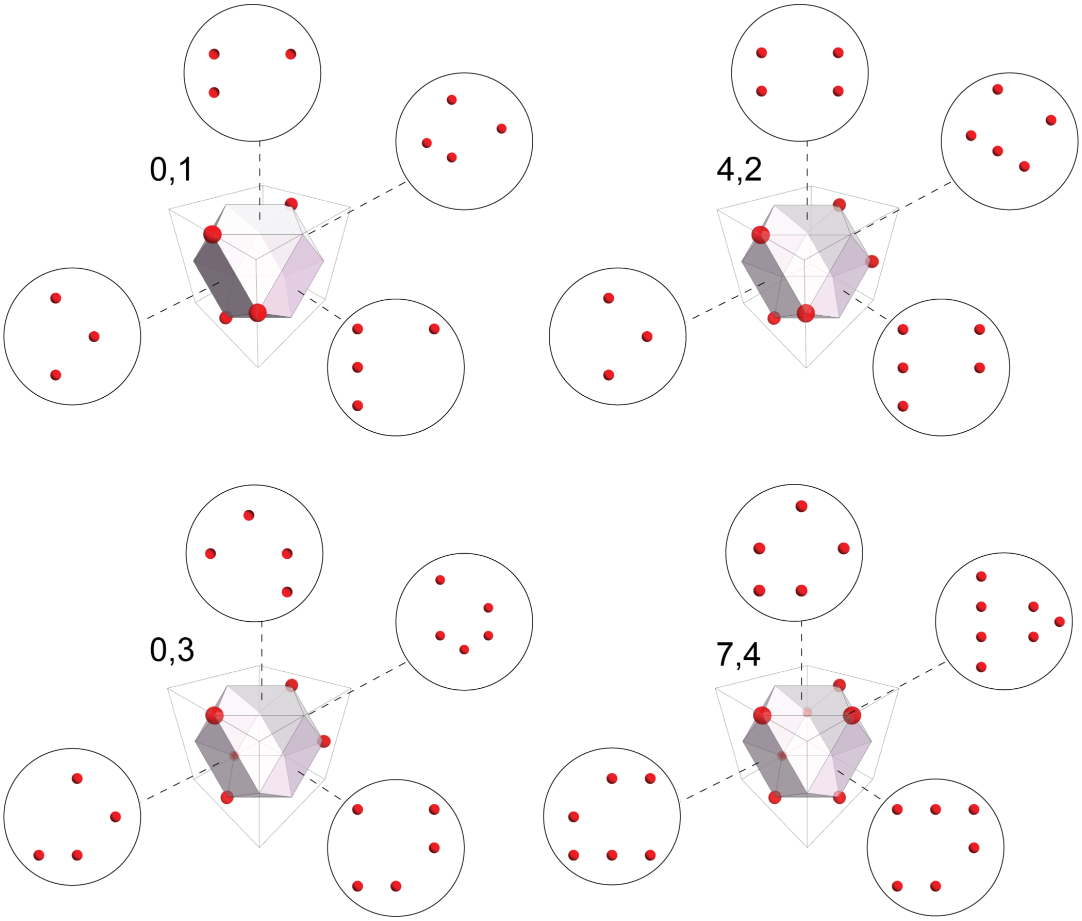

**Fig. S10 Schematics of confused patterns due to 2D projections from 3D DNA origami encryption design.** The biotinylated strands dictate the 2D projections of each pattern when imaged using 2D DNA-PAINT only.

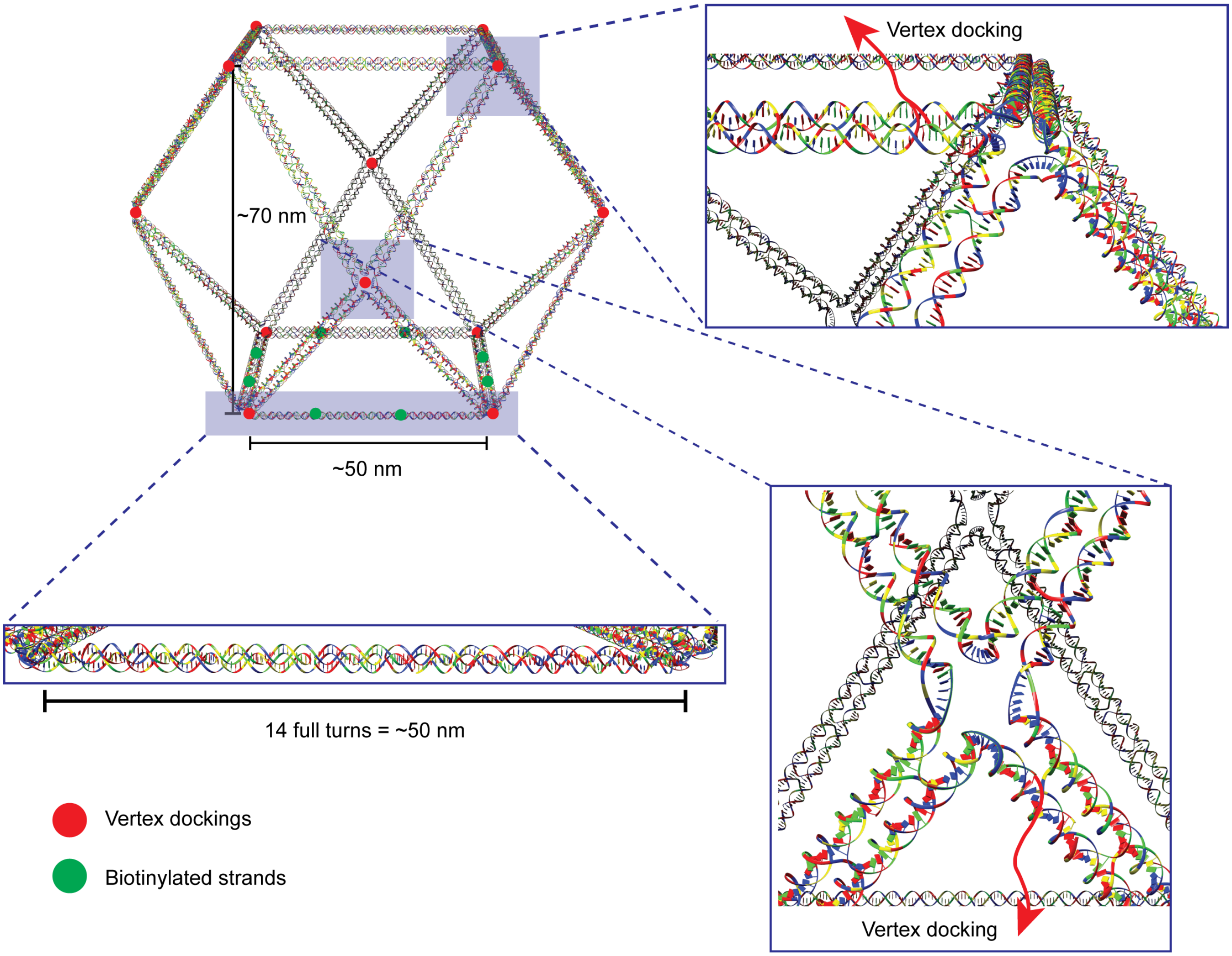

**Fig. S11 3D wireframe cuboctahedron DNA origami design.** The 3D wireframe cuboctahedron has a height of 70 nm and square faces with 50 nm side length due to 14 full-duplex turns (left). Vertex dockings and biotinylated strands are shown as red and green circles, respectively. The right panels show zoomed-in images of 2 vertices.

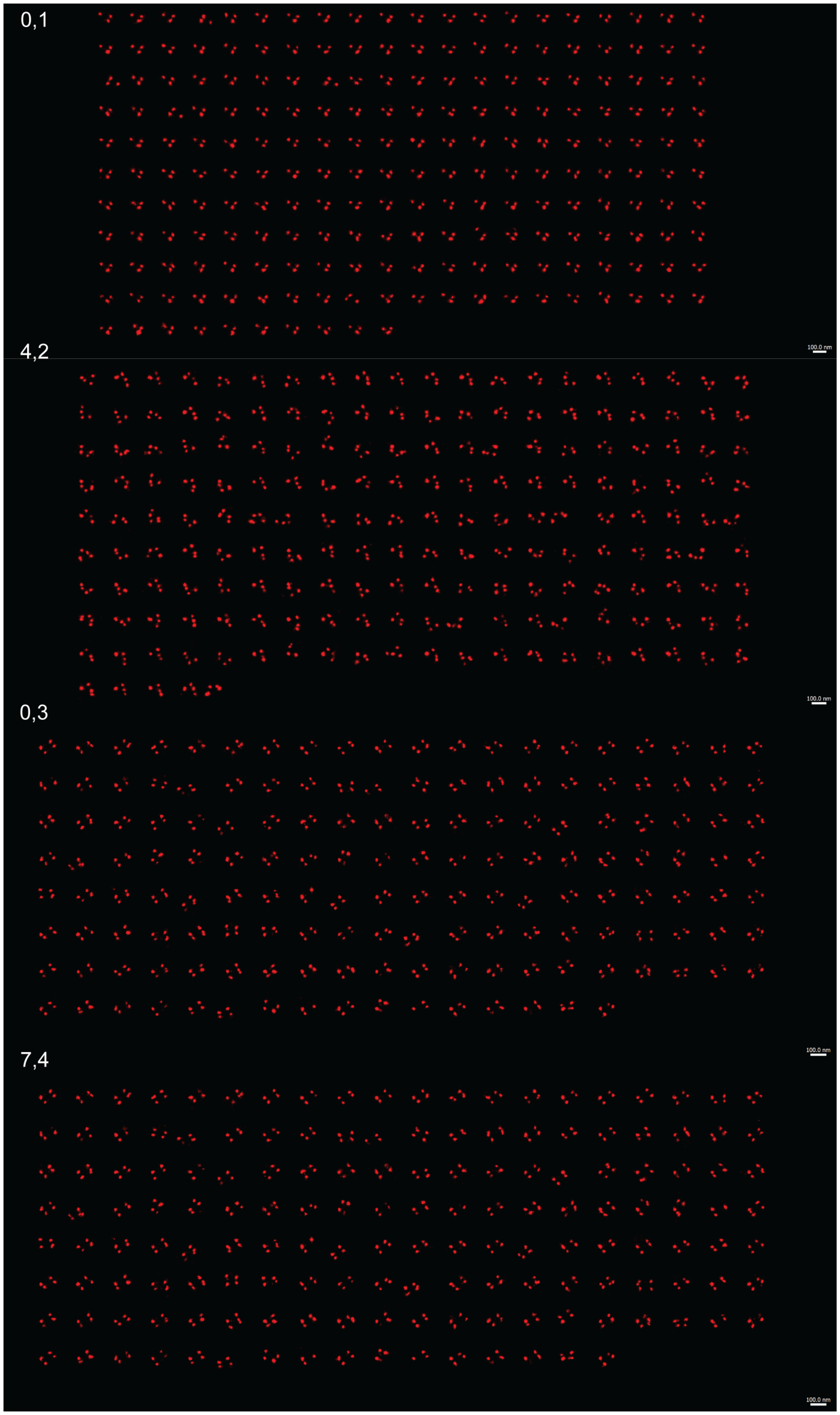

**Fig. S12 2D view of all picks analyzed in 3D DNA-PAINT “0407” dataset.** Full dataset of 2D projection of “0407” dataset following Picasso Average alignment and Picasso render unfolding. Scale bar: 100 nm.

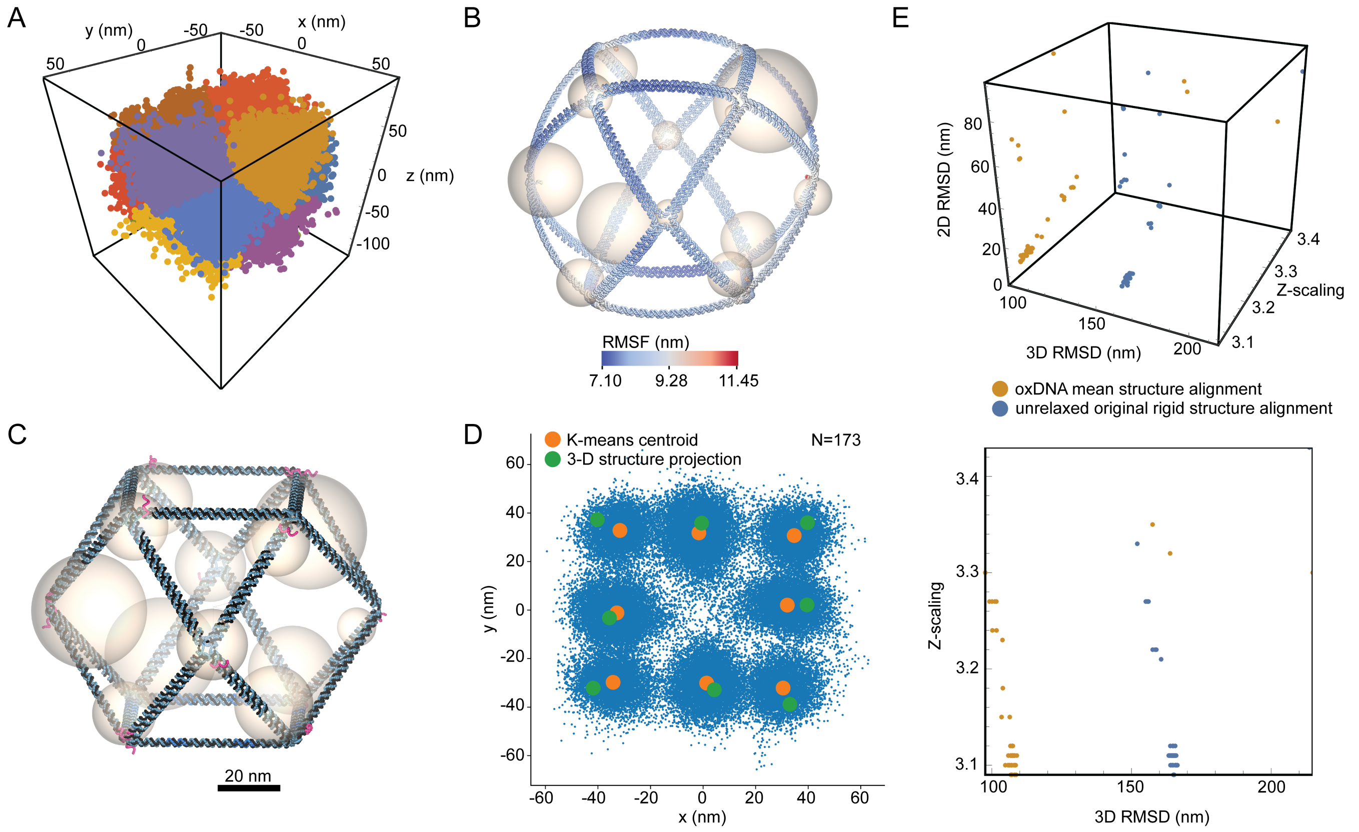

**Fig. S13** **3D clustering and alignment of DNA-PAINT experimental data and 3D cuboctahedron DNA origami structure.** (**A**) 3D k-means clustering of 3D DNA-PAINT localization data by assigning K=12. (**B**) 3D alignment of centroids from 3D k-means clustering result from **A** with the mean structure obtained from oxDNA simulation. The size of each sphere depicts the distance between the k-means centroid and the center of mass of the closest docking handle. (**C**) 3D alignment of centroids from 3D k-means clustering result from **A** with the unrelaxed structure. The size of each sphere depicts the distance between the k-means centroid and the center of mass of the closest docking handle. Scale bar in **B** and **C**: 20 nm. (**D**) An example of 2D alignment of the unrelaxed structure with docking handles' center of mass projected to x-y plane (the two stacking docking handles in $z$ direction are averaged). (**E**) (Top) Plot of 3D RMSD vs 2D projection RMSD after 3D alignment vs the Z-scaling of mean and unrelaxed structure. (Bottom) Plot of Z-scaling vs 3D RMSD showing the mean structure provides better RMSD, thus better alignment with the k-means centroid of experimental DNA-PAINT data compared to the unrelaxed structure alignment.

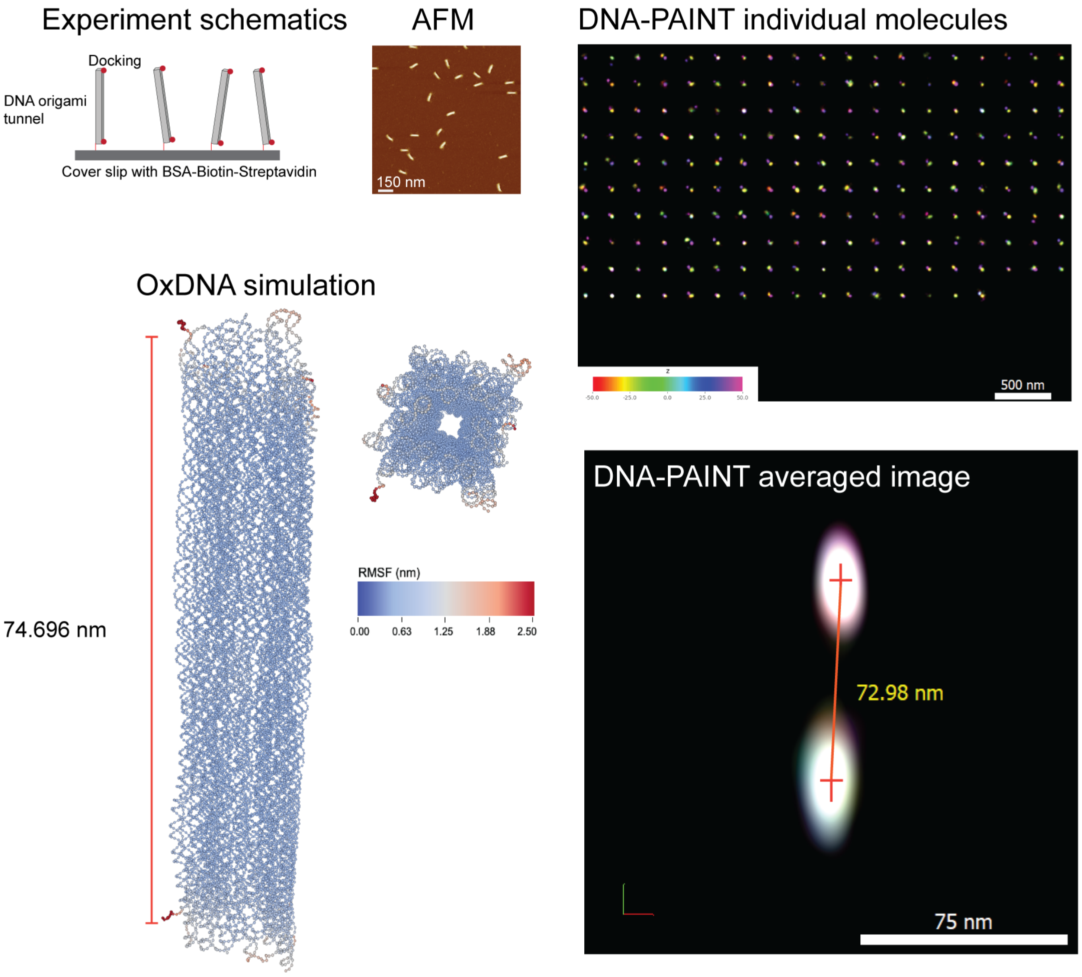

**Fig. S14** **3D tunnel DNA origami.** DNA-PAINT schematics of 3D tunnel DNA origami (Top left). AFM images of the origami (top middle), DNA-PAINT results of all picks and the averaged image showing distance of 72.98 nm (right panel) in agreement with the oxDNA simulation result (left panel)

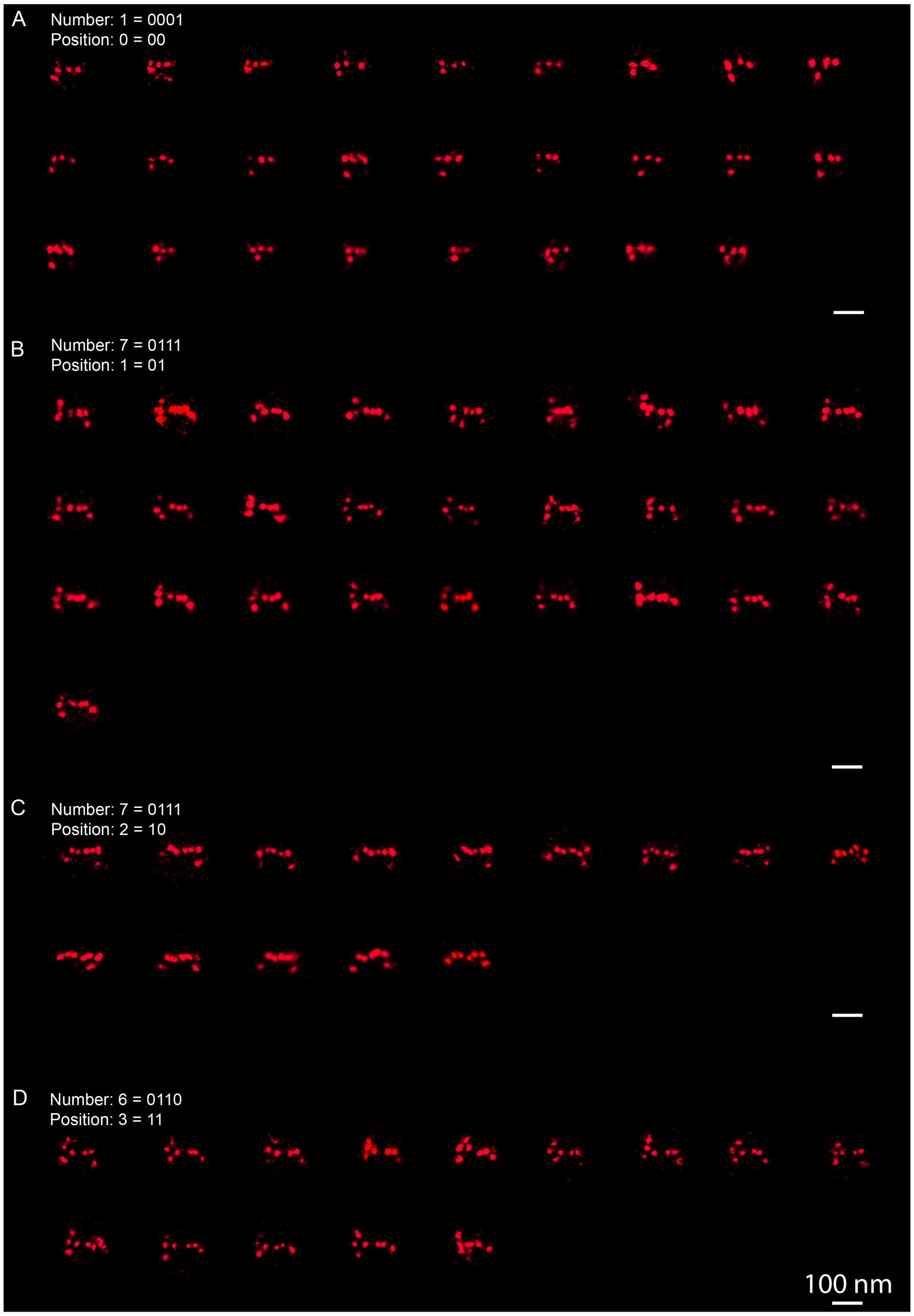

**Fig. S15 2D view of all picks analyzed in 3D DNA-PAINT "1776" dataset.** Full dataset of 2D projection of "1776" dataset. Scale bar: 100 nm.

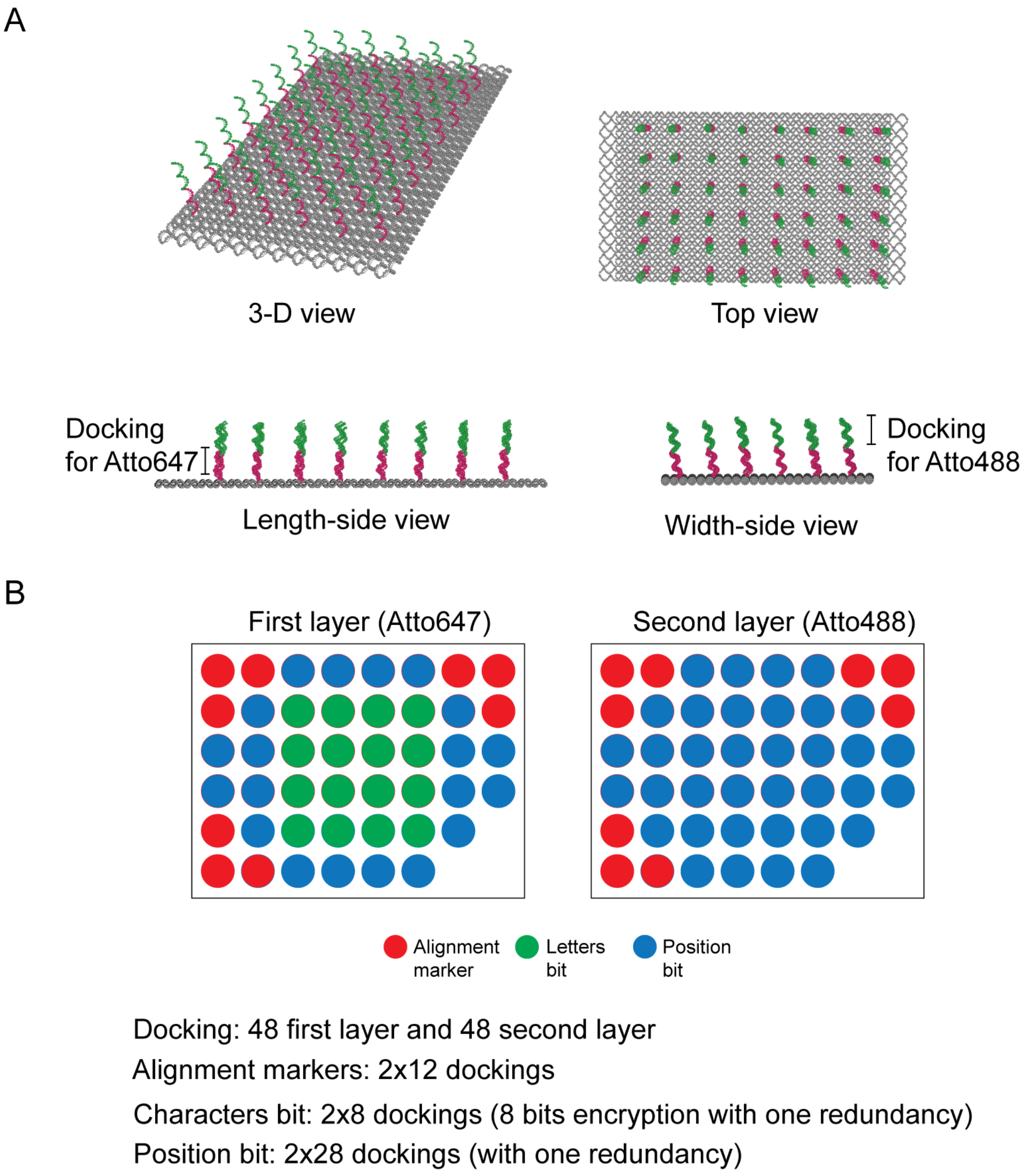

**Fig. S16 Design of two color 2D RRO encryption for high density information.** (**A**) The RRO schematic has two docking types for two different fluorophores. (**B**) The pattern encryption rules for two docking layers.

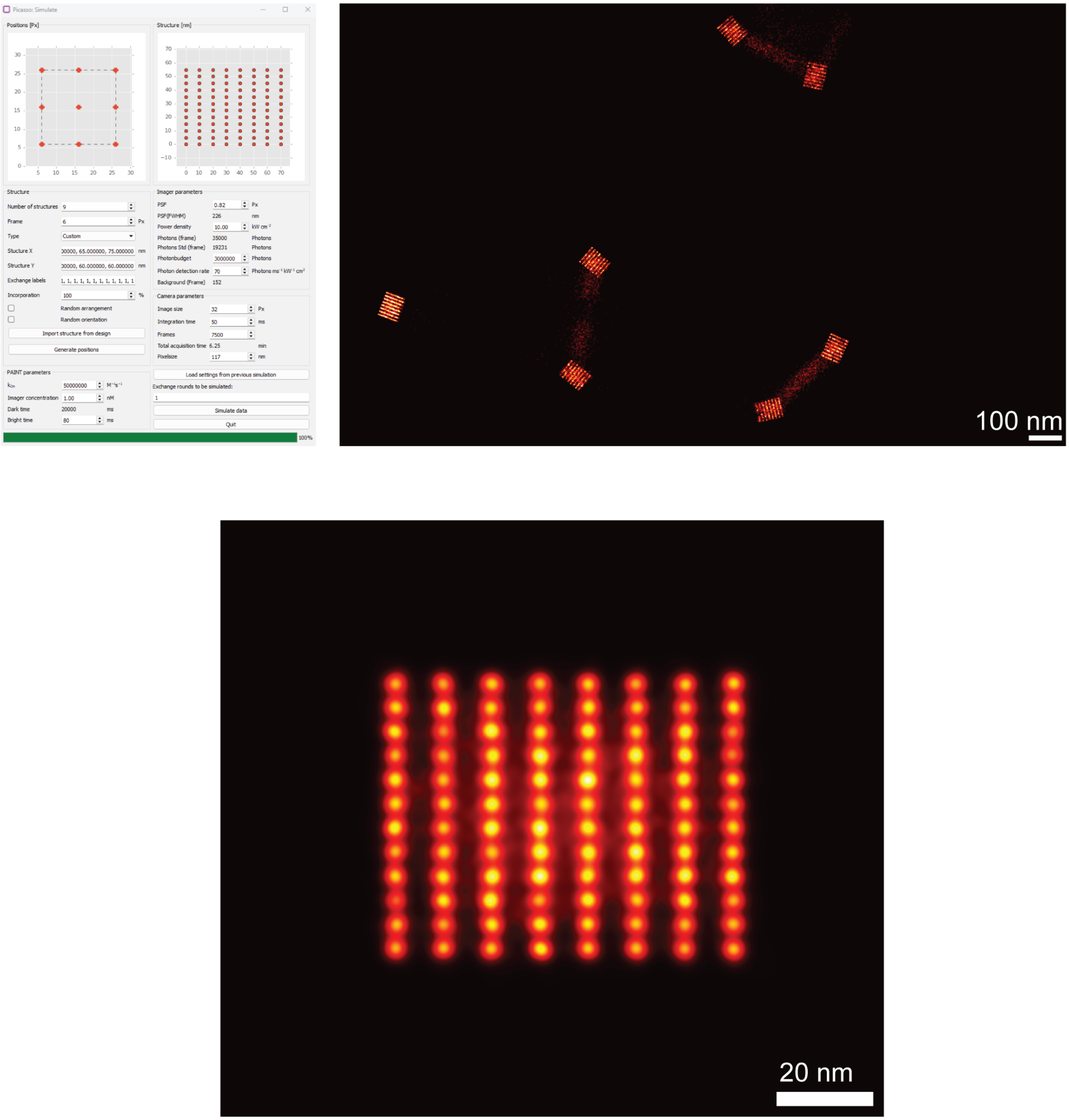

**Fig. S17 Simulated DNA-PAINT using the Picasso Simulate module** Illustration of data density when 4 bytes are used per origami. The averaged image over 9 structures shows clear 5 nm resolution in vertical direction and 10 nm resolution in horizontal direction (bottom).

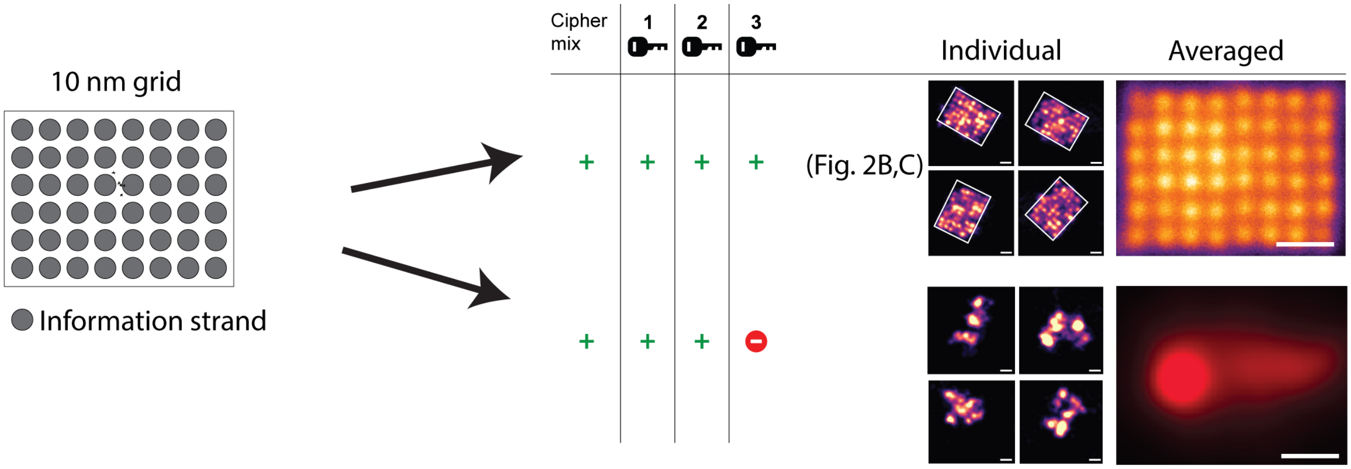

**Fig. S18 Incomplete staple incorporation prevents encrypted pattern formation.** The structure was assembeled using M13mp18 scaffold with docking strands for 10 nm grid (key 2) but other staples (key 3) were skipped. DNA-PAINT imaging and image reconstruction showed no detectable 10 nn grid pattern.

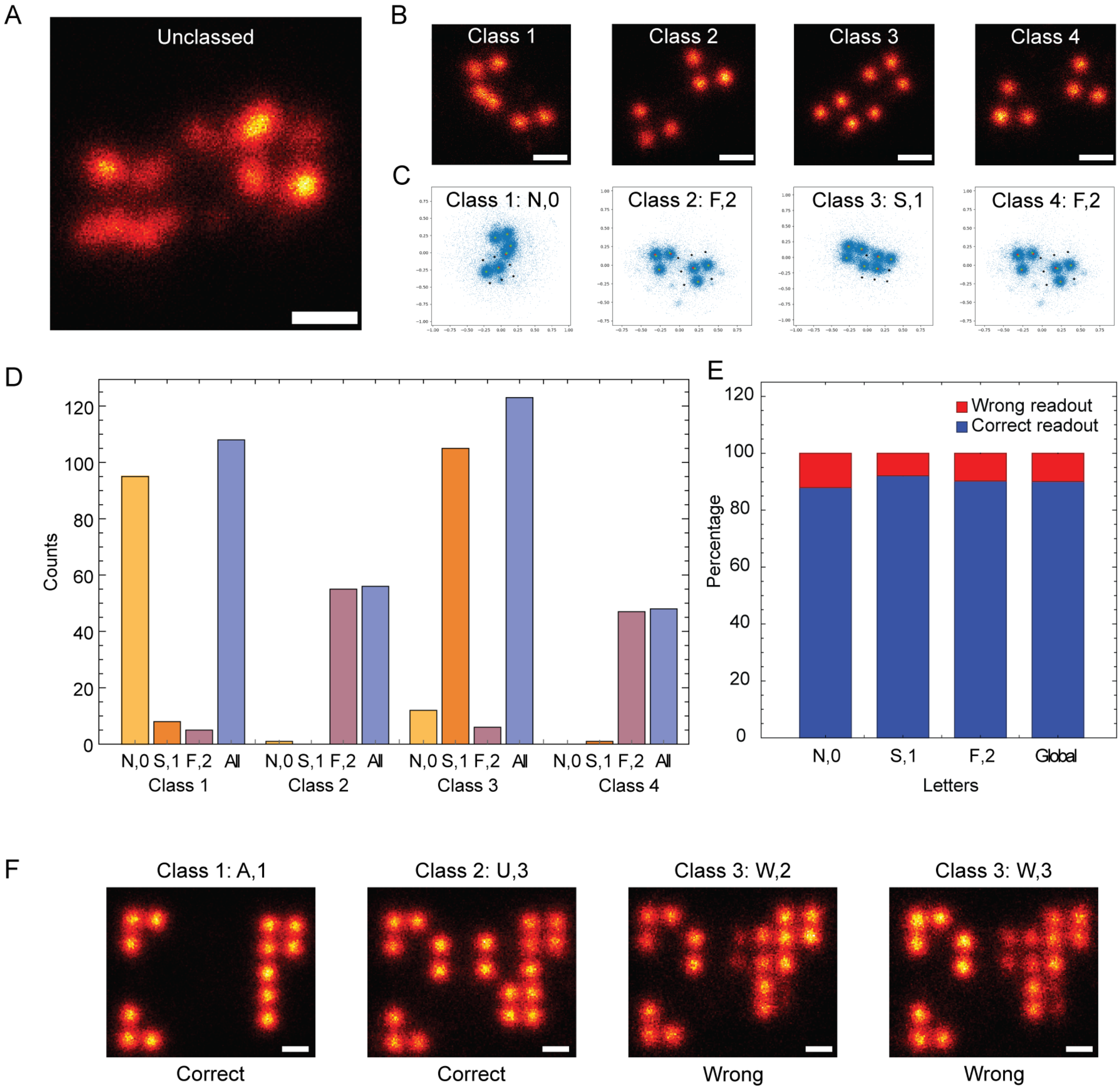

**Fig. S19 Unsupervised classification result using method described by Huijben et al. on "NSF" and "ASU" dataset.** (**A**) The superparticles of "NSF" before classification.(**B**) Superparticles of each class after classification showing 4 classes. (**C**) Readout of each class. (**D**) Classes' members show a few miss-classed patterns, thus not affecting the superparticles.(**E**) The readout accuracy of each class.(**F**) The classification of "ASU" is one redundancy dataset showing two correct classes and two wrong classes, thus making it impossible to recover "ASU". Scale bar: 20 nm in (A) and (B), 10 nm in (F).

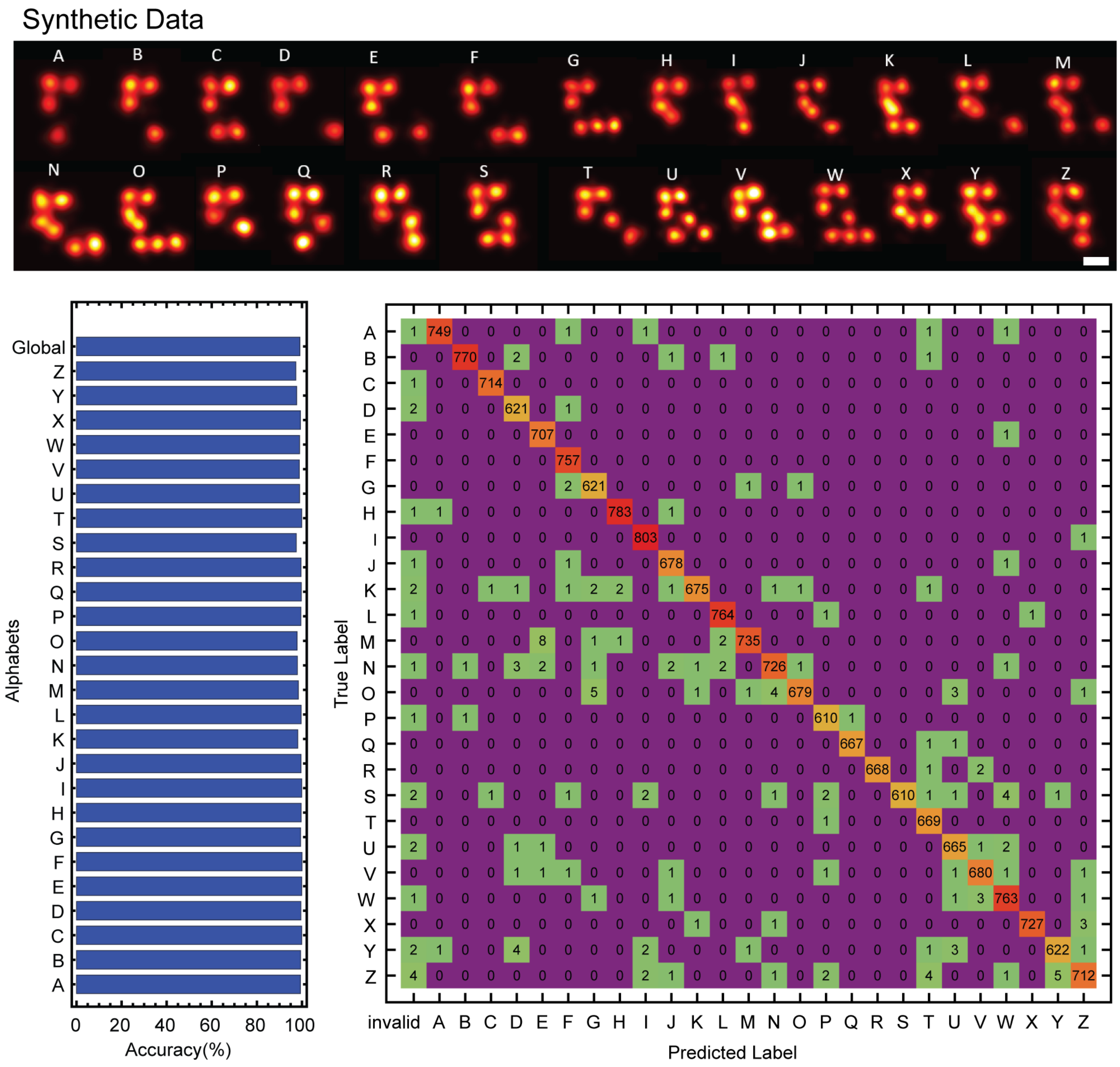

**Fig. S20 ResNet CNN results on synthetic data generated by Picasso Simulate module of letters A-Z without position encoding.** Examples of the 26 alphabets of synthetic data generated through Picasso Simulate module (top). The accuracy of each alphabet with ResNet-50 (bottom left) and the confusion matrix (bottom right).

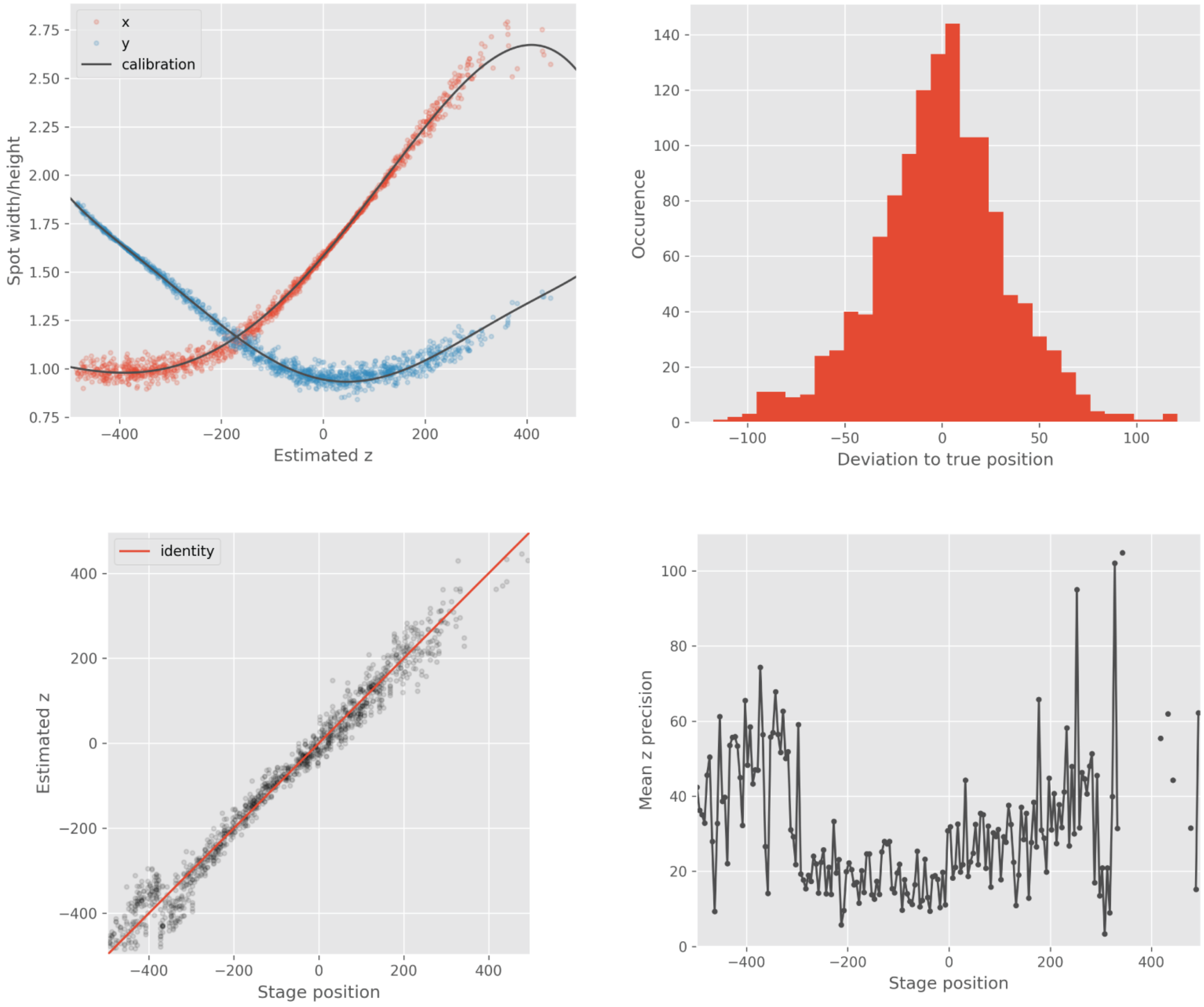

**Fig. S21** **3D calibration curve generated by the Localize feature of Picasso.**[65] (Top left) The localization spot widths and heights with the fit. (Top right) The distribution of the deviation from the true position. (Bottom left) The estimation of the $z$ coordinate as a function of stage position. (Bottom right) The mean $z$ precision is a function of stage position.
